# Dysregulation of pupal diapause in hybrid progenies, introgression and species delimitation within and beyond the Old World Swallowtail (*Papilio machaon* Linnaeus) butterfly complex

**DOI:** 10.64898/2026.06.30.735657

**Authors:** François Michel, Luc Legal, Krushnameg Kunte, Henri Descimon

## Abstract

In search of recurrent patterns of postzygotic hybrid incompatibility, we investigated the Holarctic ‘Old World Swallowtail’ (*Papilio machaon* Linnaeus) butterfly complex, whose members can easily be crossed in the laboratory. How many species this model system comprises remains unclear, as taxa with highly distinctive larvae but uncertain status come into contact with authentic *P. machaon* subspecies in Southern parts of the Palearctic region. By determining mitochondrial and ITS2 haplotypes within and away from contact zones, we found that these neighboring populations do exchange genes, as expected from F1 hybrids being generally fertile in our laboratory crosses. Nevertheless, recurrent instances of dysregulation of diapause were uncovered in hybrid progenies. In keeping with Haldane’s Rule, pupae of the heterogametic (female) sex were either unable to enter diapause or, in reciprocal crosses, unable to resume development after having initiated diapause, whereas F1 males experienced normal, photoperiod-regulated diapause, but passed on abnormal diapause regulation to part of their female offspring when backcrossed. Comparing male and female pupal weights in hybrid progenies provides estimates of parental incompatibility that allow to rank taxa and predict quantitatively the outcome of additional crosses, as expected if the same regulatory system were involved. A survey of the entomological literature confirms that diapausing pupae that cannot resume development (‘perpetual nymphs’) are a recurrent feature of interspecific lepidopteran crosses. Moreover, of the two parent species of a perpetual nymph, the paternal one generally has fewer broods per year. These observations are discussed in the light of models of the speciation process.

## INTRODUCTION

For two populations in contact to qualify as, or be able to evolve into, distinct species their ability to exchange genes needs to be severely limited by reproductive isolation. Among patterns of reproductive isolation, prezygotic barriers that prevent mating by behavioral/mechanical incompatibilities or ecological isolation are traditionally opposed to postzygotic incompatibilities, which include hybrid inviability or sterility. That distinction is an especially relevant one, for dysfunction/maladaptation of hybrids should lead to counter-selection of the propensity to generate them, resulting in increasingly efficient prezygotic isolation (Dobzhansky, 1940), whereas prezygotic incompatibility is not expected to provide grounds for promotion of hybrid inviability (at least in the absence of parental investment in the development of progeny (Coyne, 1974)); that is, except indirectly, through genetic coupling mechanisms (Servedio & Saetre, 2003); (Butlin & Smadja, 2018). This is the source of the assumption that preexisting postzygotic incompatibilies must have a decisive role in triggering the speciation process when previously isolated populations are brought into contact.

There is another, more trivial and somewhat unfortunate basis for contrasting postzygotic and prezygotic barriers. Mating preferences may in principle be investigated in the laboratory (reviewed by (Dougherty, 2020)), even though reasonable reconstitution of natural contexts can be challenging or just not feasible. However, in order to assess the viability and fertility of interspeific hybrids, it is necessary to obtain interspecific matings in the first place, something which, by essence, is not guaranteed. In this respect, Lepidoptera are noteworthy for providing unrivalled opportunities, as the incentive to generate novel, spectacular hybrid specimens has led to the development, first among amateurs and sellers and then by professionals, of various techniques that make it possible to overcome not only behavioral obstacles to interspecific copulation, but some mechanical ones as well (e.g. (Chovet, 1998)).

The most popular approach to lepidopteran provoked mating is ‘hand-pairing’ (Clarke & Sheppard, 1956a), in which stimulation and appropriate positioning of male and female abdomens triggers copulation (admittedly, with varying success depending on particular systematic groups). The opportunities offered by hand-pairing of butterflies were well illustrated by (Ae, 1979) who carried out extensive investigations of hybrid viability in the vast *Papilio* genus; some of the adult hybrids he was able to generate have now been revealed to have originated from pairs of species that diverged as many as 20 or so Myears ago (Condamine *et al*., 2023). Clarke and Sheppard themselves had undertaken to generate hybrids and even backcrosses within several *Papilio* subgroups, including *Papilio machaon* and his close relatives, but encountered serious breeding problems in the latter case (Clarke & Sheppard, 1953) (Clarke & Sheppard, 1955a) (Clarke & Sheppard, 1955b) (Clarke & Sheppard, 1956b).

*Papilio machaon*, the ‘Old World Swallowtail’, is an iconic insect – the largest resident butterfly in the British Isles – and a powerful flier, whose range extends over nearly the entire Palearctic region and parts of North America. It belongs to a set of closely related taxa that are notorious for having long defied taxonomists, to the point that the group has become a model system for the study of speciation. North American members of the *P. machaon* complex exemplify indeed the concept of ‘bad species’, that is, ‘taxonomic units that do not conform to criteria used to delimit species’ (Descimon & Mallet, 2009). At times when typology predominated, up to a dozen of the Nearctic relatives of *P. machaon* came to be regarded as species at some time or other (e.g. (Holland, 1930)). However, once genetics – first classical, then molecular – began to be taken into account, the number of accepted species sunk progressively down to five or so (e.g. (Scott, 1986); according to the somewhat extreme views of (Tyler *et al*., 1994), no more than three species – the number of major mitochondrial lineages (Sperling & Harrison, 1994) – should be retained).

Our current understanding of the *machaon* complex in the Americas rests on the genomic investigations of (Dupuis & Sperling, 2015) (Dupuis & Sperling, 2020), who showed that except for *P. indra* Reakirt, which diverged early from the rest of the group, these butterflies’ taxonomic misbehavior stems from massive, repeated exchanges of genes between members of an exclusively American mitochondrial lineage (*P. polyxenes* Fabricius, *P. zelicaon* Lucas) and various Nearctic offshoots of the predominantly Old World *P. machaon*. Three of these offshoots (*brevicauda* Saunders, *joanae* Heitzman, *kahli* Chermock and Chermock), which are commonly regarded as separate species because they inhabit widely separate geographic areas and exhibit distinctive wing patterns, were found to pertain to a particular *machaon* mitochondrial subclade and to share related, hybrid nuclear genomes. By contrast, the remaining, parapatrically distributed taxa (*aliaska* Scudder, *pikei* Sperling, *hudsonianus* Clark, *dodi* McDunnough, *oregonia* Edwards, *bairdii* Edwards), display gradual morphological and genetic differentiation along a mainly North to South axis, which makes it difficult to decide where to draw the line that would separate some of them from *P. machaon* proper.

The *machaon* complex has proved particularly attractive to geneticists as well, because of the ease with which interspecific crosses can be generated and the fact that hybrids, when viable – which is usually the case for males and often true also of females, generally proved at least partly fertile when backcrossed to their parent species (reviewed in (Ae, 1979)). These observations, which are consistent of course with the prevalence of introgression in the evolution of the group, raise questions about how these taxa maintain their integrity in natural settings. Part of the answer is ecological differentiation, as discussed at length by (Sperling, 2011). For example, the larvae of *pikei*, *dodi*, *oregonia* and *bairdii*, which are sympatric with *P. zelicaon*, use *Artemisia* (Asteraceae) as larval foodplant, whereas a majority of members of the *machaon* complex, including *zelicaon*, feed preferentially on Apiaceae. As for *joanae* adults, they were reported to be confined to forest clearings, which host their larval foodplants in the Ozark Mountains of Missouri and Arkansas, whereas sympatric *polyxenes* frequents preferentially meadows and roadsides (Heitzman, 1973). Allochrony may also prevent gene flow: on Cape Breton Island (Nova Scotia), where the ranges of *polyxenes* and *brevicauda* overlap, adults of the latter emerge in between the two flights of the former (Ferguson, 1953). Nevertheless, individuals with apparently hybrid phenotypes are not infrequent in nature. The most extreme case remains that of *P. zelicaon* and *P. machaon dodi*, which have been reported to generate ‘hybrid swarms’ in south-western Alberta, with up to 90 percent of individuals exhibiting hybrid phenotypes (Sperling, 1987) (as subsequently shown, however, by (Dupuis & Sperling, 2016), morphology and various molecular criteria result in different estimates of the frequency of hybrids and the time frame of hybridization remains uncertain).

The question of how genetic integrity is maintained in spite of enduring hybridization is relevant as well in Corsica and Sardinia, where *P. machaon* cohabits with *P. hospiton* Géné, an endemic species with markedly distinct larval (Figure 1) and, to some extent, adult (Figure S1) morphologies. (Aubert *et al*., 1997) investigated the situation in detail and their and some previous authors’ findings may be summarized as follows. Despite partial ecological separation, the ranges of *hospiton* and *machaon* overlap at low to moderate elevations and both sexes of the two species may be found together, especially close to hilltops, where males of *hospiton* spend most of the day waiting and looking for females. When placed in free-flying cages, males, whether of *machaon* or *hospiton*, are strongly attracted to females of the other species and spontaneous interspecific matings are easily obtained. In fact, individuals that look like lab-generated F1 hybrids – these are readily distinguished from their parents, both as larvae (Figure 1) and adults – are rather commonly encountered in nature; their frequency was estimated to reach up to *ca* 5% at some places in Corsica. Moreover, a majority of F1 hybrid individuals of both sexes were verified to be fertile in lab-carried backcrosses, so that all conditions appeared to be fulfilled for gene flow. Solid evidence of introgression was eventually obtained by (Cianchi *et al*., 2003) who determined allelic frequencies at allozyme loci that they had found to be diagnostic by comparing *P. hospiton* with continental specimens of *P. machaon*. While 147 individuals with a *hospiton* morphology turned out to be largely devoid of *machaon*-specific alleles, high levels (up to 43%) of introgression of *hospiton* alleles into 281 Corsican and Sardinian *machaon* individuals were uncovered at several autosomal loci. Additionally, (Cassar *et al*., 2025) recently reported having detected introgression from *hospiton* to *machaon* in genomic DNA sequences. Unfortunately, their sample – three *machaon* individuals from the same Corsican locality and a single *hospiton* – is far too small for any definite quantitative conclusions to be drawn.

**FIGURE 1.**
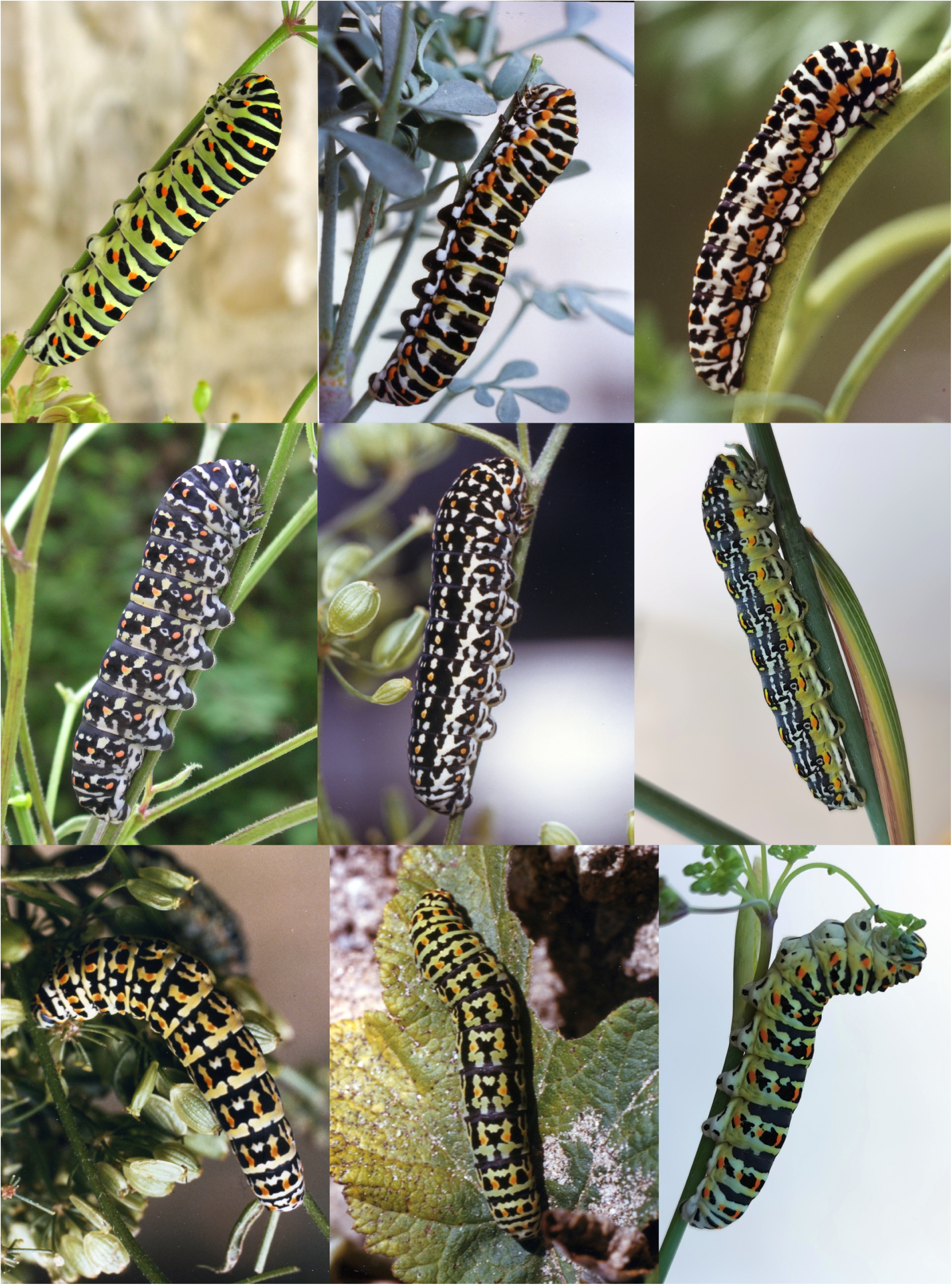
Mature (L5) larvae of Old World members of the *P. machaon* complex and some hybrids. First row, left: *P. machaon* (France, Rochefort-en-Yvelines); middle: *P. machaon mauretanica* (Morocco, *ca* 100 km South-West of Marrakech); right: *P. machaon saharae* (Morocco, Tizi n’Tiniffift). Second row, left: *P. ladakensi*s (India, Ladakh, Leh); middle: *P. everesti* (China, Tibet, Nyalam); right: *P. hospiton* (Corsica, Novella). Third row, left: lab-generated F1 hybrid with a *P. everesti* father and a *P. machaon* mother; middle: presumed hybrid between *P. everesti* and *P. machaon* photographed at Nyalam (China, Tibet) on Aug. 6, 1987; right: lab-generated F1 hybrid with a *P. hospiton* father and a *P. machaon* mother.

Assuming that gene flow from *hospiton* into *machaon* is an ongoing process (this remains an open question, see discussions in ((Aubert *et al*., 1997); (Cianchi *et al*., 2003); (Cassar *et al*., 2025)), a clue to what might limit it was provided by the finding that regulation of diapause is perturbed in hybrid progenies (Aubert *et al*., 1997). In male hybrids, pupal diapause is induced by larval exposure to long nights and development resumes after prolonged exposure to cold conditions, just as in their *machaon* parent. In female hybrids, however, two opposite outcomes were observed, depending on which of the two species the mother belonged to (recall that females are the heterogametic sex in Lepidoptera). Hybrid female pupae with *hospiton* mothers were wholly unable to enter diapause, whatever the breeding conditions, and would develop directly to the adult stage. By contrast, females resulting from the reciprocal cross (*machaon* mother x *hospiton* father) were unable to resume development after having entered diapause: their fate was to become ‘perpetual nymphs’, unless diapause had been prevented by rearing larvae under both short nights and elevated temperatures.

That interspecific crosses involving members of the *machaon* complex other than *hospiton* could result in delayed development of females or, in some cases, in the complete absence of adult females from hybrid progenies was well known (e.g. (Clarke & Sheppard, 1956b); (Ae, 1964); (Oliver, 1969). On the other hand, complete inability to diapause had not previously been reported (at least explicitly so), possibly because of poorly controlled or insufficiently diverse rearing conditions. That only the heterogametic sex should be affected by hybrid malfunction is in accordance of course with Haldane’s Rule (Haldane, 1922) and reflects some form of incompatibility between the autosomes of one parent species and, in Lepidoptera female hybrids, the single Z chromosome inherited from the other parent. Still, having reciprocal crosses affected in the same function, although in seemingly inverse ways, was somewhat unusual and clearly deserved being further investigated.

Corsica and Sardinia are the only places in the Palearctic region where two undoubtedly distinct species of the *machaon* complex coexist extensively. Nonetheless several Old World ‘subspecies’ of the widely distributed *P. machaon* (Figure 2) have recurrently been suspected to correspond in fact to different species. One of these is *hippocrates* C. & R. Felder from Japan and Sakhalin, whose summer-generation females have wings much obscured (Figure S1). Three other candidates for species status are noteworthy for having larvae (Figure 1) that depart strikingly from the *P. machaon* pattern – the latter is shared, with minor variations, by all other members of the complex except *P. hospiton*. Among them, *saharae* Oberthür, from desert edges in North Africa and the Arabic peninsula, has been bred and crossed with other members of the *machaon* complex a number of times ((Clarke & Sheppard, 1956b); (Clarke & Larsen, 1986); (Pierron, 1990)). By contrast, it is only a few years ago that pictures of the mature larvae of *everesti* Riley (previously known as *sikkimensis* Moore, see (Nazari *et al*., 2023)) from East Tibet and *ladakensis* Moore from the Western Himalayas became available ((Igarashi & Harada, 2015); (Hervillard *et al*., 2018)) and crosses involving these taxa had not been reported so far.

**FIGURE 2.**
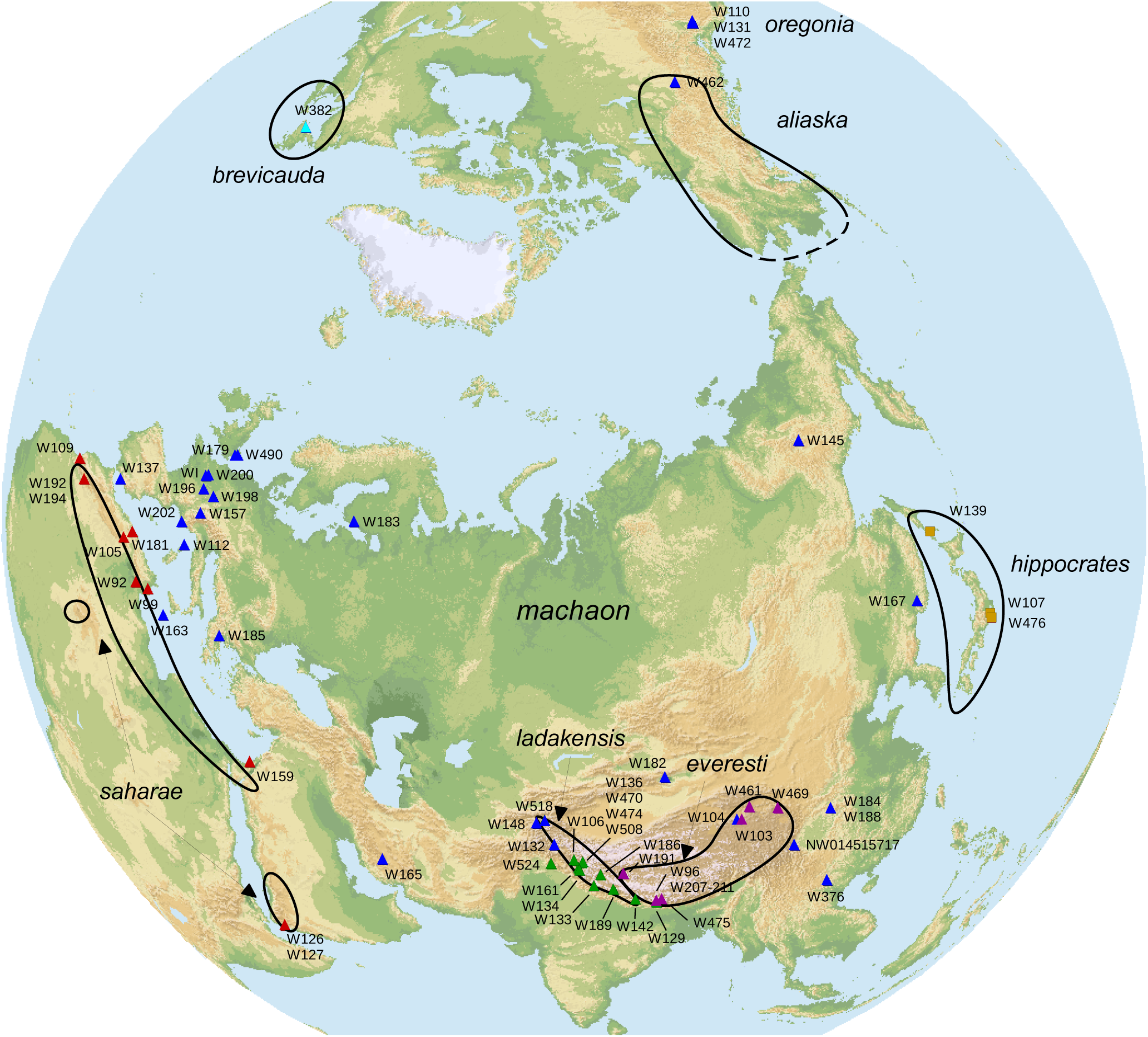
Geographic distribution of members of the *P. machaon* complex and origin of sampled individuals. Colors correspond to major groups based on mitochondrial phylogeny (Figure 3 and Table 1). Black curves indicate the approximate distribution areas of two North American taxa and major Old World entities other than *machaon* proper.

**Table 1.**
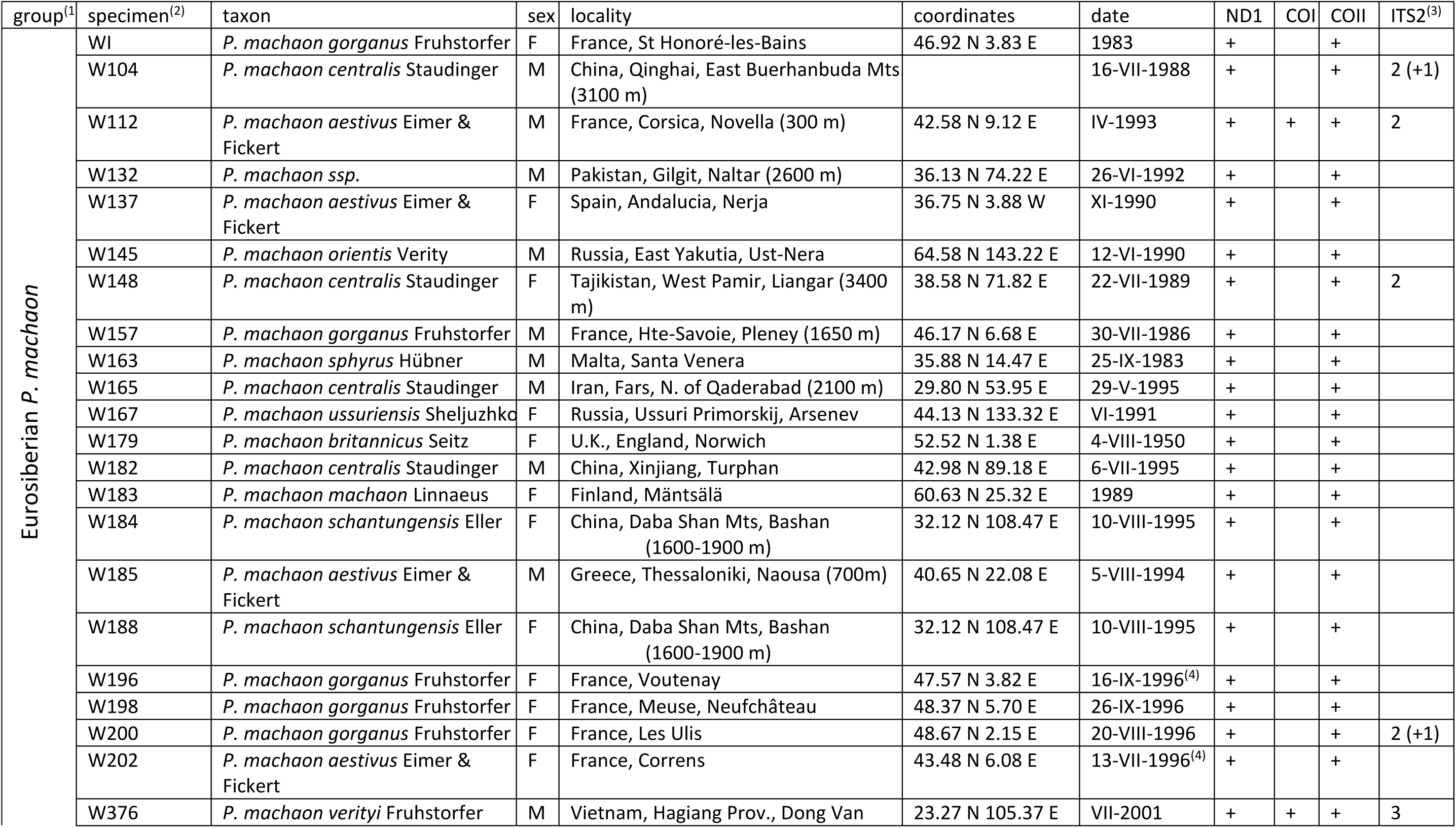
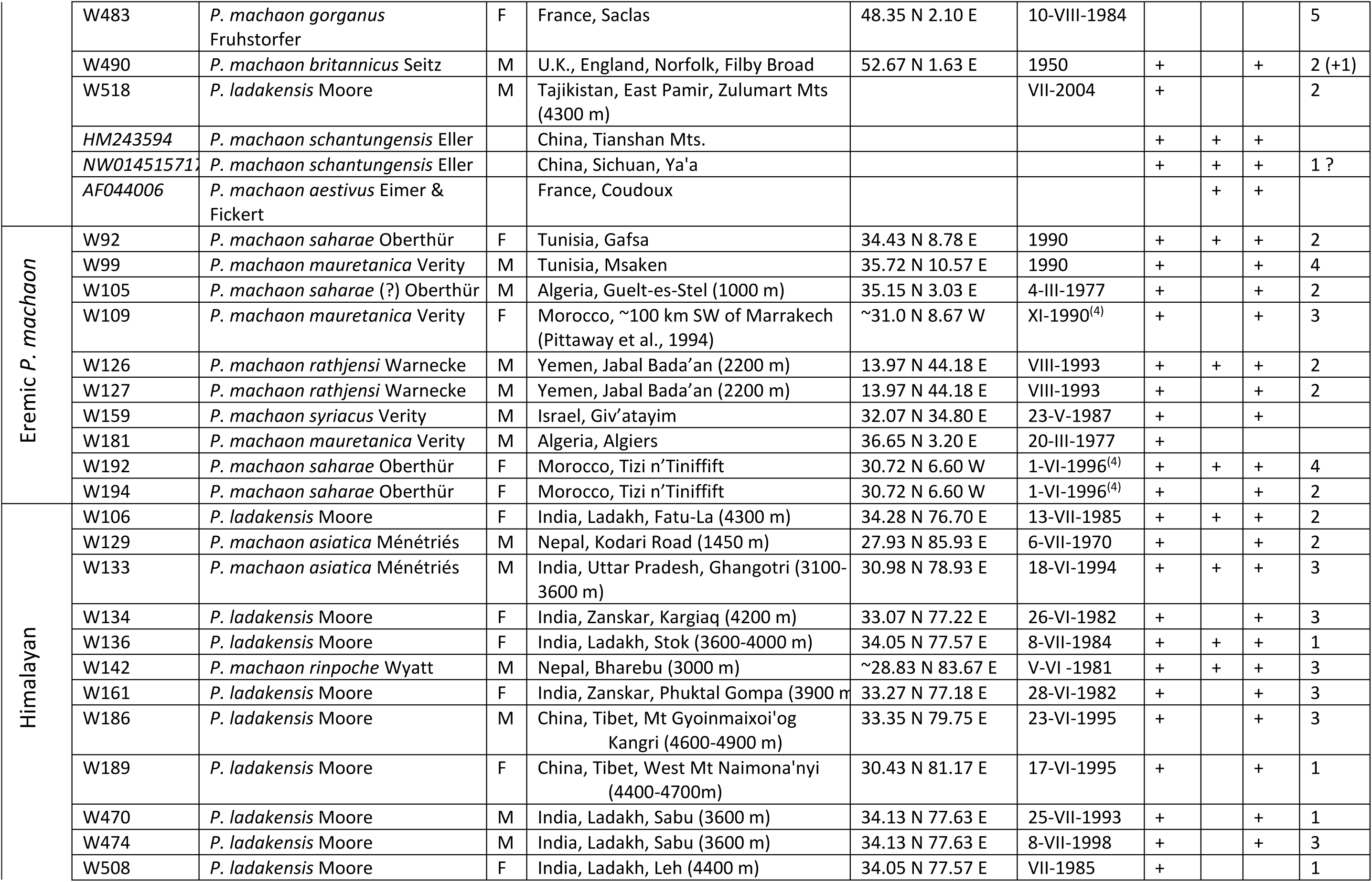
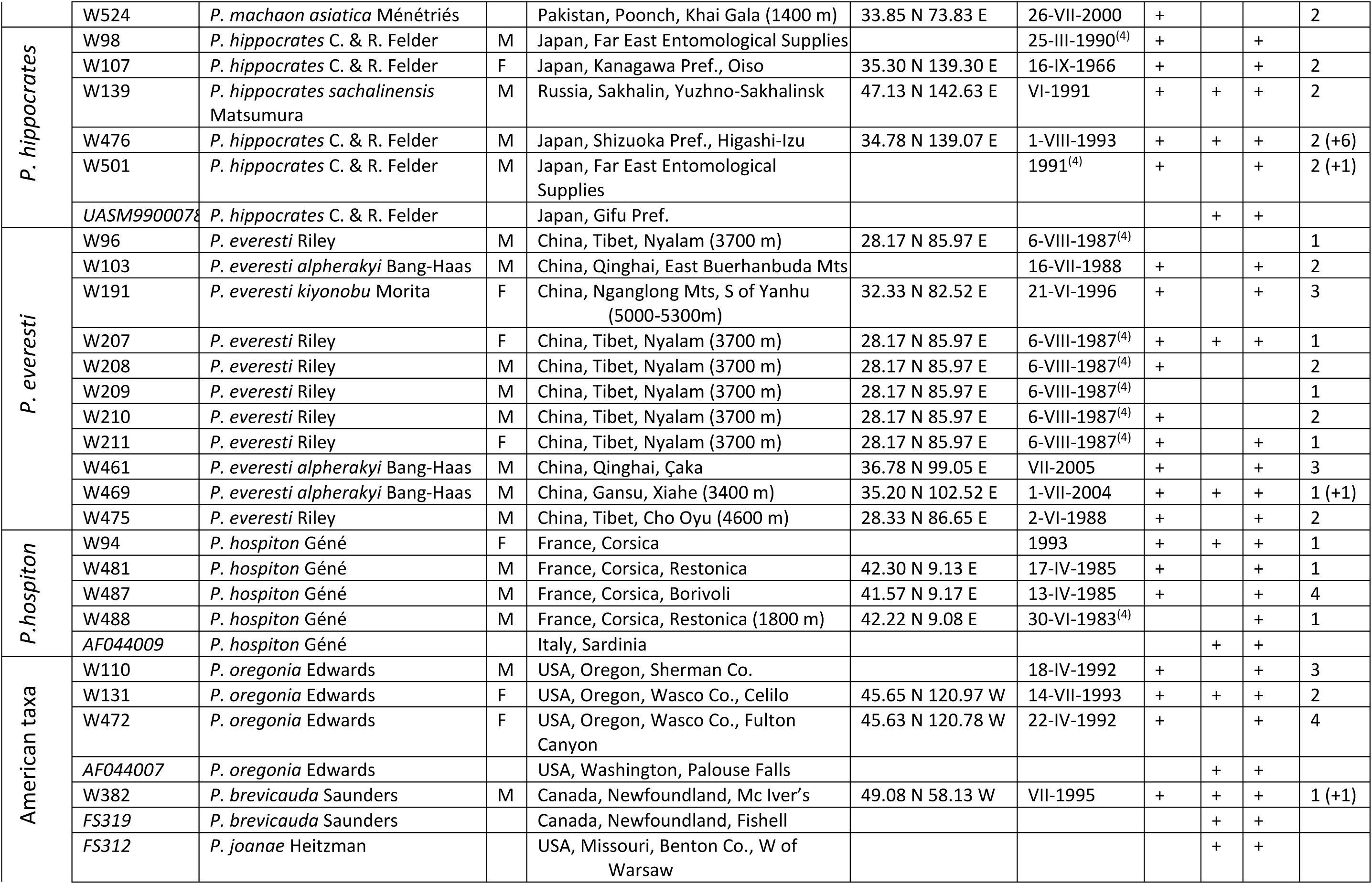
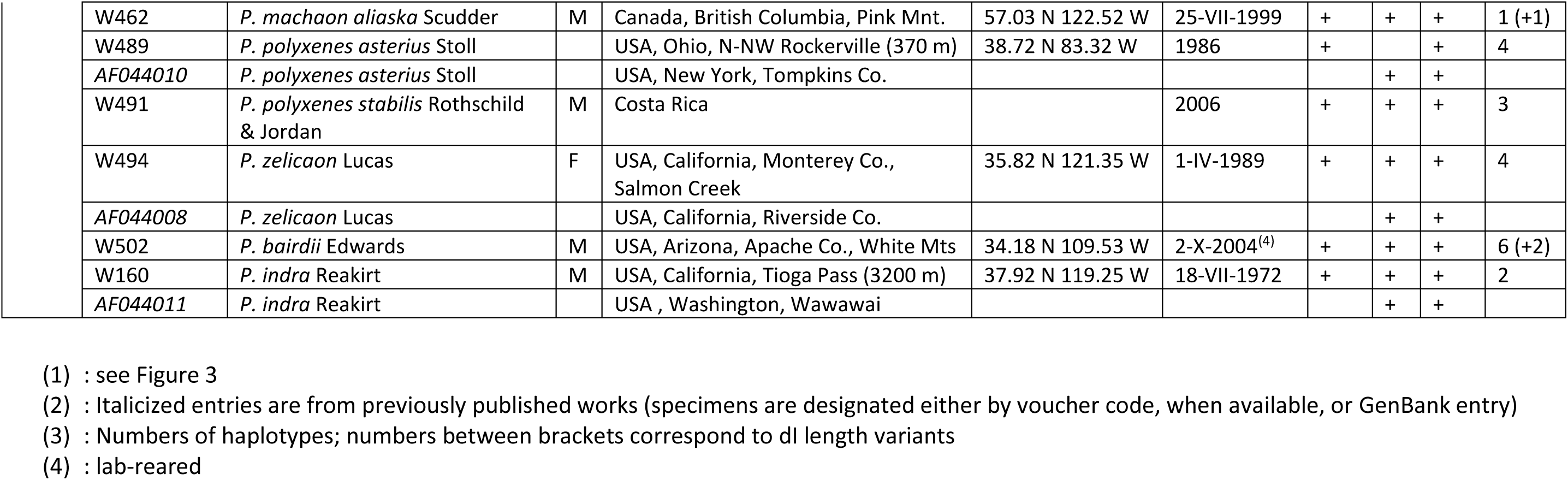
Specimens and sequences.

We undertook to determine mitochondrial and nuclear ITS2 haplotypes for sets of individuals belonging to the above-mentioned taxa of uncertain status and samples of *P. machaon* specimens collected within and away from the zones of contact. Our aim was to determine the phylogenetic relationships of candidate species within the *P. machaon* complex and to find out whether gene exchanges with their *machaon* neighbors could be detected. We chose to use the ITS2 segment of ribosomal DNA (rDNA) repeats as a source of nuclear haplotypes because its sequence evolves rapidly, yet it is easily and reliably amplified by PCR from highly conserved flanking sequences. Moreover, despite concerted evolution of rDNA units (Dover, 1982), individual repeats may retain considerably more intraspecific and intragenomic variations than is usually assumed (reviewed in (Wang *et al*., 2023)), as we found to be the case indeed for our own dataset.

In parallel to molecular investigations, laboratory crosses were carried out in order to estimate the viability and fertility of hybrids. We now report that not only introgression, but also defective regulation of pupal diapause in female hybrids is far from uncommon in the Palearctic section of the *P. machaon* complex. Moreover, when taxa are ranked according to the direction and extent of diapause dysregulation in F1 hybrid progenies, a correlation becomes apparent with the propensity to multivoltinism. This observation, which appears to hold in several families of Lepidoptera in which interspecific crosses were reported to result in ‘perpetual nymphs’, raises intriguing questions about its evolutionary significance and possible underlying molecular mechanisms.

## MATERIALS AND METHODS

### Specific and subspecific allocation of specimens

The Palearctic section of the *machaon* complex (excluding *hospiton*, but including *everesti*, *ladakensis*, *hippocrates* and *saharae*) has been divided into an unreasonably large number of ‘subspecies’ based primarily on rather minor variations in adult wing pattern and close to 40 of these names appear to be still in use (Domagala and Lis, 2022). Fortunately, a recent review of the group (Nazari *et al*., 2023) brought this number down to 25 and we chose to follow this revised nomenclature, except for specimens W132 (unclassified), W159 (‘*syriacus*’ Verity) and W142 (‘*rinpoche*’ Wyatt; this individual collected at 3000 m looks strikingly different from *asiatica* Ménétriés specimens flying at lower elevations). Nevertheles, subspecific names in Table 1 should be understood to have been provided for convenience only, most of them were inferred merely from the geographic origin of specimens and it has not been our intention to take part in any nomenclatural issue. Based on larval (Figure 1) and/or adult (Figure S1) morphology, only *saharae*, *ladakensis*, *everesti* and *hippocrates* are sufficiently distinct in fact from *machaon* to undoubtedly deserve being set apart and the last three of them are actually deemed to be separate species in the present work. Although the larva of typical *saharae* is quite different from that of *machaon* (Figure 1), adults are notoriously difficult to distinguish. In the absence of any information about larval morphology and foodplants at Guelt-es-Stel (Algeria), specimen W105 (Table 1) was nevertheless concluded to belong possibly to *saharae* based on its small size, reduced hindwing anal red spot and 31 antennal segments (Pierron, 1990).

Sampling and breeding of *P. hospiton*, a protected species, was made possible by a permit from the French Ministry of the Environment to H.D. (Aubert *et al*., 1997). Phylogenetic analysis of mitochondrial sequences

The three DNA segments that were targeted for analysis are part of the ND1, COX1 and COX2 genes, their lengths are 471, 664 and 617 nt (totaling 1752 nt) and their sequences were determined for 74, 21 and 69 individuals, respectively (Table 1; see Extended Methods for DNA extraction, PCR amplification and sequencing). Alignment of mitochondrial sequences was straightforward in the absence of indels.

Phylogenetic analysis of mitochondrial data was carried out with MrBayes 3.2.7 (Ronquist *et al*., 2012) and involved the 75 specimens for which we had obtained sequences together with 11 other specimens from GenBank for which DNA sequences were available over at least two of the three DNA segments used in this work (Table 1). Data were partitioned by codon position (1 together with 2, separately from 3), while the three DNA segments were pooled. Analysis assumed a GTR (general time-reversible) model of nucleotide substitutions and a fraction of invariable sites, while variable sites were distributed over four categories of evolutionary rates. Parameters were estimated separately for each of the two partitions and a strict clock model was used. For time calibration, the divergence of *P. indra* from the rest of the *P. machaon* group was assumed to have occurred 10.8 My ago (Condamine *et al*., 2012).

### Generation and phylogenetic analysis of ITS2 sequences

The ITS2 DNA segment was amplified and sequenced from 59 extracts (Table 1). In only 12 cases did amplified DNA yield a unique sequence. For all other samples, a complex pattern resulting from the superimposition of several sequences over part of the length of the DNA segment of interest was observed. In the most commonly encountered situation (22 cases), in which just two haplotypes differing by one or more indels were involved, individual sequences could generally be inferred by comparing reads from the two ends of the PCR product. However, more complex cases required additional PCR and sequencing runs with haplotype-specific primers (Table S1), the 3’ end of which was chosen to coincide with the first or first two positions at which haplotype sequences had been observed to diverge (see Table S1; we did not attempt to identify component sequences that were estimated from chromatograms to constitute less than 10 to 15 percent of the total). In this way, presumably genuine haplotype sequences with unambiguous combinations of indels and substitutions could most often be obtained; only for highly complex mixtures (e.g. W476, W502; Table 1) did the exact number and sequence of haplotypes remain tentative. DNA was reextracted from several individuals that had yielded particularly complex patterns in order to ascertain that those patterns were genuine, rather than reflecting DNA contamination. It was also verified that the 33 nt deletion in stem III of the P. machaon rathjensi H127B haplotype was consistently recovered from separate PCR runs (judging from sequencing chromatograms, this haplotype is about as abundant as H127A; however, see Discussion).

Out of a total of 98 aligned sequences from a subset of 44 Palearctic individuals of the *P. machaon* complex (other than *P. hospiton*) to which were added the closely related *P. machaon aliaska* W462 and *P. brevicauda* W382, we recovered no fewer than 71 different haplotypes (Figure 4 and Table 1; see Extended Methods for the alignment procedure). Among these 71 haplotypes, 57 were observed only once, and the most frequent haplotype was found in no more than 5 individuals (Figure 4); for this reason, we deemed it more convenient to keep designating haplotypes by the coding number of the individual from which they originated, followed by letters and/or numbers if necessary, rather than renaming them according to sequence identity.

**FIGURE 3.**
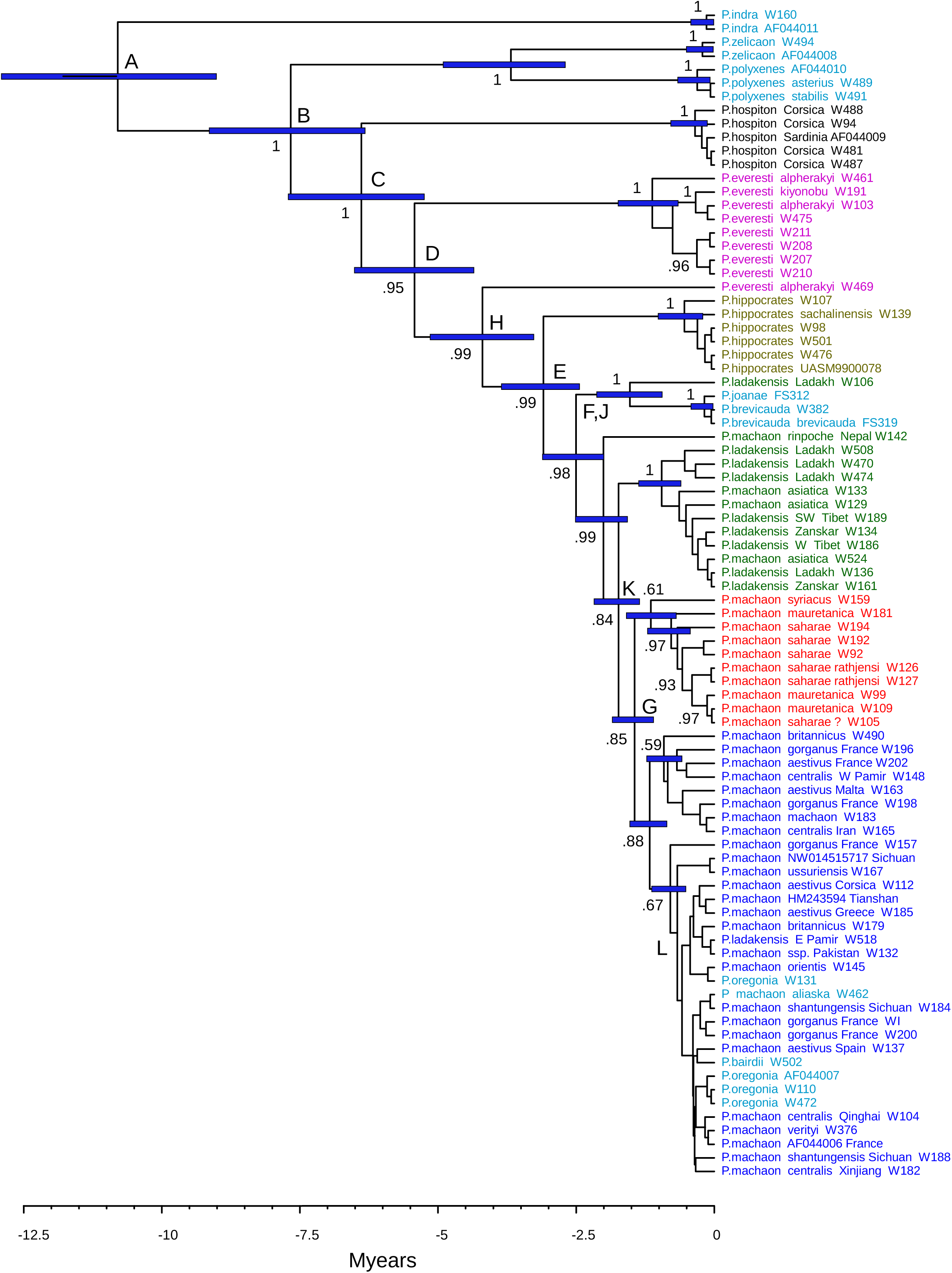
Phylogenetic tree of the *P. machaon* complex based on mitochondrial DNA sequences. Numbers next to nodes are Bayesian estimates of clade probabilities, blue bars indicate 95% credibility intervals for branch lengths (see Materials and Methods for tree-building procedure). Time calibration was achieved by assuming the deepest node, i.e. divergence of *indra* from the rest of the complex, to be 10.8 Myears (million years) old, see Text. Colors correspond to major phylogenetic (this tree) and/or geographic and morphologic entities (see Text and Figures 1,2): eurosiberian *P. machaon* (deep blue), eremic *P. machaon saharae, P.m. mauretanica, P.m. rathjensi* (red), Himalayan *P. machaon* and *P. ladakensis* (green), *P. hippocrates* (dull yellow), *P. everesti* (magenta), *P. hospiton* (black). North American specimens are shown in light blue.

**FIGURE 4.**
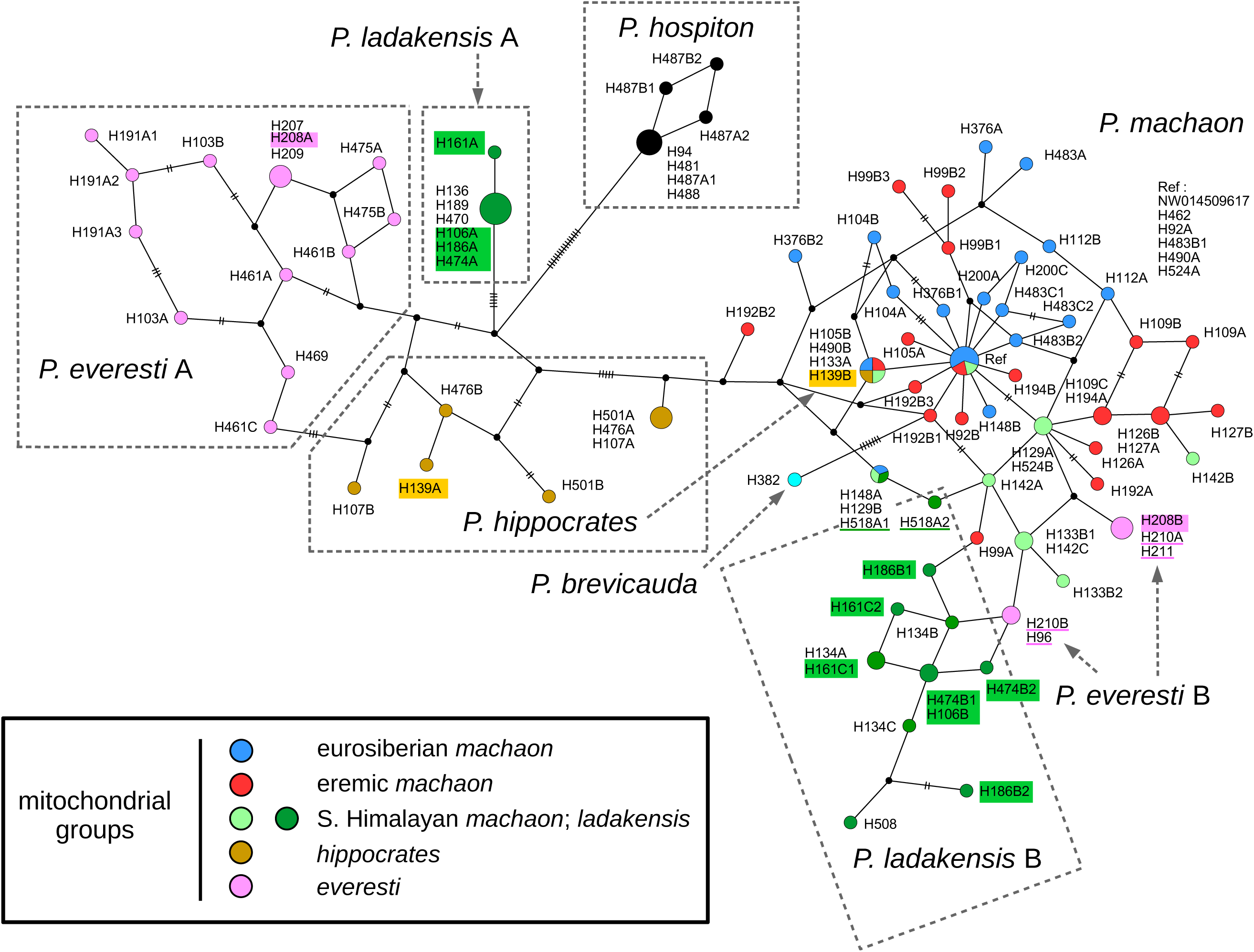
TCS network of ITS2 haplotypes in Old World members of the *P. machaon* complex, together with *P. machaon aliaska* and *P. brevicauda*. Small black dots correspond to inferred, unobserved network nodes. The surface of observed nodes is proportional to the number of haplotypes of identical sequence included (‘Ref’ stands for the reference NW014509617 sequence and five identical haplotypes). Observed nodes are colored according to the individual(s) the haplotypes come from (see Figures 2 and 3 for color coding of specimens; in the few cases of identical haplotypes existing in individuals that belong to different mitochondrial groupings, nodes with multiple colors result). Haplotypes that belong to distinct ITS2 groups, as defined in this Figure, yet coexist in the same individual, are highlighted. Small bars across lines connecting nodes indicate numbers of substitutions, when more than one. The length of node-connecting lines was set proportional to the square root of associated numbers of substitutions, except in the densest areas of the machaon sub-network, which were slightly blown up.

We used statistical parsimony (Posada & Crandall, 2001) and the TCS algorithm, as implemented in the PopART-1.7 software (Leigh & Bryant, 2015), to build a haplotype network from this dataset together with the *P. hospiton* sequences. However, since PopART does not distinguish nucleotide deletions from missing data, we resorted to the following recoding procedure in order to avoid losing a considerable fraction of phylogenetic information. For each indel, the first ‘missing’ nucleotide was replaced by a base absent in all the other sequences at that site (there always existed at least one), while the remaining positions were left missing; as a result, indels were counted as single mutational events, irrespective of their lengths. A second alignment was separately generated and recoded for seven American specimens (2 *polyxenes*, one *zelicaon*, 3 *oregonia*, 1 *bairdii*) with numerous, closely related haplotypes differing by a total of 8 substitutions and 12 indels. The corresponding TCS network is shown in Figure S4.

For more divergent haplotypes, evolution was assumed to have been largely clonal and a tree-building approach using MrBayes 3.2.7 was selected. A reduced dataset including 56 variable sites out of 387 indel-free alignable positions was partitioned into paired and unpaired nucleotides (stems and loops), to which the ‘doublet’ and GTR models of base substitution, respectively, were applied. Otherwise, analysis was run as for mitochondrial sequences, with a strict clock model enforced. Two sets of 2000 trees were generated, the first 500 trees in each set were discarded and a consensus tree was calculated. Graphical superimposition of individual trees (Figure 5a) was achieved with DensiTree version 2.7.7.

**FIGURE 5.**
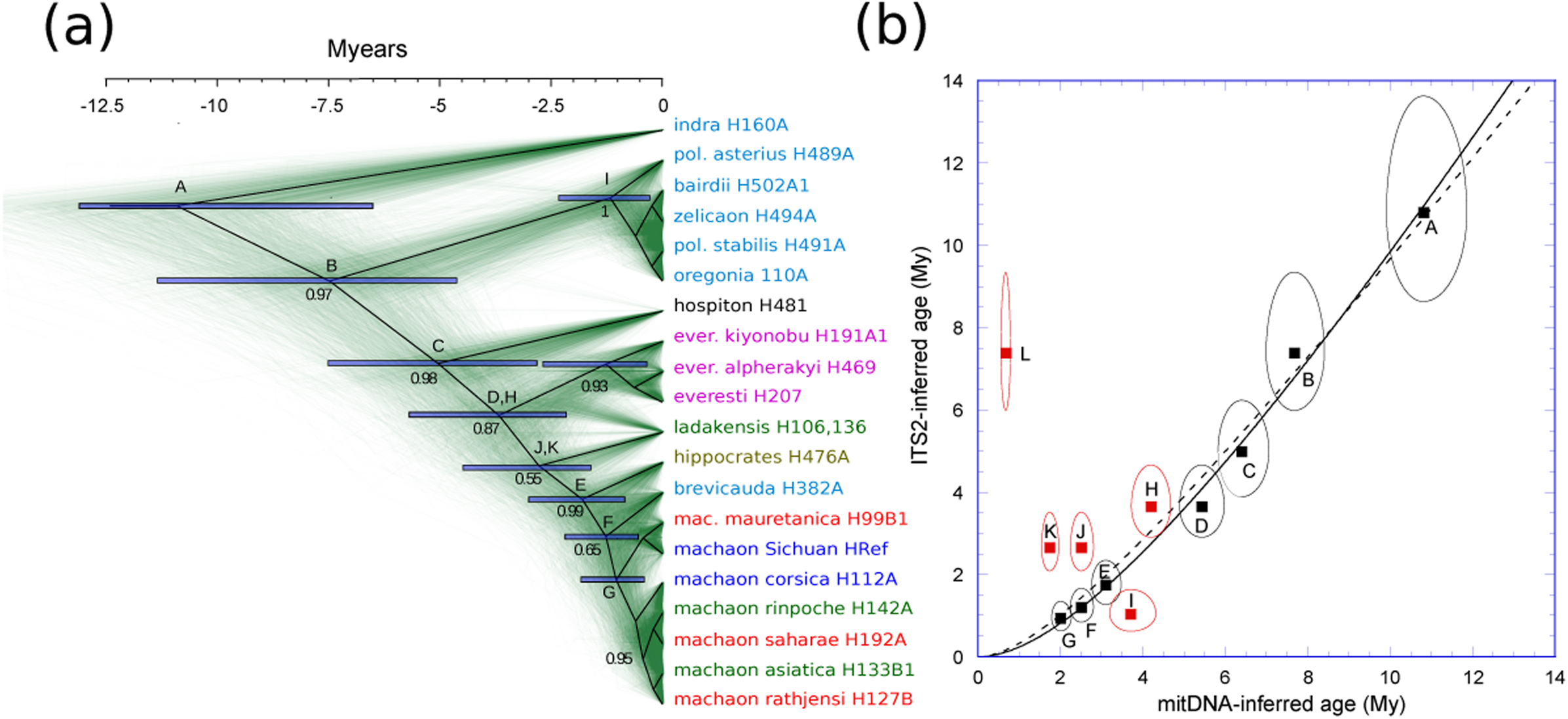
Phylogenetic tree of the *P. machaon* complex based on ITS2 sequences and comparison of mitochondrial-estimated and ITS2-estimated rates and times of divergence. A subset of ITS2 sequences chosen to represent the molecular diversity of major taxa on the one hand, and *P. machaon sensu stricto* on the other, was selected from the network in Figure 4. (a) ITS2 phylogeny. Black lines: consensus tree generated by MrBayes 3.2.7, with a strict clock model, from a set of 3000 trees (Materials and Methods). Superimposition of individual trees using DensiTree v2.2.7. is shown in green. Numbers next to nodes, blue bars, time calibration and colors of taxa as in Figure 3. (b) ITS2- versus mtDNA-estimated times of divergence. Data are from Figure 3 (abscissa) and panel (a) (ordinates). Axes of ellipses correspond to 68% credibility intervals (1σ). Presumed mito-nuclear concordance is indicated by black squares, whereas red squares illustrate definite or likely instances of mito-nuclear discordance. (D), divergence of *everesti* (except W469) from Old World *machaon* and allies; (H), divergence of *everesti* W469; (J), divergence of *ladakensis* W106; (K), divergence of ‘*ladakensis* A’ other than W106; (I), divergence of *polyxenes asterius* from *zelicaon*; (L), divergence of *bairdii* and *oregonia* W110 from reference *machaon* sequence. Full curve (*y* = 1.780 (x – ln (1 + 0.322 x)/0.322)) – see Materials and Methods – fits points A to G (in black; Pearson’s R = 0.996); dashed curve (y = 1.395 (x – ln (1 + 0.663 x)/0.663)) attempts to fit points A to K (R = 0.952).

### Modelling ITS2 rates of divergence

When the divergence of mitochondrial sequences is compared to that of ITS2 sequences, the latter first appears to lag behind the former and then catches up (Figure 5b). In an attempt to account for this observation we have assumed that divergence of ITS2 haplotypes is initially slowed down by the efficiency of recombination between the many copies of the ribosomal RNA genes. However, as sequences and their insect hosts begin to diverge from one another, both the frequency of genetic exchanges between incipient species and the efficiency of recombination between different haplotype sequences should progressively decrease, with a concomitant increase in the rate of divergence. In the simplest conceivable model we could come up with, the rate of divergence of mitochondrial sequences (dx/dt) is constant over time, whereas that of ITS2 sequences (dy/dt) is obtained by subtraction of a term that is inversely proportional to a linear function of the time elapsed since divergence:

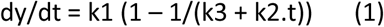

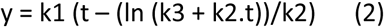

k3 is expected to be approximately equal to 1 (as assumed indeed in Figure 5b), since the diversity of ITS2 haplotypes should be close to 0 when t = 0. The fact is, that when only nodes A to G of Figure 5b are retained (see Results), fitting the data with equation (2) (with k1, k2 and k3 as adjustable parameters) yields k3 = 0.975, with divergence at zero time estimated to be 6.1 10^-4^ per site. For this particular ITS2 dataset, in which five out of 20 individuals were found to retain haplotype diversity after removal of all indel-corresponding sites, the actual initial diversity, averaged first over all haplotypes associated with any individual and then over all individuals, amounts to 5.8 10^-4^ substitution per site.

### Modelling the response to the photoperiodic signal

In order to propose an explanation for the correlation between the magnitude and direction of diapause dysregulation in female hybrids on the one hand, and voltinism of their male parent on the other, we have modelled photoperiodic control of diapause as follows. The critical parameter is assumed to be the relative concentrations of two substances: substance ‘Z’ is under control of one or several loci on chromosome Z, substance ‘A’ depends on a number of autosomal loci, and the [z]/[a] concentration ratio governs the length of the L5 larval feeding stage, pupal weight and pupal diapause (with a threshold at [z]/[a] = 1; see Discussion). Photoperiodic control is ensured by having a fraction of substance Z, 1 – [z]/z_T_, sequestered by equilibrium binding to a molecule P whose (limiting) total concentration p_T_ is proportional to daily exposure to light (here p_T_ = H/12, with H varying from 0 to 24 h; i.e. we chose an ‘hourglass timer’ (Veerman, 2001)). Thus,

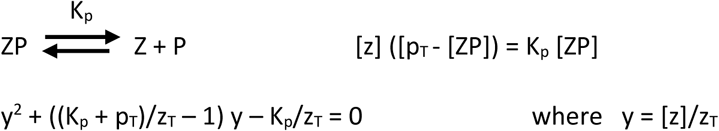

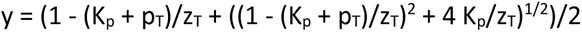

Curves describing the photoperiodic response in Figure 9 correspond to various combinations of [a] and z_T_ (in arbitrary units) with K_p_ = 0.5. In F1 hybrids, values of [a] and z_T_ are averages of parental values, except for the single Z chromosome of females whose associated z_T_ is the paternal one.

## RESULTS

### Mitochondrial phylogeny

Collection sites of the 74 specimens from which we obtained mitochondrial DNA sequences are shown in Figure 2 and listed in Table 1 (GenBank sequences from 11 additional individuals were included in our analyses). 64 individuals were from Old World locations and based on collection sites and adult morphology (Materials and Methods), 11 of them could be referred to *everesti*, 10 to *ladakensis*, 5 to 6 to *saharae* and 5 to *hippocrates*. Among other specimens, 11 came from places located next, or rather close to the distribution areas of several of these taxa: three were from the southern slope of the Himalayas, two from northern Pakistan, one from Pamir, one from Tibet, one from Israel and three from North Africa.

As was to be expected, the well-supported overall topology of our mitochondrial-based phylogeny (Figure 3) is generally compatible to a number of previously published ones with fewer, less diverse taxa ((Aubert *et al*., 1999); (Condamine *et al*., 2012); (Dupuis & Sperling, 2015); (Nazari *et al*., 2024)). The first event was the separation of the lineage leading to *P. indra* from the rest of the group, and this was followed by the divergence of the ancestor of exclusively American species *P. polyxenes* and *P. zelicaon* from predominantly Old World lineages. Within the Palearctic realm, the deepest node separates geographically restricted *P. hospiton* from the vastly distributed *P. machaon* and its close allies. Among the latter, East Tibetan *everesti* was the first entity to diverge, some 5.5 Myears ago. Interestingly, this taxon has retained significant mitochondrial diversity despite its restricted distribution, for two separate, paraphyletic mitochondrial lineages were found to coexist within the set of nine individuals that we examined. Even though the wing patterns of specimen W469 are quite similar to those of individuals attributed to ssp. *alpherakyi* by (Nazari *et al*., 2023), its mtDNA differs markedly from other available *everesti* mtDNAs, with divergence from W207 mtDNA over the COX1 ‘barcoding’ segment amounting to 3.5 percent (23 substitutions out of 663 nt). The next node sets the six very similar *P. hippocrates* sequences apart from *P. machaon*. Next come two partly intermingled sequence subsets. One of them corresponds to a mitochondrial lineage previously identified by (Dupuis & Sperling, 2015), whose prominent members are three related North American taxa of hybrid origin, *brevicauda*, *joanae* and *kahli*. The other subset groups together the Himalayan sequences, thus failing to distinguish individuals with a *ladakensis* wing pattern, which come from the dry inner valleys to the North of the main Himalayan range, from dwellers of the South slope, which experience temperate to subtropical climates.

The remaining part of the phylogenetic tree, which spans the last 1.5 Myears, is rather poorly resolved, except for a well-supported monophyletic subset that groups together not only *saharae* (three to four specimens from North Africa; Materials and Methods) and *rathjensi* (two specimens from Yemen), which are known to share similarly patterned larvae ((Clarke & Larsen, 1986); (Pittaway *et al*., 1994)), but also three additional North African individuals collected within the distribution area of *mauretanica*, which has a *machaon*-like larva (for W99 – from larvae collected on fennel, a *machaon*, not *saharae*, larval foodplant – and W109, see (Pierron, 1990) and (Pittaway *et al*., 1994), respectively; as for individual W181, it was collected next to Algiers; note also that an additional specimen, W159, from central Israel may belong to that clade). The other noteworthy point, which was already emphasized by (Dupuis & Sperling, 2015), is the remarkable genetic homogeneity of mitochondrial DNA in the rest of the *machaon* clade, not only throughout Eurasia (to the exclusion of the Himalayas, North Africa and Yemen), but over parts of Western North America as well. Even though sequencing did reveal a fair amount of local diversity – the sequences of three individuals collected within no more than *ca* 250 km of each other in Northern France were found to differ from one another by 9 to 11 substitutions (out of 1088 aligned sites) – variation appears largely disconnected from geographic distance. For example, sequence W137 from Southern Spain differs from sequence W462 (from Alaska) by a mere 2 substitutions (out of 1085 aligned sites) and from W502 (*P. bairdii*, from Arizona, at a distance of more than 15 000 km by land and Behring Strait from Spain) by only two additional ones.

### Old World ITS2 haplotypes

Concerted evolution of ribosomal DNA repeats ((Eickbush & Eickbush, 2007) and references therein) requires particularly efficient recombination mechanisms, which should result in highly reticulate evolution that only networks can adequately picture. The TCS network shown in Figure 4 was generated from an alignment of 98 ‘Old World’ ITS2 sequences (71 distinct haplotypes) from *P. machaon* and allies (one *P. brevicauda*, one *P. machaon aliaska* and seven *P. hospiton* sequences were added). The sequences in that alignment had been recoded so that not only base substitutions, but also indels might be taken into account (Materials and Methods; out of a total of 526 aligned sites, 56 were found variable and 37 parsimony-informative). One striking feature of this network is the remarkably high fraction of haplotypes that were observed only once (57 out of 71, excluding *P. hospiton*). Moreover, while 9 samples yielded but a single haplotype, considerable diversity was found among the other 35 individuals, with an average of 2.5 detectable haplotypes per specimen. Still, this is certain to be a far underestimate of actual diversity, first because variation at the tip of helix I was discarded, and also since only major components of PCR amplification mixtures were sequenced, see Materials and Methods.

In Figure 4, not only ITS2 haplotypes from *P. hospiton*, but a majority of those from individuals belonging to *P. everesti* on the one hand and *P. hippocrates* on the other are seen to form separate sub-networks. By contrast, haplotypes from Eurasiatic and North African *P. machaon* specimens are merged into a single, rather compact, poorly structured, highly connected cluster. More unexpectedly, the *ladakensis* haplotypes were found to fall into two quite distant sub-networks. One of these (*ladakensis* A), with two haplotypes and seven sequences, is directly connected to the central, ‘root’ node that links together the *hospiton*, *everesti* and *hippocrates* sub-networks. The other one (*ladakensis* B), with a somewhat larger number of haplotypes and sequences, lies a few steps away from the main *machaon* cluster. Strikingly, a majority of the nodes connecting the latter two networks to one another correspond to haplotypes obtained from individuals (W129, W133, W142, W524 and four *everesti* specimens) collected South, rather than North, of the main Himalayan range. The two distinct *ladakensis* ITS2 molecular subsets actually coexist within a large part of the *ladakensis* range, as illustrated by the fact that four individuals (W106 and W474 from Ladakh, W161 from Zanskar and W186 from Tibet) were found to be ‘heterozygotes’ that possessed both type A and B sequences.

Two additional, clear-cut examples of individuals with mixed ITS2 ancestry are visible in Figure 4. One of them is provided by *everesti* specimen 208, from which both a typical Tibetan *everesti* (208A) sequence and a typically Himalayan (208B) sequence were obtained. Four other individuals from the same population, located at the very southern edge of the *everesti* range, yielded type B sequences and the remaining two, type A sequences. All these individuals had typical everesti wing patterns and possessed *everesti* mitochondrial DNA (Figure 3), so that the presence of Himalayan-like ITS2 sequences in some of them is a clear case of introgression from neighboring Nepalese *machaon* populations. The other case of suspected hybridization involves specimen W139 from Sakhalin (just North of Japan), from which both hippocrates-like and Eurasiatic machaon-like ITS2 sequences were obtained.

### Overall ITS2 phylogeny of the *P. machaon* complex

As populations and species differentiate, they become increasingly unlikely to exchange DNA. Moreover, population- and species-specific ITS2 haplotypes also diverge with time, so that shared segments of identical sequence become smaller, which will result in decreased rates of recombination. These two processes are expected to combine and make ITS2 evolution progressively more clonal with time elapsed since initial divergence. In the ITS2 phylogenetic tree of the *machaon* complex shown in Figure 5a, emphasis was put on deeper nodes and only a representative sample of more closely related haplotypes was retained. Moreover, indels were discarded from analyses, as poorly conserved segments cannot be reliably aligned.

Comparison of mitochondrial-based (Figure 3) and ITS2-based (Figure 5a) trees reveals both considerable similarity and clear cases of complete incongruence. With *P. indra* in basal position, the deepest node in both trees separates exclusively American species *polyxenes* and *zelicaon* from predominantly Old World taxa and among the latter, *hospiton* was the first one to diverge, followed by *everesti*. On the other hand, the ITS2 haplotypes of *oregonia* and *bairdii*, whose mitochondrial DNA was found to be essentially indistinguishable from that of geographically far distant Eurasiatic *machaon*, are closely related to those of their *polyxenes* and *zelicaon* neighbors (the TCS network in Figure S4, which makes visible connections between all ITS2 haplotypes of American origin, to the exclusion of *machon aliaska* H462 and *brevicauda* H432, shows that there actually exist shared haplotypes between *oregonia* and *bairdii* on the one hand, and *oregonia* and *zelicaon* on the other).

In order to assess more precisely the degree of congruence of the phylogenetic trees in Figures 3 and 5a, we compared sequence dissimilarity and estimated times of divergence between subsets of taxa at twelve nodes of our mitochondrial-derived and ITS2-derived phylogenetic trees (Materials and Methods and Figure 5b). Attempts to fit all nodes but L (divergence of *machaon* reference sequence from *bairdii* and *oregonia*) reveal several instances of significant discrepancy. Dates of divergence estimated from mitochondrial DNA are smaller than those derived from ITS2 comparisons for individuals harboring *ladakensis* ‘type A’ ITS2 haplotypes (nodes J and K). Conversely, mitochondrial DNA indicates an older date of divergence of *zelicaon* from *polyxenes* than ITS2 haplotypes (*polyxenes* actually appears polyphyletic in Figure 5a).

### Viability and fertility of hybrid individuals

Interspecific matings between members of the *machaon* complex other than *P. indra* are generally fertile ((Ae, 1979) and references therein; for *machaon* x *hospiton* crosses, a detailed assessment was provided by (Aubert *et al*., 1997)). Unsurprisingly, this is true as well of crosses involving *ladakensis* or *everesti*. Out of 12 hand-pairings of *everesti* (8 males, 4 females) with either *machaon* or *hospiton* (six individuals of each), 10 proved mostly fertile, i.e. a majority of eggs hatched. For *ladakensis*, 9 crosses were attempted that involved 6 males and 3 females with 8 *machaon* and 1 *hospiton* individuals and out of this total 7 were fertile. These ratios do not differ significantly from rates of success for intraspecific hand-pairing, which we estimate to be around 80 percent under optimal conditions (i.e., when three-day old males and one-day old females are used). Moreover, provided due care was exercised, most caterpillars developed all the way to the pupal stage, with no significant deviations from a 50:50 sex ratio.

For male hybrids resulting from crosses between *ladakensis* on the one hand, and either *machaon* or *hospiton* on the other, diapause was found to be facultative: larvae raised under long nights gave rise to diapausing pupae, which resumed development after overwintering, whereas pupae formed by larvae raised under short nights developed directly (the same is true of male hybrids between *machaon* and *hospiton* (Aubert *et al*., 1997)).

The fertility of male hybrids was assessed by backcrossing them with *machaon* females. Only one of the fourteen *machaon* x *ladakensis* males tested proved sterile, the others being highly fertile since in each case more than 90 % of eggs underwent development, most of those eggs hatched, and more than half of the resulting larvae made it to the pupal stage, as typically observed in fact with healthy broods from intraspecific crosses. By contrast, the fertility of hybrids between *machaon* and *everesti* appears to be significantly lower, as only two of the five males tested were fertile. Similarly, only two out of seven male hybrids between *hospiton* and *everesti* proved fertile when crossed to *machaon* females.

Data pertaining to the viability and fertility of hybrid females with one *P. machao*n parent are summarized in Table 2. For *machaon* x *hospiton* hybrids, our data confirm and extend previous findings (Aubert *et al*., 1997). The female progeny of a cross between a *hospiton* male and a *machaon* female consists almost entirely of perpetual nymphs, unless larvae were reared under both short-nights conditions and elevated temperatures. Conversely, hybrid female pupae with a *machaon* father and *hospiton* mother were found to be unable to enter diapause, even when larvae were reared under long-nights conditions and at temperatures that did not exceed 17°C (one experiment was performed outdoors in the second half of September in the neighborhood of Paris).

**Table 2.** Viability and fertility of hybrid progeny.

| Taxon 2 | F1 crosses (female offspring) |  | F2 (F1 x F1) offspring |
| --- | --- | --- | --- |
|  | *machaon* male x taxon 2 female | taxon 2 male x *machaon* female |  |
| *hippocrates* | viable, fertile | viable, fertile | – 1/1 cross fertile<sup>(1)</sup><br>– viability rather low |
| *saharae* | n.t. | – 1 cross<br>– all 25 female pupae developed directly (short-nights rearing)<br>– single crossed female fertile | – 1/1 cross fertile<sup>(1)</sup><br>– viability rather low (35 larvae and 11 pupae/200 eggs) |
| *ladakensis* | – 2 crosses<br>– all 19 female pupae viable, but unable to enter diapause after long-nights rearing.<br>– 2/3 backcrossed females fertile | – 4 crosses<br>– 53/54 female pupae **perpetual nymphs**.<br>– Single hatched female fertile | – 1/1 cross fertile<br>– viability intermediate (48 pupae/195 eggs) |
| *hospiton*<sup>(2)</sup> | – 5 crosses<br>– all female pupae viable, but unable to enter diapause after long-nights rearing.<br>– 2/2 backcrossed females fertile | – 14 crosses<br>– 58/59 diapausing female pupae **perpetual nymphs** (long-nights rearing).<br>– female pupae viable (short-night rearing)<br>– 8/13 backcrossed females fertile | – 1/20 crosses fertile<sup>(3)</sup><br>– viability intermediate (57 pupae/144 eggs) |
| *oregonia* | n. t. | – 1 cross<br>– all 12 female pupae **perpetual nymphs**. | n. t. |
| *everesti* | – 1 cross<br>– 7 viable female pupae<br>– 2/5 backcrossed females laid minute eggs<br>– 1/90 eggs hatched, yielding 1 pupa | – 4 crosses<br>– all 123 female pupae **perpetual nymphs**. | n.t. |
|  | *hospiton* male x taxon 2 female | taxon 2 male x *hospiton* female |  |
| *everesti* | – 3 crosses<br>– 38 female pupae developed directly<br>– 1/8 backcrossed females fertile | – 2 crosses<br>– all 10 female pupae **perpetual nymphs**. | 0/3 crosses fertile |

(1) Parents were siblings
(2) Data include Aubert et al.'s (1997); n.t., not tested
(3) In 11 sterile crosses parents were siblings

Similar results were obtained when using *ladakensis* or *everesti* instead of *hospiton*. The female progeny of crosses between *machaon* females and *ladakensis* or *everesti* males consists almost exclusively of perpetual nymphs, whereas the reciprocal crosses yield females that develop directly, without pupal diapause, even when larvae are exposed to long-nights conditions. However, whereas a majority of females obtained from crosses between *machaon* on the one hand, and *hospiton* or *ladakensis* on the other are fertile, male *machaon* x female *everesti* hybrid females were found to be markedly sterile. Most eggs were minute and shapeless, 30% at best would develop and but a single larva was obtained.

The outcome of crosses between *hospiton* and *everesti* is much the same at first sight, with one type of cross resulting in perpetual female nymphs and the reciprocal one in female pupae that do not diapause (Table 2). However, the cross that yields perpetual nymphs is now the one that uses *hospiton* females, whereas when *machaon* is involved, *hospiton* males are required to generate perpetual nymphs.

Contrary to interspecific F1 crosses and backcrosses of hybrid individuals to their parent species, F1 x F1 crosses were expected to be largely sterile. This had been reported to be the case in particular when *hospiton* and *machaon* were involved ((Clarke & Larsen, 1986); (Aubert *et al*., 1997)). Still, one out of the twenty F1 x F1 *hospiton* x *machaon* crosses we attempted proved not only significantly fertile, but sufficiently so for a majority of eggs to develop and hatch and most of the resulting larvae to make it to the pupal and adult stages. We also carried out tests involving *machaon* on the one hand, and *hippocrates*, *saharae* or *ladakensis* on the other (Table 2); in each of these cases a single attempted F1 x F1 cross proved fertile, inasmuch as at least half of the eggs developed, although subsequent larval viability was rather low when *hippocrates* or *saharae* were involved (the latter observations are nevertheless subject to caution, since both crosses involved siblings; we have repeatedly observed that among members of the *machaon* complex, the fertility of sib matings tends to be very low even for intraspecific crosses).

### Larval growth correlates of diapause dysregulation

Hybrid females exhibiting abnormal regulation of diapause (Table 2) were found to be affected as well in larval development. In contrast to the situation in intraspecific crosses, pupation of female larvae destined to become perpetual nymphs is markedly delayed – by several days (Figure S5) – compared to their male siblings. Conversely, females whose pupae are unable to enter diapause tend to pupate earlier than males (data not shown).

Unsurprisingly, lengthening/shortening of the larval stage results in increased/decreased pupal weight. In *P. machaon*, females pupate about one day later than males at 20°C (Figure S5) and we determined the average female/male pupal weight ratio (R_f/m_) to be 1.13 ± 0.01 (Figure S6; similar values were obtained for other members of the *machaon* complex). As shown, however, in Figure 6, quite different values may be observed in some interspecific crosses. Moreover, we found the female/male weight ratio to be highly reproducible for any particular type of cross, at least as long as larvae are well fed and healthy; as an example, four crosses between *ladakensis* males and *machaon* females yielded the following values: 1.60 ± 0.07 (standard error, n=23 individuals), 1.56 ± 0.09 (n=30), 1.55 ± 0.06 (n=31), 1.52 ± 0.12 (n=21).

**FIGURE 6.**
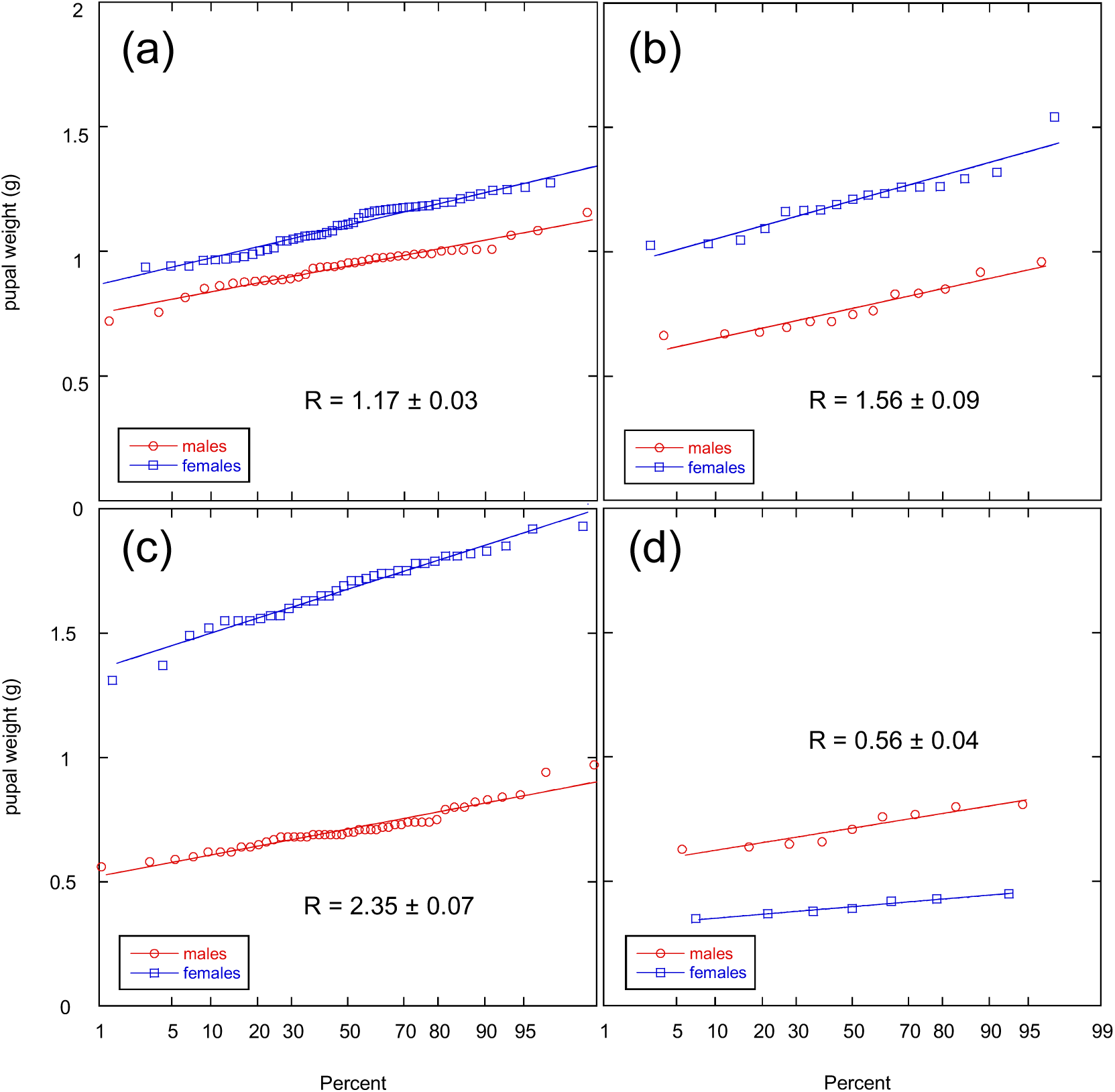
Distribution of female *vs* male pupal weights in the progeny of a *P. machaon* female and three F1 hybrid crosses. Normal probability plots (in which Gaussian distributions are represented by straight lines with a slope proportional to the variance, see (Harding, 1949)) were generated with KaleidaGraph 4.5. (a) Field-collected *P. machaon* female (France, Voutenay); (b) *ladakensis* male (India, Leh) x *machaon* female (France, Rochefort-en-Yvelines); (c) *everesti* male (China, Nyalam) x *machaon* female (France, Saint-Hilaire); (d) *machaon* male (France, Saint-Hilaire) x *everesti* female (China, Nyalam).

Two striking facts emerge from these data. The first one is that severe dysregulation of diapause in hybrids is associated with markedly divergent pupal weight ratios. Crosses between *P. machaon* females and *P. hospiton* males generate hybrid female pupae that may or may not avoid becoming perpetual nymphs depending on rearing conditions (Table 2) and these females are moderately heavier than their male siblings (R_f/m_ = 1.51 ± 0.05). By contrast, we were unable to find conditions that would prevent hybrid females from crosses between *P. everesti* males and *P. machaon* females, which are extraordinarily heavier than their male siblings (R_f/m_ = 2.35 ± 0.07), to escape perpetual diapause. The second point is that pupal weight ratios always diverge in inverse ways in each pair of reciprocal crosses. This relationship is a remarkably quantitative one indeed, as shown in Figure 7a for crosses involving *machaon* on the one hand and *hospiton*, *ladakensis* and *everesti* on the other; when R_f/m_ values for a pair of reciprocal crosses are compared to, i.e. divided by, the R_f/m_ estimated for intraspecific crosses (1.13), almost exactly inverse values result. For example the R_f/m_ for male *hospiton* x female *machaon* and male *machaon* x female *hospiton* crosses are 1.51 and 0.86, respectively and when these numbers are divided by 1.13, the results are 1.336 and 0.761, the product of which (1.017) is close indeed to 1.

**FIGURE 7.**
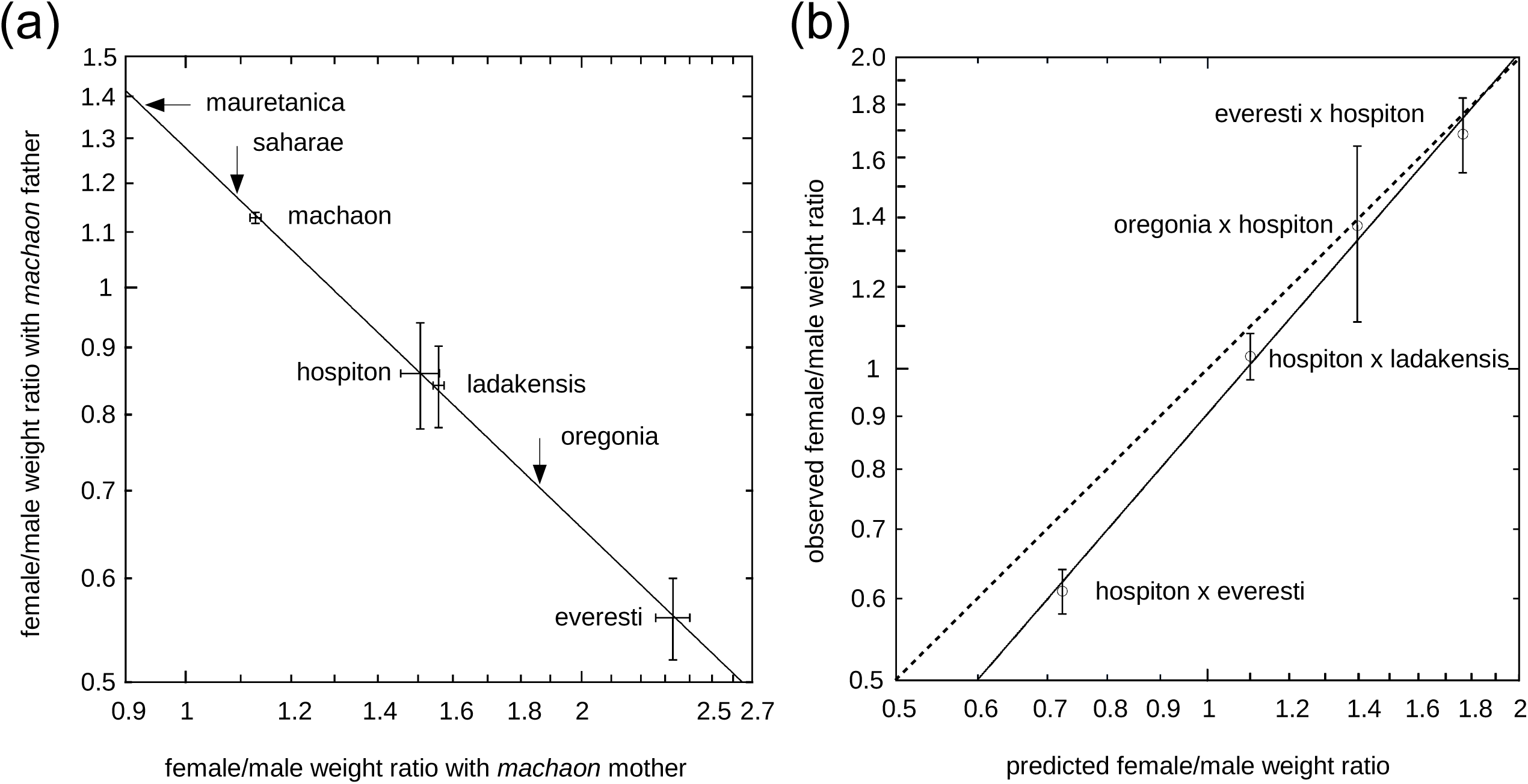
Female *vs* male weight ratios in progenies of reciprocal F1 crosses. (a) Inverse female vs male (R_f/m_) weight ratios in progenies of reciprocal F1 crosses involving *P. machaon*. Bars indicate standard error. Data for *P. machaon* control, *ladakensis* male x *machaon* female and *hospiton* male x *machaon* female are averages of 8, 4 and 2 separate crosses, respectively. Arrows correspond to three combinations – *mauretanica* female with *machaon* male, *saharae* and *oregonia* male with *machaon* female – for which reciprocal crosses were not available. Data for pairs of reciprocal crosses were fitted (continuous line) to a power law, y = 1.278 (x)^-0.960^ (y = 1.005 (x)^-0.960^ would have resulted had all R_f/m_ values been divided by 1.13; this is the average of observed R_f/m_ values in intraspecific crosses, see Results and Figure S6; a power of -1.0 would correspond to a perfectly inverse relationship). (b) Observed compared to predicted female *vs* male weight ratios for crosses that did not involve *P. machaon*. Weight ratios were predicted from data in panel (a) (see Text). For observed values, which correspond to single crosses, bars indicate standard error. Data were fitted to a power law (full line of slope .904); had observed values precisely coincided with predicted ones, a slope of 1.0 (dashed line) would have been observed.

For *saharae* and *oregonia*, only crosses with *machaon* females were available, while for *mauretanica*, the only cross was with a *machaon* male. These three taxa may nevertheless be incorporated into the graph in Figure 7a on the assumption that the inverse effect on pupal weights observed for *hospiton*, *ladakensis* and *everesti* in reciprocal crosses with *machaon* holds as well in their case. The question that then arises is whether the diapause dysregulation scale defined by the diagonal axis in Figure 7a, with *everesti* at one end and *mauretanica* at the other, is specific to the reference species (here *machaon*) or might rather be regarded as an absolute one, not only in the sense that the order of taxa would remain the same whatever the reference taxon, but on quantitative grounds as well. In order to test the latter possibility, several of the taxa in Figure 7a were crossed with *hospiton* and the outcome of these experiments was compared with predictions. For instance, when an *everesti* male is crossed with a *hospiton* female, the expected R_f/m_ would be ((R_f/m_ *everesti* vs. *machaon*) / (R_f/m_ *hospiton* vs. *machaon*)) x 1.13, that is, (2.35 / 1.51) x 1.13 = 1.76. Now, the observed value (Figure 7b) is 1.69 ± 0.14, which is definitely compatible. Also consistent with predictions are (i) the inverse relationship between relative pupal weights for the two reciprocal crosses involving *everesti* and *hospiton*, and (ii) an R_f/m_ not too different from 1 for the cross between a *hospiton* male and a *ladakensis* female, given that the two taxa are located close to one another in Figure 7a (predicted and observed values are 1.07 and 1.03 ± 0.05, respectively).

### Backcrosses to parent species

Backcrossing to *machaon* females the male hybrids generated by pairing *machaon* females with *hospiton*, *ladakensis* or *everesti* males will result in individuals that have inherited a variable part of the Z chromosome of their paternal grandfather within an autosomal context that is predominantly (statistically 3/4) *machaon*. When the progeny of such crosses (between a *machaon* female and a *ladakensis* x *machaon* male hybrid) was reared under long nights conditions, all pupae were found to diapause and virtually all males (159 out 160) resumed development after overwintering. However, only about half of female pupae (63 out of 133) did so and the remaining ones turned out to be perpetual nymphs. Interestingly, and in line with what was observed with F1 hybrids, females destined to become perpetual nymphs experienced delayed pupation and were considerably heavier than their siblings. As shown indeed by comparing Figure S7 and Figure 8 to Figure S5B and Figure 6b, respectively, both the number of days by which pupation is delayed and pupal weight ratios are similar to those observed in F1 crosses. By contrast, females destined to resume development after overwintering pupated shortly after males (they were delayed by half a day on the average – that is, even less than in intraspecific crosses, Figure S5a), and the ratio of their mean pupal weight to that of males (1.08 ± 0.03) was slightly lower than that observed in intraspecific crosses.

**FIGURE 8.**
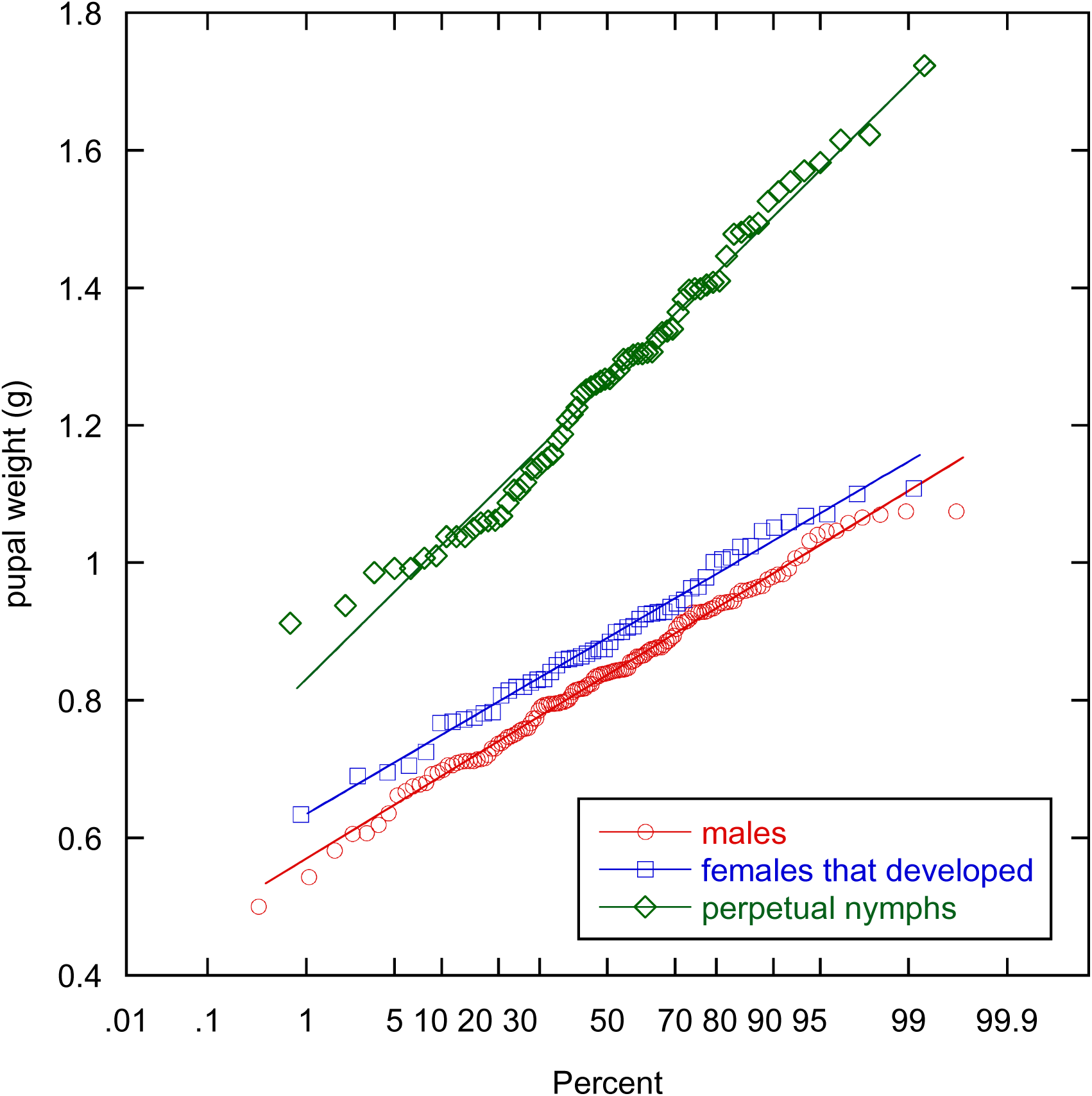
Normal probability plots of the distribution of pupal weights in the progeny of backcrosses of (*machaon* x *ladakensis*) male hybrids to *machaon* females. Data from nine backcrosses were pooled together (Extended Methods). Mean pupal weights: 0.837 ± 0.010 g (n=143) for males, 0.913 ± 0.017 g (n=63) for females that resumed development after diapausing (female/male pupal weight ratio, R_f/m_ = 1.09 ± 0.03), 1.265 ± 0.022 g (n=70; R_f/m_ = 1.51 ± 0.04) for perpetual nymphs.

Further backcrossing was undertaken in order to assess the consequences of introducing foreign Z chromosomes or parts of them in increasingly *machaon* autosomal contexts. The experiments shown in Figure S8 took advantage of the absence of diapause in female F1 hybrids obtained from *hospiton* fathers and *machaon* mothers when larvae have experienced both short nights and elevated temperatures at the same time (otherwise, female larvae would become perpetual nymphs; Introduction and Table 1). One of these F1 females was then backcrossed to a *machaon* male, thus generating hybrids whose autosomal stock was 3/4 *machaon* on average, but which had inherited either a complete *hospiton* Z chromosome, when males, or no *hospiton* Z genes at all, when females. These second-generation hybrids were backcrossed in turn to *machaon* partners, and progenies of male and female hybrids were compared with one another (larvae had been bred under diapause-inducing conditions). In the absence of *hospiton* Z genes, many pupae – in variable numbers from one cross to the next – either did not diapause at all when kept at room temperature or developed into adults after a few additional weeks (Figure S8, panels A and B). Moreover, all remaining pupae gave birth to adults almost simultaneously after having been removed from cold storage. By contrast (panel C), only three pupae with fathers that carried a *hospiton* Z chromosome hatched prematurely and development of at least half of female pupae, as well as a few male ones, was found to be markedly delayed after they had been brought back to room temperature. We interpret the inability of a large fraction of pupae lacking *hospiton* Z genes to experience stable diapause as resulting from the residual presence of *hospiton* autosomal genes (1/8 of autosomal genetic material on average). Presence of part of the *hospiton* Z chromosome, on the other hand, not only protects from this effect but results in much delayed pupal development in many females, even though few of these may be genuine perpetual nymphs (in contrast with the *ladakensis* x *machaon* backcross in Figure 8).

## DISCUSSION

Our investigations of Palearctic members of the *machaon* complex have resulted in two main lines of findings. First, several cases of mito-nuclear discordance in phylogenetic relationships were detected and at least two of these may be ascribed to some form of introgression, either ongoing or recent. Then, we have uncovered several novel instances of dysregulation of diapause in hybrid broods and in all cases that we could thoroughly investigate, the progenies of reciprocal crosses turned out to be affected in inverse ways, not just qualitatively, with female hybrids being unable either to enter, or terminate, diapause (Table 2), but quantitatively as well, as revealed by weighing pupae (Figures 6-8). As introgression and dysregulation of diapause happen to involve the same combinations of taxa, the possibility that the balance between these two phenomena has a decisive role in setting the fate of incipient species after they have been brought into contact needs to be contemplated. Finally, we provide evidence which suggests that dysregulation of pupal diapause in hybrids, which may be more widespread than currently perceived, ensues from evolutionary divergence of parental life histories, a possibility that would contribute to make speciation predictable to some extent.

### ITS2 variation and evidence for introgression

Concerted evolution of rDNA units (Dover, 1982) has long been held to result in ITS sequences being at the same time species-specific and mostly conserved within any particular taxon. However, a rapidly increasing number of reports have uncovered not only significant intraspecific diversity, but intragenomic variation as well (reviewed in (Wang *et al*., 2023)). Still, the extent of variation revealed by our attempts to recover as many ITS2 haplotypes as possible from 44 Palearctic individuals of the *P. machaon* complex other than *P. hospiton* is unexpectedly high. Elevated rates of both mutation and accumulation of variants, coupled with the vagility of adults of the *P. machaon* complex, especially gravid females, undoubtedly concur to the high levels of variation – and information – in our dataset. Even so, the actual diversity of ITS2 sequences in the populations and taxa we investigated is certainly much higher, as should become apparent indeed by sampling larger numbers of individuals and trying to recover additional minor components from PCR product mixtures. It must also be recalled that the PCR amplification process is bound to be biased against some amplicon sequences, which could go unnoticed (e.g. (Keller *et al*., 2006)). Amplification biases are frequently observed indeed when attempting to reveal the presence of ITS2 haplotypes from both parents in hybrids: whenever PCR amplicons happen to have markedly different lengths, the smaller product is generally favored (e.g. Fig. 3 in (Michel *et al*., 2013). Nevertheless, despite *P. machaon* ITS2 amplicons being 10 percent longer than their *P. hospiton* counterparts, they remain readily detectable in the PCR amplification products of a F1 hybrid, or in individuals that inherited the *hospiton* ITS2 sequence in the progeny of a backcross (Figure S9). Reliable quantitative estimates of the relative abundance of individual haplotypes would require cloning of ITS2 DNA or PCR-free, long-read sequencing of rDNA repeats (provided the length of reads is well in excess of that of rDNA repeats).

A related question is whether a majority of observed ITS2 variants are selectively neutral. As discussed in Extended Methods, there is no reason to assume that even such a seemingly major change as the 33-nt deletion in haplotype 127B should have been seriously detrimental to the individual that carried it. If correct, then ITS2 haplotype variants should constitute ideal markers to trace genetic exchanges within and between populations of the *P. machaon* complex. The ITS2 network in Figure 4 does offer several informative examples of genetic continuity within taxonomic units. Three individuals (W106, W142 and W469) whose mitochondrial sequences are too divergent to fall within the same subtrees as their presumed taxonomic relatives (Figure 3) possess nevertheless ITS2 haplotypes that connect them genetically to their geographic neighbors (Figure 4): W469 and *everesti alpherakyi* W103, both from Eastern Qinghai, have similar ITS2 sequences that differ by only three changes; W106, with a typical *ladakensis* wing pattern, shares both its type A and type B ITS2 haplotypes with individual W474, also from Ladakh; and two of the three ITS2 haplotypes of specimen W142 from central Nepal are identical or closely related to those of other individuals (W129, W133) from the southern slopes of the Himalayan range. A similar case of coexistence of divergent mitochondrial haplotypes within populations that are otherwise genetically homogenous has been linked to cycles of fragmentation and merging of populations during and in between Pleistocene glaciations (Hinojosa *et al*., 2019) (Després, 2019); such an explanation seems particularly appropriate for dwellers of Tibet and its surrounding ranges.

ITS2 sequences can also keep track of introgression, during which some genetic markers cross species boundaries. A particularly clear case was provided by the presence, in part of the *everesti* population at Nyalam, of ‘type B’ ITS2 haplotypes, which are quite unlike the ‘type A’ haplotypes of other *everesti* specimens, but closely related instead to sequences from several *machaon asiatica* individuals, including one that was collected along the Kodari Road (Nepal). The latter place is located a mere 30 km south of Nyalam, in the same valley, but at a much lower elevation (ca 1500 m instead of 3700 m). Interestingly, no fewer than five distinct combinations of the three haplotypes (one of type A, two of type B) present in the Nyalam population were recovered from the six specimens examined (Figure 4), but whatever their ITS2 haplotype(s), all adult individuals from Nyalam exhibited a typical *everesti* wing pattern and possessed *everesti*-like mtDNA. Still, among some twenty L5 caterpillars that were encountered at Nyalam on their *Heracleum* (Apiaceae) foodplant (probably *H. candicans*; F.M., unpublished observation), several had a greenish, rather than white background color, and one of them, which looked quite different, was eventually realized to be very similar to lab-generated F1 hybrids between *machaon* and *everesti* (Figure 1, bottom row), an observation which offers independent evidence of genetic introgression from *machaon* into *everesti*.

Since *ladakensis* and *machaon asiatica* individuals have very similar mtDNAs (Figure 3), the ITS2 haplotypes responsible for mito-nuclear discordance are not type B ones, as was the case for *everesti* at Nyalam, but the type A ones (spots J and K in Figure 5b). However, by far the greatest extent of discordance in Figure 5b is associated with spot L, which corresponds to individuals of *oregonia* (and *bairdii*), whose mtDNA hardly differs from that of eurosiberian *P. machaon* (Figure 3), whereas their IT2 haplotypes are either identical or closely related to those of the W494 *P. zelicaon* specimen (Figure S4). This does not come as a surprise, since the nuclear DNA of *oregonia* and sister taxa (*dodi*, *bairdii*) is known to consist of a mosaic of genes from *P. machaon* and allies on the one hand, and *P. zelicaon*/*P. polyxenes* on the other (Dupuis & Sperling, 2020). Even so, it is worth recalling that transfer of ITS2 (or mtDNA, depending on the events that gave rise to *oregonia* and *bairdii*), involved taxa which are believed to have diverged some 7 Myears ago. Although the ITS2 sequences of *machaon* and *zelicaon* still share an uninterrupted segment of 108 identical nucleotides in the distal section of stem III, it seems unlikely that recombination within that segment would lead to functional molecules given the degree of divergence of sequences that flank it on both sides (Figure S3 – the sequences of *zelicaon* and *polyxenes* are closely related); as discussed in Results, clonal inheritance of highly divergent ITS2 variants segment could account indeed for relative rates of divergence of ITS2 sequences on the one hand, and mtDNA on the other, tending towards a fixed value, as seen in Figure 5b. Note also that contrary to the three *oregonia* specimens we examined, the single *P. brevicauda* individual that was investigated has retained a *machaon*-like ITS2 haplotype (Figure 4; as indicated by spot F being on the curve fitting points A to G in Figure 5b, observed relative extents of mitochondrial and ITS2 divergence are consistent with vertical inheritance in this separate hybrid lineage (Dupuis & Sperling, 2020)).

Identifying candidate introgression events would not have been possible in the absence of a fair degree of overall concordance between mitochondrial DNA-derived (Figure 3) and ITS2-derived (Figures 4,5) phylogenies. According, however, to (Dupuis & Sperling, 2020), nuclear SNP data are not in favor of the generally accepted phylogeny of the *P. machaon* complex, as inferred from mitochondrial sequences. Among these authors’ conclusions, a particularly unexpected one was that rather than being sister to *P. machaon* and closely related Palearctic allies, *P. hospiton* diverged in fact much earlier, being ‘sister to the remainder of the species group (excluding *P. indra*)’. Reanalysis of their SNP data (Figure S10a) does seem to support their assertion, at least as long as all of the 19 specimens that were selected to generate a ‘phylogenetic dataset’ are retained. However, removal of the ten or so individuals of proven or presumed hybrid origin results in a different, well-supported nuclear DNA phylogeny (Figure S10b) which fully agrees with its mitochondrial counterpart. Overall phylogenetic concordance of several separate DNA datasets (ITS2 sequences were absent from Dupuis and Sperling’s SNP dataset) leaves little doubt that the hierarchy of ancestral divergence events has been correctly recovered.

### Symmetrical outcomes of reciprocal F1 crosses and correlation with differences in parental voltinism

In each of the four cases of reciprocal hybrid crosses we were able to investigate, female over male pupal weight ratios (R_f/m_) were found to take almost exactly inverse values once numerical data were corrected for the mean R_f/m_ determined in intraspecific crosses (Figures 7a,b). By assuming that such symmetry is a general rule, we could place all six members of the *machaon* complex for which we had at least one cross with the reference taxon (*P. machaon*) on a single diagonal axis (Figure 7a) according to their associated R_f/m_ ratio(s). Remarkably, we found that the outcome, in terms of R_f/m_ values, of any cross that did not involve *machaon* could then be predicted quantitatively from the locations on that axis of the two entities that had participated in the cross. In other words, relative positions on the ‘diapause dysregulation’ scale in Figure 7a do not depend on the reference species or population, but reflect an intrinsic property of individual taxa (a ‘trait’) and we regard this as highly suggestive evidence that one and the same regulatory system is involved in the fate of all hybrid progenies we generated. We suggest to call provisionally this trait ‘*dDiap*’, which could be meant to be read ‘dysregulation of diapause’, but also, alternatively, ‘depth (intensity) of diapause’, as discussed below.

As might have been expected, average pupal weights of different categories of individuals in the same cross were found to correlate closely with the duration of the larval stage and particularly so with the number of days spent in the 5^th^ (last) larval instar (some moderate larval asynchrony in the progeny of crosses that give rise to perpetual nymphs is already manifest at the time of the L4-L5 moult). Compared to males, which spend some 8-9 days in L5 at 20 °C, pupation of hybrid females is liable to be delayed by as many as five (Figure S5B) or more days, resulting in R_f/m_ ratios that can exceed twice the average value for intraspecific crosses (in the progeny of the cross between a male *everesti* and female *machaon*), or else advanced by up to two to three days (data not shown), whereas the normal interval between average male and female pupation, resulting in a R_f/m_ of *ca* 1.13 (Figure S6), typically amounts to just one day in intraspecific crosses (Figure S5A).

In addition to being correlated with one another, duration of the last larval instar and pupal weight were found to be predictive of the fate of pupae in interspecific crosses. Females that turned out to be unable to enter diapause, despite having been reared under long nights, pupated early and were lighter on the average than their brothers, while the converse is true of perpetual nymphs. In the backcross experiment shown in Figure 8, just 11 out of 70 perpetual nymphs weighed less than a ‘critical’ weight of 1.055 g, whereas only 11 out of 63 females that underwent direct development weighed more. This observation is compatible with the possible existence, within the continuous distribution of weights, of a threshold that makes it possible to acquire a discrete character, ‘perpetual’ rather than ‘temporary’ nymph. However, while we could not find conditions that would allow larvae destined to undergo obligate direct development to avert that fate, there appears to exist a graded scale of situations from normal to compulsory, permanent diapause. Irrespective of larval rearing conditions, crosses between an *everesti* male and a *machaon* female invariably yielded giant female pupae, which entered irreversible diapause. By contrast, the fate of hybrid females from male *hospiton* x female *machaon* crosses, which are of intermediate weight (Figure 7), depends on breeding conditions (Aubert *et al*., 1997): female larvae reared under short nights and at elevated temperatures escape from becoming perpetual nymphs. Also worth noting, in the progeny of these and male *ladakensis* x female *machaon* crosses, a few diapausing females – 2 out of 24 in the latter crosses – managed to resume development after a total of some 16 months at 2°C (Extended Methods; the remaining, dormant pupae were subsequently verified to respond positively to ecdysone injection; see (Clarke & Willig, 1977)).

Yet another apparent correlation is between relative weights of hybrid females and their propensity to become perpetual nymphs on the one hand, and voltinism of parental species on the other. In crosses that give rise to permanently diapausing females, the father comes from a species or population that tends to have fewer generations per year than the maternal one. An extreme case is provided by male *everesti* x female *machaon* crosses, which are located at the very end of the diapause dysregulation axis (Figure 7). At low altitudes, French *P. machaon* populations are multiple-brooded, with two to three generations per year. By contrast, the possibility of a second generation can be ruled out for the *P. everesti* population at Nyalam (3700 m, from which our stock originated), since adults hatch from overwintering pupae in mid-June and their progeny had not yet pupated in early August 1987 (when larvae were bred indoors under short-night conditions, a single female – out of 36 pupae – underwent direct development). At the other end of the spectrum, *mauretanica* is multiple-brooded (three generations per year in most of its range), while *saharae* has two generations, although the second one may be skipped in dry years (Tarrier & Delacre, 2008). Crosses whose R_f/m_ values fall in between these extremes correspond to more complex situations. Thus, *P. hospiton* is usually single-brooded (although some individuals may skip diapause when raised indoors), but the Corsican populations that use *Peucedanum paniculatum* as larval host-plant have a partial second generation, which can be abundant (Aubert *et al*., 1996) (Aubert *et al*., 1997). Similarly, *P. ladakensis* is generally thought to be univoltine and is almost certainly so North of Leh (Ladakh), at 4400 m. To our surprise, however, our laboratory stocks proved responsive to the ambient photoperiod, as all pupae obtained from larvae raised outdoors in May-June underwent direct development, whereas all larvae that had been bred in late summer gave rise to diapausing pupae. In the case of *oregonia*, however, the correlation does not appear to hold, since this taxon is definitely bivoltine throughout its range in Northwestern United States.

### Perpetual nymphs in the entomological literature

The entomological literature offers numerous records of lepidopteran hybrid pupae that do not hatch and eventually die, but for these to qualify as genuine perpetual nymphs, pupal viability should have been verified by ecdysone injection. Among studies that meet this requirement, those of the *Papilio glaucus* group were pioneering ones. *Papilio glaucus* and its close relatives are swallowtail butterflies (Papilionidae), like *P. machaon*, but the two groups colonized temperate North America independently from one another – their most recent common ancestor is estimated to have lived some 31 Myears ago (Condamine *et al*., 2012). *Papilio glaucus* itself is multiple-brooded throughout its distribution area, in Eastern USA and Northeastern Mexico and a rather narrow hybrid zone in Northern USA separates it from its sibling species, *P. canadensis*, which experiences obligate pupal diapause. Crosses between *glaucus* females and males of the western American species, *P. eurymedon*, *P. rutulus* and *P. multicaudatus*, revealed marked dysregulation of diapause in the female progeny (Clarke *et al*., 1972); (Clarke & Sheppard, 1957); (Clarke & Willig, 1977); (Clarke *et al*., 1989); (West & Clarke, 1988); (Scriber *et al*., 1990); (West, 1995). As summarized in the most comprehensive study (Scriber *et al*., 1990), the highest degree of incompatibility was with *eurymedon*, which is strictly single-brooded, except in parts of Southern California: 25 crosses yielded 223 males, as against a single adult female (and 250 pupae, that eventually died). Then came *rutulus*, with 362 males, 12 females and 407 dead pupae (26 crosses) and finally *multicaudatus*, with 30 males, 6 females and 31 dead pupae; the latter is single-brooded in Northwestern USA, but multiple-brooded further South (Scott, 1986). By contrast, and as also observed in the *machaon* group, females were fully viable in reciprocal crosses. Moreover, as noted by (West, 1995), 8 out of 10 females in the progeny of a cross between a *glaucus* male and an *eurymedon* female were found to undergo direct development, whereas all their brothers diapaused. Worth mentioning as well, reciprocal crosses between *eurymedon* and *rutulus* did not reveal any significant sex-ratio bias among adult progeny.

With regard to *canadensis*, a large number of reciprocal crosses with *glaucus* (summarized in (Rockey *et al*., 1987); larvae were reared on short-night conditions) revealed that in male glaucus x female canadensis F1 progenies, less than 10 % of individuals diapaused, whereas when the male parent was *canadensis*, 96.7 % of females did, as against 17.8% of males (dysregulation of diapause was moderate, since many females eventually developed, but it is unclear whether some of the even larger number of still undeveloped pupae at the time the paper was published were genuine perpetual nymphs). In addition to *canadensis*, the USA are home to at least one other member of the *glaucus*-group, called *P. appalachiensis* (Pavulaan & Wright, 2002). This single-brooded, belatedly described taxon, was shown to be a stable hybrid species with a *P. glaucus*-like mitochondrial DNA and predominantly *P. canadensis*-like Z chromosome (Kunte *et al*., 2011). Remarkably, although predictably so, out of ten pupae from a cross between a *P. appalachiensis* male and a *P. glaucus* female, the seven males eclosed after overwintering whereas the three females failed to do so (Pavulaan & Wright, 2002).

To summarize, this vast mass of data is definitely in favor of the above-stated rule, according to which, in hybrid crosses that generate perpetual nymphs or, at least, a large excess of females among diapausing individuals, the male parent tends to have fewer generations per year than the female one.

Perpetual nymphs of hybrid origin are not limited to Papilionidae, nor to butterflies. Among moths, the genus *Actias* (Saturnidae) offers a particularly convincing illustration of the link between discordant parental voltinisms and the presence of perpetual nymphs in hybrid progenies. When males of *Actias isabellae*, the only European species, confined to the French Alps and Spanish mountains at moderate elevations and strictly univoltine, are crossed with multivoltine females of either temperate (*A*. *luna*, *A*. *artemis*; (Adès *et al*., 1989); (Adès *et al*., 2005b)) or tropical/subtropical origin (*A*. *truncatipennis*, *A*. *selene*, *A*. *sinensis*, *A*.*isis*; (Adès *et al*., 1994); (Adès *et al*., 1993); (Adès *et al*., 2005a); (Vuattoux *et al*., 2001)), all F1 female pupae remain in diapause until they die or are injected with ecdysone. By contrast, female pupae from a cross between *A. sinensis* and *A. dubernardi*, whose ranges overlap in South-East Asia, develop normally (Renner *et al*., 2006). Also in keeping with expectations, female adults obtained by crossing a (tropical) *A. maenas* male with a (temperate) *A. luna* female were found to hatch before their brothers (Vuattoux, 1998).

In Sphingidae as well, and particularly in the genus *Hyles*, some interspecific crosses have long been known to generate female perpetual nymphs ((Loeliger & Karrer, 1996) and references therein). From such crosses, a series of hierarchical relations can be inferred, just as in the *P. machaon* complex. However, connections with voltinism remain somewhat unclear, one reason being that more often than not, authors failed to mention where the stocks they used came from.

Geographical variation in the way species response to the photoperiodic signal, especially when they have a broad geographic distribution can be spectacular indeed. In an extreme example, *ca* 90 % of *Pieris napi* larvae from Abisko (N. Sweden) were found to give rise to diapausing pupae when raised under near-continuous light (Light-Dark (LD) 23:1 hr), whereas 100 % of larvae of the same species from Barcelona underwent direct development under the same conditions (Pruisscher *et al*., 2017). Now, when the two stocks were crossed, percentages of diapausing F1 females were found to differ utterly in reciprocal crosses: under a 18L:6D regimen, 100 % of female larvae from the male Abisko x female Barcelona cross turned into diapausing pupae, whereas only 8.3 % of those from the reciprocal cross did (the response of males was an intermediate one – 40 to 60 % of diapausing larvae – as is generally found when populations with different photoperiodic thresholds are crossed (Danilevskii, 1965)). Not only does this study outline the role of the Z chromosome in the regulation of diapause initiation, but it also provides a striking example of reciprocal crosses with quantitatively inverse outcomes for females, when males are taken as reference.

It is intriguing that the symmetry apparent in Pruisscher et al.’s reciprocal *P. napi* crosses and in Table 2 and Figure 7 of the present work was generally ignored in papers reporting on perpetual nymphs. One reason could be that the progeny of crosses reciprocal to those yielding perpetual nymphs was most often reared under the same short-nights conditions, something which would conceal the possible inability of females to enter diapause. Another possibility is that differences in duration of the larval stage were deemed anecdotal. By contrast, rates of development and symmetry were made central in a somewhat overlooked piece of work by Charles G. Oliver (Oliver, 1983). Drawing on his experience with interspecific crosses in the genus *Phyciodes* (Nymphalidae), this author concluded that reciprocal crosses obeyed inverse trends when female over male relative developmental times were compared. Moreover, in some experiments in which females developed slowly, all of them would diapause, when none of their brothers did (diapause occurs at the larval stage in *Phyciodes*). Additionally, a correlation with voltinism was apparent: when *Phyciodes* taxa in Table 1 of (Oliver, 1983) are arranged hierarchically according to the identity of the male and female parents in crosses that yielded slow-/fast-developing females, *P. batesii* (univoltine) is seen to stand at one extremity, followed by *P. pascoensis* (nowadays *P. cocyta*, (Wahlberg *et al*., 2003)) and *P. pratensis* (=*pulchella*), which are uni- and uni- to trivoltine, respectively; then comes *P. tharos* (multivoltine) and, finally*, P. phaon* (multivoltine as well, but with the most southerly distribution of all; data on voltinism are from (Opler, 1999)).

### Interpreting correlations between developmental rates, inability of female hybrids to initiate/exit diapause and parental voltinism: a formal model

The correlation we observe between the duration of the last larval stage, pupal weight and the ability of pupae either to enter or exit diapause inevitably suggests the existence of a direct causal relationship. Let us assume that during the last larval instar (L5 for *Papilio*), well-fed caterpillars accumulate some ‘diapause effector’ – a substance or molecular complex, the amount of which at the end of the growth period determines whether development will either go on or stop shortly after pupation. Then, upon exposure to winter conditions, some mechanism in diapausing pupae actively and progressively removes the same effector until a second threshold is reached, after which development can resume. Thus, too short the last instar (and too light a pupa) would directly lead to inability to initiate diapause, for want of a sufficient quantity of the effector, whereas the longer the last instar (and heavier a pupa), the more likely would it be for the system in charge of depleting the diapause effector to reach saturation before the critical threshold for resumption of activity is attained, and this would result in a permanent nymphal state.

Should such a system actually operate under normal physiological conditions, one would expect the last instar to last longer in larvae destined to give rise to diapausing pupae than in those that will experience direct development. Now, as discussed in (Denlinger, 2022), this appears to be frequently, although possibly not always the case. For example, Masaki and colleagues (Watanabe *et al*., 1970) reared larvae of the moth *Hyphantria cunea* (Arctiidae) individually under well-controlled conditions, which included the use of an intermediate photoperiod (14.5 h light) that induced diapause in about 3/5 of larvae. Under those conditions, duration of the larval stage did not differ significantly until the last instar. During the latter, however, feeding was found to last definitely longer (by 15 and 19 % in males and females, respectively) in larvae destined to enter pupal diapause when compared to non-diapausing ones. The difference was found to be even larger (32 % and 36 % for males and females) when photoperiods of 13 and 16 hours were compared. In the butterfly *Pieris napi* (Pieridae) as well, (Friberg *et al*., 2012) observed that ‘development time was not consistently different until the ultimate instar’. However, ‘In the ultimate instar, individuals preparing for diapause spent about a day longer before pupating than the direct developers did’. Under the conditions used by these authors, this corresponds to an increase of about 33 % in the duration of the last instar.

Any model that aims to explain why dysregulation of diapause is female-specific in F1 crosses must include some asymmetry in the roles of the Z chromosome and autosomes. We propose that the length of the last larval instar, which we found to be predictive of the ability to enter, and subsequently exit from, diapause, is governed by the balance between two ‘substances’, one of which, ‘Z’, is under control of one or several loci on the Z chromosome, while the other one, ‘A’, has its concentration adjusted by a number of autosomal loci. Let us assume (arbitrarily, since the system is symmetric) that increasing the [z]/[a] ratio will lengthen the last instar, whereas decreasing it has the opposite effect. Photoperiodic control can then be introduced in a number of ways: we chose somewhat arbitrarily to have it operate by modulating the concentration of product Z, using an ‘hourglass timer’ for the sake of simplicity (Materials and Methods). Figure 9 shows first (panels A-B) how an evolutionary trajectory along which a multivoltine entity (species or population) is transformed into a univoltine entity can be modelled; a key point is that it is possible to avoid ending with truly giant pupae under ordinary circumstances – i.e. in a well-adapted, single-brooded species – by increasing (differentially) the values of both [z] and [a] at the same time. Then, in panels C-E, it is shown why crosses between two such species or incipient species as in panels A and B should result in dysregulation of growth and diapause in female, but not male, hybrids. The reason is that male F1 hybrids have two copies of all chromosomes – one from each parent -- so that the photoperiod response curve is intermediate between the parental ones ([a] and [z] concentrations are themselves assumed to be intermediate). By contrast, in female hybrids, the curves are shifted beyond either one or the other parental curve, depending on which Z chromosome females inherited ((Huylmans *et al*., 2017) observed full dosage compensation for non-sex-biased genes in *Papilio machaon*; accordingly, we assumed that the contribution of individual Z chromosomes to the concentration of the Z product is halved in males compared to females, irrespective of the actual mechanism of dosage compensation).

**FIGURE 9.**
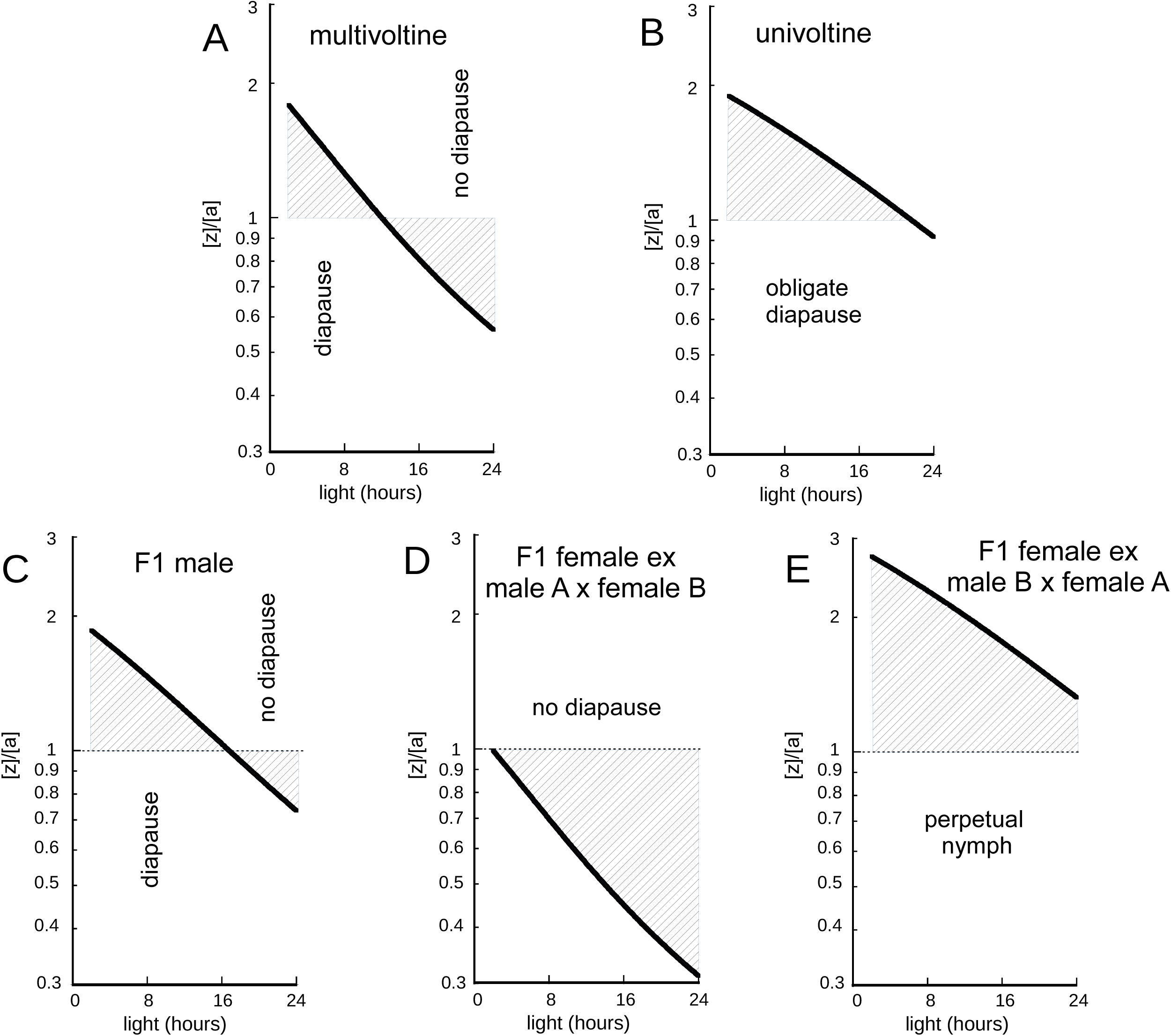
Modelling the photoperiodic response along an evolutionary trajectory from multi- to univoltinism and in the F1 progeny of a cross between a multiple-brooded and a single-brooded taxon. Modelling was as described in Materials and Methods. Values of parameters z_T_ and [a], the ratio of which is proposed to control pupal diapause (see Text), were adjusted so as to illustrate qualitative consistency with experimental data in general (i.e. not to fit a particular dataset). (A) Multivoltine taxon, z_T_ = 1, [a] =0.5; (B) Univoltine taxon, z_T_ = 2.6, [a] =1.3; (C) F1 male hybrid between taxa A and B, z_T_ = 1.8, [a] =0.9; (D) F1 female hybrid between male A (multivoltine) and female B (univoltine), z_T_ = 1, [a] =0.9; (E) F1 female hybrid between male B (univoltine) and female A (multivoltine), z_T_ = 2.6, [a] =0.9.

### Prospects for identifying genes involved in dysregulation of diapause in hybrids

Despite its efficiency in accounting for our and others’ observations, the above formal model may be perceived as somewhat disconcerting. First, it is unlike explanations most commonly put forward to account for X/Z-autosomal incompatibilities at the root of Haldane’s rule (reviewed by (Cowell, 2022)); these rely on the concept of genetic dominance, which fails to predict the systematically symmetric outcome of reciprocal crosses which we observe. Alternate, more complex systems that rest on networks of regulators (e.g. (Lenormand & Roze, 2025)) need be invoked, as we have attempted indeed. Then, a model that makes use of only two adjustable parameters and makes no specific prediction about the identity of genes possibly involved cannot hope to do justice to the complexity and subtlety of actual physiological interactions that control insect diapause (reviewed in (Denlinger, 2022)), nor be of much practical use to physiologists.

There have been a fair number of attempts to identify genes involved in the control of diapause in Lepidoptera by genetic analyses. One approach, exemplified by (MacDonald *et al*., 2020) for *P. machaon dodi* and (Cong *et al*., 2015) and (Ryan *et al*., 2017) for the *P. glaucus*/*P. canadensis* pair of taxa consists in genome-wide comparisons of populations or sibling species that differ in diapause induction, termination and/or voltinism. These holistic endeavours generate long lists of hot spots for divergence and/or positive selection, but the interpretation of these lists requires additional information, either from traditional genetic analyses (usually of backcrosses) in similar systems or the physiological literature.

In an early attempt to locate genes in control of diapause on the Z chromosome, (Hagen & Scriber, 1989) backcrossed F1 *glaucus* x *canadensis* hybrid males to *glaucus* females. Seven successful backcrosses yielded 92 viable adult females, 41 of which had diapaused after having been reared under short nights (the actual fraction of diapausing pupae was larger, since an unspecified number of – presumably female – pupae were injected with ecdysone after one year and only about half of these made it to the adult stage). Most diapausing females were found to have *canadensis* alleles at two out of the three Z-located loci examined, which was interpreted as evidence of strong linkage between those two loci and a (hypothetical single) locus responsible for obligate diapause in the *canadensis* parent.

Much more recently, (Pruisscher *et al*., 2018) compared by genomic scans two Swedish populations of the butterfly *Pararge aegeria* (Nymphalidae) with markedly different photoperiodic thresholds for the induction of pupal diapause. Among particularly divergent DNA sections, 15 candidate loci were selected and their possible association with diapause phenotypes was examined by means of an F2 cross in the progeny of which 29% of females as against 13.6% of males had diapaused (as in other similar systems, reciprocal F1 crosses differed significantly in the fraction of diapausing females, but not of diapausing males, and in the cross yielding the highest percentage of the former category, the father was from the northern population, 98% individuals of which had diapaused in control crosses as against 2% from the southern population). Significant association was found between propensity to diapause and alleles at three genes within a *ca* 150 kb autosomal DNA section, much of which appeared to be under strong selection. One of these genes was *timeless*, a circadian clock gene, and strikingly enough, characterizing individuals in three populations with intermediate critical photoperiods, in addition to the two original ones, revealed an abrupt change along a North-South gradient in allele frequencies at that locus, with no heterozygotes. Moreover, the same was found to be true of another circadian clock gene, *period*, which is located on the Z chromosome and had been found to be marginally associated with diapause phenotypes in females.

In yet another study, (Pruisscher *et al*., 2021) attempted to identify loci involved in the control of diapause induction in *P. napi* by backcrossing to Swedish females some hybrid males with Swedish and Spanish parents (Pruisscher *et al*., 2017). Sequencing and comparing pools of directly developing and diapausing individuals in five backcrosses suggested that only chromosome Z was involved. However, no particular region of that chromosome stood out until the distribution of hotspots of sequence divergence between the Swedish and Spanish populations was taken into account. By further restricting the search to exons of annotated genes and fixed, non-synonymous differences, the authors eventually reached the conclusion that just as in other systems, *period* was the most likely candidate for the control of diapause induction.

It is instructive to compare this piece of work with a rather recently published paper (Xiong *et al*., 2023) in which genomic analyses of backcrosses between sibling *Papilio* species (*P. bianor* and *P. dehaanii*) were carried out in order to try and find genes involved in the control of pupal weight in hybrid females: just as shown in Figure 7 of the present work, the authors had found these weights to differ in inverse ways in reciprocal F1 crosses when hybrid males were taken as reference. As in (Pruisscher *et al*., 2021), no significant association was found with any autosomal locus, whereas the Z chromosome appeared to contribute to hybrid female pupal weight almost continuously over its entire length. While the authors acknowledged the difficulty of choosing between a fully polygenic model and one with two or more loci as major contributors, they clearly favored the former.

One potential problem with backcrosses of F1 hybrids is the simultaneous segregation of autosomal and sex chromosomal markers, which makes it difficult to untangle individual contributions. Such complexity may be alleviated in part by recurrent backcrossing, an example of which is provided in Figure S8. It is our hope that by combining this approach with the simultaneous use, as in Figure 8 of this work, of a quantitative trait (pupal weight) and related qualitative threshold traits (diapause initiation and/or termination), we will be able to identify discrete sections of DNA that contribute to postzygotic incompatibilities in hybrid individuals.

### Species delimitation: to what extent does diapause dysregulation in hybrids may contribute to isolation from neighbors?

Extracting reliable information about dysregulation of pupal diapause in crosses involving Nearctic relatives of *P. machaon* is a complex, but not impossible task, as may be realized by reading our Extended Discussion (Supplementary information). It will suffice to state here that once Nearctic taxa are taken into account, the diapause dysregulation axis in Figure 7 reads tentatively as follows (groups of taxa flanked by vertical bars cannot be further divided at present due to insufficient quantitative data):

*mauretanica* | *machaon*,*hippocrates*,*saharae*,*polyxenes* | *britannicus*,*brevicauda,joanae* | *hospiton*,*ladakensis* | *zelicaon*,*oregonia* | *everesti*

Assuming that dysregulation of pupal diapause in female hybrids increases together with the distance between taxa along that scale, as we found to be the case in the subset we could study, we may then ask whether some relationship appears to exist between the magnitude of diapause dysregulation as measured in laboratory settings and the extent of genetic exchanges between those taxa that happen to be in contact in nature.

It must be made clear from the outset that not even the highest observed degree of post-zygotic incompatibility – in crosses between *machaon* and *everesti* – completely prevents genetic exchanges. Not only are F1 hybrid females from these crosses either unviable or sterile, but about half of the daughters of a F1 hybrid male backcrossed to *machaon* will turn into perpetual nymphs (our own unpublished observations, see also Figure 8 for *ladakensis*/*machaon* backcrosses; while the *machaon* stocks used in our laboratory crosses were from France, we expect that controls with local, multivoltine *machaon* populations – here from the South slope of the Himalayan range, at elevations below 2000 m – would result in essentially identical outcomes). Still, both larval morphology and ITS2 sequence data provide undeniable evidence of introgression of *machaon* genes into *everesti* at Nyalam. This is all the more remarkable that despite the distance between the *machaon* population along the Kodari Road and the *everesti* population at Nyalam not exceeding 30 km along the Sun Kosi river, the surroundings are extremely different, being subtropical at an elevation of 1500 m and subalpine at 3700 m at Nyalam (Dobremez, 1974). Whether populations belonging to the *machaon* complex inhabit intervening biotopes is unknown to us, but it must be recalled that *P. machaon* is a powerful flier, capable of crossing the 100 or so km that separate British Dorset from the French coast ((Collins *et al*., 2020) and references therein). What is truly surprising is that mechanisms should not have evolved to prevent more effectively interspecific matings in this complex, given their possible cost in terms of reproductive efficiency. Intraspecific courtship is minimal and brief whether for *P. machaon hippocrates* (Takeuchi, 2019) or *P. polyxenes* (Lederhouse, 1982) and in the case of *P. hospiton*, hybrids with *machaon* are frequently encountered in nature ((Aubert *et al*., 1997), and references therein).

At Nyalam, introgression from *machaon* does not appear to affect adult wing patterns, which were typical of *everesti* in all individuals examined. Nor was *machaon* mtDNA detected, as expected from F1 hybrid females with a *machaon* mother being perpetual nymphs. Moreover, pupal weight distributions in progenies of crosses with *machaon* and *hospiton* did not differ significantly between *everesti* individuals (a total of ten wild-collected individuals with different ITS2 genotypes were tested; data not shown). While it remains to be determined how far to the North of Nyalam may introgression be traced, it should be recalled that we did not find mtDNA or ITS2 sequences from *machaon* in *everesti* individuals collected at five additional locations in Tibet and its surroundings.

Along their zone of contact, which extends from Southern Alberta to Northern New Mexico, Rocky Mountains *P. zelicaon* and Great Plains *P. polyxenes* offer another instance of limited introgression in the presence of strong physiological incompatibility coupled with a rather steep ecological gradient, as multiple-brooded *polyxenes* occurs at typically lower elevations than single-brooded *zelicaon*. A predominantly black, rather than yellow, form of *P. zelicaon* called *nitra* is thought to have originated through hybridization with the ‘Black Swallowtail’ (*P. polyxenes*), for it is seldom encountered except in the vicinity of the area of distribution of *P. polyxenes*, where its frequency can reach up to 20 percent (Scott, 1986). Otherwise, molecular evidence of contemporary genetic exchanges between the two species, which have distinct mtDNAs, is essentially lacking (Dupuis & Sperling, 2020).

Incompatibility between *machaon* and *hospiton* is marked, but not to the point of completely preventing females from a male *hospiton* x female *machaon* cross to reach the adult stage, since larval growth at such elevated temperatures as encountered in late spring at low elevations in Corsica and Sardinia should allow at least some of them to escape from becoming perpetual nymphs (Aubert *et al*., 1997). However, as a majority of *hospiton* populations are single-brooded ((Aubert *et al*., 1996) and references therein), the males that these hybrid females (and also those with a *machaon* father and *hospiton* mother, which undergo obligate direct development) will encounter should belong predominantly to *P. machaon*. Similarly, non-diapausing male F1 hybrids will have little choice but to mate with *machaon* females. This asymmetry may be the main reason why introgression was found to be almost exclusively from *hospiton* towards *machaon* (Cianchi *et al*., 2003). However, populations of *P. hospiton* that utilize *Peucedanum paniculatum* as larval foodplant in Corsica have a second, summer brood, which may be abundant (Aubert *et al*., 1997), so that opportunities should exist for hybrids to mate with *hospiton*. It has been proposed indeed that in those populations, the potential for multivoltinism resulted from introgression from *P. machaon* (Aubert *et al*., 1997). Still, (Cianchi *et al*., 2003) found only limited evidence of introgression in a sample of 27 *hospiton* individuals from one of these populations (a set of nine diagnostic allozyme loci was used). In any case, it should be of interest to perform further crosses of individuals from multiple-brooded *P. hospiton* populations with *P. machaon* and determine precisely the extent of diapause dysregulation in their progenies.

At the other end of the incompatibility spectrum lies the pair formed by *zelicaon* and *oregonia*, whose female hybrids do not appear to suffer from diapause dysregulation (Thompson, 1988a). Interestingly, *zelicaon* and *dodi* (a close relative of *oregonia*) have been reported to generate ‘hybrid swarms’ in Southwestern Alberta (Sperling, 1987). The area within which hybridization is intense is rather circumscribed, possibly because of stronger ecological differentiation in other parts of Alberta (Dupuis *et al*., 2019). Nonetheless, it seems unlikely that the level of admixture observed in the core of the hybrid zone could have been reached in the absence of physiological compatibility.

*Papilio machaon mauretanica* and *saharae* constitute another pair of taxa located close to one another on the diapause incompatibility scale of Figure 7. Speculations about whether *saharae* should be regarded as a subspecies of *machaon* or a distinct species have a long history (e.g. (Clarke & Larsen, 1986) (Pittaway *et al*., 1994)). The *saharae* larva is highly distinctive, but populations with larval morphologies intermediate between *saharae* and *mauretanica* exist in Morocco (Figure 1; (Tarrier, 2015)). Adults of the two entities cannot be reliably separated from their wing patterns, but they differ somewhat in genitalia and number of antennal segments ((Eller, 1936) (Pierron, 1990) (Pittaway *et al*., 1994)). The rather low viability of both our and Clarke’s (Clarke & Larsen, 1986) single *machaon* x *saharae* F2 crosses (Table 2) is inconclusive, since it could result from consanguinity – parents were siblings in both cases – as well as from genuine incompatibility (Pierron’s claim that *saharae* x *hospiton* crosses revealed higher genetic compatibility than those involving *saharae* and *machaon* (Pierron, 1990) was not appropriately documented).

As seen in the mitochondrial phylogeny in Figure 3, a well-supported subtree groups together all *saharae* individuals we examined, whether from North Africa or Yemen. However, this subtree includes all three *mauretanica* individuals of our dataset as well. Moreover, a recently published study by (Cassar *et al*., 2025) about genomic differentiation of Mediterranean members of the *machaon* complex includes a mitochondrial phylogenetic tree (Fig. S2) in which a *saharae* clade (only North African and Lampedusa specimens were investigated) also comprises three *mauretanica* specimens. Actually, while ITS2 haplotypes (Figure 4 of the present work) do not offer sufficient resolution to separate *saharae* from either *mauretanica* or North Mediterranean *machaon* specimens, data obtained by (Cassar *et al*., 2025) not only confirmed that nuclear genomic sequences do not make it possible to distinguish *mauretanica* from *saharae*, but revealed that the two entities diverged from European *machaon* no more than 0.44 Myears ago, according to these authors’ calibration. That latter date is about three-fold lower than our own estimate (*ca* 1.5 Myears ago, Figure 3), based on mitochondrial DNA divergence. However, other divergence times proposed by Cassar, Condamine and colleagues (e.g. *ca* 1.35 Myears for divergence of *hospiton* and *machaon*, *ca* 0.95 Myears for separation of *zelicaon* and *machaon*; Fig. 4) are utterly incompatible with all previous estimates, including those from the Condamine lab (e.g. 5.2 and 6.7 Myears for divergence of *hospiton* and *zelicaon*, respectively, from *machaon* in (Condamine *et al*., 2023)). Such discrepancies cannot but question the pertinence of elaborate paleobiogeographic inferences when interpretable fossils are lacking.

Cassar et al. concluded from their data that *saharae* (including *mauretanica*) and *machaon* are distinct species, in spite of rather low molecular divergence (see also Figure 3). However, while pooling *mauretanica* with *saharae* results in *machaon* and *saharae* being allopatric in the Western Mediterranean Basin (at least until putative records of *saharae* from Sicily (Moonen, 2012) (Cassar, 2018) are documented at the DNA level), the two entities do coexist in Southern Israel, where they generate hybrid populations (Benyamini & John, 2020) (actually, a *P. machaon* individual from the surroundings of Jerusalem was found to carry some *saharae* DNA – Fig. 2 of (Cassar *et al*., 2025); note also that individual W159 from the surroundings of Tel-Aviv may be part of the *saharae* mitochondrial clade, see Figure 3 of this work). While ecological isolation – *saharae* and *machaon* have different foodplants (Clarke & Larsen, 1986) (Pittaway *et al*., 1994) – is expected to affect the frequency and viability of hybrids, these should not experience diapause dysregulation, which makes it of particular interest to examine the genetic structure of the hybrid zone by genomic analyses.

Even more enigmatic than *saharae* is *ladakensis*. This taxon with short tails and broad submarginal lunules, which was described as a distinct species (Moore, 1884), inhabits rather dry valleys –the annual rainfall at Leh, the main city of Ladakh, is less than 100 mm – north of the main Himalayan range. It flies there from about 2800 m to above 4000 m and at the lower elevation, is encountered together with such butterflies as *Pieris krueperi* and *Pontia chloridice* (Tshikolovets, 2005), whose presence betrays affinities with the fauna of East Mediterranean mountains at moderate elevations. The population from Ladakh has attributes of a distinct species: its larva is markedly different from that of *P. machaon* (Figure 1 and (Hervillard *et al*., 2018)) and as reported in the present work, pronounced dysregulation of diapause in hybrid progenies is observed when crossing individuals with European *P. machaon*. Somewhat surprisingly, however, mtDNA sequences of specimens from Ladakh (except W106) cannot be separated from those of individuals from the Southern slopes of the Himalayan range (Figure 3). At the same time, two types of ITS2 sequences coexist in the *ladakensis* population, only one of which (type B) is clearly related to sequences from specimens – whether belonging to *machaon* or *everesti* – collected south of the main Himalayan range. Out of nine individuals examined (excluding the single specimen from Pamir), three had type A ITS2 sequences, two were of type B and the four remaining ones (two from Ladakh, one from Zanskar and one from South-West Tibet; Table 1) harbored a mixture of the two types (Figure 4). These data are compatible with frequencies of haplotype combinations being at equilibrium over the entire area, although obtaining statistical confirmation will require far more work.

One possibility suggested by this apparently free exchange on a broad geographic scale of two divergent subgroups of haplotypes with roughly similar frequencies is that just like *P. appalachiensis* in the *P. glaucus* group, *ladakensis* is in fact a hybrid species, which arose from crosses between males of a taxon with type A ITS2 sequences and females of *machaon asiatica*. A clue to the possible origin of the *ladakensis* type A ITS2 haplotypes is provided by Figure 5A, which shows that either this lineage diverged from *machaon* just after *everesti* and before *hippocrates*, or only slightly less likely, that its closest relatives are the type A sequences in *everesti*. Should *ladakensis* actually constitute a hybrid species, the question of how isolation from putative parental taxa, *machaon asiatica* and *everesti*, is achieved needs to be addressed. The westernmost known locality of *everesti*, where specimen W191 was encountered at elevations between 5000 and 5300 m, is 250 km distant from the places where *ladakensis* specimens W186 and W189 were collected, at only slightly lower elevations (Table 1) and nothing is known of possible contacts between the two taxa (recall, however, that crosses between *everesti* from Nyalam and *ladakensis* from Ladakh should result in about the same level of diapause dysregulation as observed in the laboratory between *everesti* and *hospiton*). In Ladakh and Zanskar, *ladakensis* is shielded from *machaon asiatica* by the main Himalayan range, the altitude of which does not go below 5000 m all the way to the Suru valley in Kashmir. Further west, however, no such barriers exist and based on adult wing patterns, *ladakensis* and *machaon* have been claimed either to ‘merge’ into one another (e.g. (Mackinnon & de Niceville, 1898)) or to coexist (as subspecies !) in Northern Pakistan (Tshikolovets, 2016) and the Western part of Ladakh (Tshikolovets, 2005). Only future investigations will be able to tell us whether such characters as larval morphology and the degree of diapause incompatibility with *machaon* change gradually or abruptly as one moves from the area of *ladakensis* into that of *machaon asiatica*.

Nearctic relatives of *P. machaon* are the source of a somewhat similar problem of species delimitation. As mtDNA sequences of *P. oregonia* and closely related taxa *bairdii* and *dodi* are very similar to those of West-palearctic *P. machaon* (Figure 3), these Nearctic entities are generally designated as subspecies of *P. machaon* in mtDNA-based phylogenies (e.g. (D’Ercole *et al*., 2021)). However, laboratory crosses by us (Results) revealed that female pupae with an *oregonia* father and a (French) *machaon* mother are perpetual nymphs (as shown by weighing pupae, incompatibility between the two parents is considerable indeed, see Figure 7). Are geographically intermediate populations intermediate as well in physiological incompatibility, or does there exist within the exceptionally vast distribution area of *P. machaon* – possibly in Western Canada, Alaska or Siberia – some narrow zone within which compatibility changes abruptly ?

More generally, it remains to be determined to what extent does position on the diapause incompatibility scale vary among populations of any particular species/taxon. Our own interspecific crosses, which involved *P. machaon* individuals of diverse populations from the North, Center and South of France, did not reveal significant differences that could have been attributed to the origin of the *machaon* parent when the extent of dysregulation of diapause in interspecific hybrids was compared. In Corsica as well, *hospiton* females from three distinct types of populations – low altitude, with larvae on *Ferula communis*; middle altitude, larvae on *Peucedanum paniculatum*; high altitude (1800 m), larvae on *Ruta corsica* (Aubert *et al*., 1997) – consistently gave rise, when mated to *machaon* males, to hybrid females that were unable to undergo pupal diapause (Table 2 and data not shown). However, these examples do not preclude the existence of significant differences between distantly located geographical areas. One may wonder in particular to what extent changes in voltinism affect diapause incompatibility. In the latter respect, it should be particularly interesting to investigate the situation in California, where *P. zelicaon* has become multiple-brooded in the course of two successive larval host-plant shifts, first from native Apiaceae to introduced fennel, then to *Citrus* (Shapiro & Masuda, 1980). Crossing a *P. zelicaon* male from Los Angeles with a *P. polyxenes* female was reported to result in plenty of adult males and not a single adult female (out of equal numbers of F1 hybrid pupae) (Oliver, 1969), just as had previously been found for *P. zelicaon gothica* (Ae, 1964). However, this was back in the 1960’s and additional crosses involving individuals from different populations appear highly desirable.

## CONCLUSION

In this work, we posit the existence of a quantitative physiological trait, ‘*dDiap*’, whose primary role could be to control the intensity of diapause. Although we ignore at present how to assess this trait directly in individual organisms, its value or, rather, the difference between its values in two individuals of opposite sex can be precisely determined by crossing them and comparing male and female pupal weights in their F1 progeny (Figure 7). Large differences in parental *dDiap* values result in severe, although female-limited, F1 post-zygotic incompatibility, in the form of inability either to enter or terminate pupal diapause. Moreover, incompatibility reoccurs in part of the progeny of hybrid males, resulting in marked overall reproductive failure.

Our laboratory crosses failed to provide evidence of significant variation of *dDiap* among individuals within the populations we could examine. Nor did our preliminary investigations find clear differences in *dDiap* between populations of the same species within a territory of limited geographic dimensions and reasonably uniform climate. On the other hand, given that *dDiap* is bound to be under strong selection by ecological conditions, we do expect it to vary within widely distributed species, especially when differences in voltinism are involved. For instance, the marked difference in *dDiap* we observed between French *P. machaon* and *P. (machaon) oregonia* from Northwestern United States cannot be taken as proof that these two taxa belong to different species until it has been determined whether variation of *dDiap* within the unusually vast range traditionally attributed to *P. machaon* is continuous or discontinuous.

How ecologically driven, pre-existing differences in *dDiap* may impact processes of speciation and make their outcome predictable to some extent is illustrated by situations in which mutually fertile members of the *machaon* complex endowed with quite different *dDiap* values happen to coexist in sympatry – *P. machaon* and *P. hospiton* in Corsica and Sardinia – or parapatry – *machaon* and *everesti* at the border between Nepal and China or *zelicaon* and *polyxenes* at the contact between Rocky Mountains and the Great Plains of North America. The former situation possibly results from *machaon* (which in Corsica uses preferentially fennel as larval foodplant) having taken opportunity of the onset of agriculture in the Tyrrhenian islands to invade *hospiton*’s range: differences in *dDiap* between the colonizing and resident taxa could have helped preserve the genetic integrity of the latter by providing potential coupling with the barrier effect resulting from the use of different foodplants (female ovoposition preferences in two other members of the *machaon* complex were shown to be determined primarily by their Z chromosome (Thompson, 1988b)). Coincidence of multiple barrier effects could explain as well the stability of parapatric contacts, despite hybridization, when differences in *dDiap* coincide with steep ecological gradients, as at the Nepalese-Chinese border or at the foot of the Rocky Mountains Front Range. Inversely, however, crosses between taxa with very different climatic and ecological requirements and, accordingly, markedly different *dDiap* values could, in case they succeeded in giving rise to a genetically stable hybrid population with an intermediate *dDiap*, provide this new hybrid entity with the ability to colonize biotopes with somehow intermediate characteristics: as suggested by our DNA sequence data, this is possibly how *P. ladakensis* came into existence. Comparative genomic description of the populations in contact on the one hand, and in-depth genomic analyses of laboratory backcrosses on the other, may make it possible to untangle the contributions of *dDiap* and a number of other traits to these and other processes of speciation and hybridization that shaped the *machaon* complex.

Finally, how ancient and widespread are diapause-linked postzygotic incompatibilities of the type we have been investigating in the *P. machaon* complex? As already mentioned, studies that rest on insufficiently controlled or else invariant rearing conditions may miss not only the inability of female pupae to enter diapause, but inability to terminate diapause as well, when the latter phenotype is conditional or dead pupae are not sexed. Still, despite the likelihood of perturbations of diapause having frequently gone unnoticed, interspecific crosses were found to result in perpetual nymphs not just in two subdivisions of the vast *Papilio* genus, but in at least two moth families as well (see Discussion), and we regard this as evidence that the regulatory network whose perturbation underlies hybrid dysregulation of diapause predates the origin of butterflies. In this respect, it is also worth noting that a recent phylogenetic analysis by (Halali *et al*., 2024) suggests that pupal diapause may have been the ancestral state in butterflies. Furthermore, it should be recalled that pupal diapause may not be the only form of diapause that is subject to dysregulation in hybrid progenies; in fact, (Oliver, 1983) observed that in some reciprocal interspecific crosses which result in inverse perturbations of larval developmental rates, larval diapause may be affected as well.

## Supporting information

Machaon complex phylogeny_Supplemmentary file

## ACKNOWLEDGEMENTS

We thank H.G. Allcard, F. Carbonell, Y. Delmas, M. Grinell, J.-F. Hervillard, G. Labeyrie, T. Michel, J. Pagès, B. Turlin and especially J.-C. Weiss for supplying us with specimens and are particularly grateful to A. Cotton for communication of unpublished data. M. Joron, L. Després and, above all, M. Elias provided useful comments on a draft version of the manuscript. Experimental data were generated by F.M. and L.L. at the former Centre de Génétique Moléculaire du C.N.R.S. (Gif-sur-Yvette, France) and by H.D. at the Université de Provence (Marseille, France).

## DATA AVAILABILITY

Mitochondrial and ITS2 sequences have been submitted to GenBank. Photographs of adult specimens from which DNA was extracted will be made available in the Zenodo repository (https://doi.org/10.5281/zenodo.20368257).

## SUPPLEMENTARY INFORMATION

Supplementary Information consists of Extended Methods, Extended Discussion, Table S1 and Figures S1 to S10.

