## Supplementary material for "Dysregulation of pupal diapause in hybrid progenies, introgression and species delimitation within and beyond the Old World Swallowtail (*Papilio machaon* Linnaeus) butterfly complex": Machaon complex phylogeny_Supplemmentary file

#### EXTENDED METHODS

#### Laboratory breeding

Unless otherwise stated, our *P. machaon* stocks originated from females collected in the surroundings of Paris, in Bourgogne or at moderate elevations in the mountains of South-Eastern France. *Papilio hippocrates* pupae were bought from Far East Entomological Supplies. *Papilio.* (*machaon*) *saharae* eggs were obtained from two females collected by H.D. at Tizi n'Tiniffift (Morocco). *Papilio machaon mauretanicus* was from eggs and larvae collected some 100 km South-West of Marrakech (Morocco) by F. Diemer (Pittaway *et al.*, 1994). *Papilio ladakensis* eggs were obtained from two females collected in Ladakh. *Papilio hospiton* was from Corsica (Aubert *et al.*, 1997). *Papilio oregonia* was obtained from pupae sent to H.D. from Celilo (Wasco Co., Oregon, USA). *Papilio everesti* larvae and adults were collected between Kodari (Nepal) and Nyalam (Tibet, China). Except for some spontaneous pairings in 1 $m^3$ cages between *P. machaon* virgin females and *P. hospiton* males (and vice versa; Aubert *et al.*, 1997), crosses were carried out by hand-pairing (Clarke & Sheppard, 1956a).

*Papiilio machaon* caterpillars were fed either carrot (*Daucus carota*), fennel (*Foeniculum vulgare*), parsnip (*Pastinaca sativa*) or rue (*Ruta graveolens*), depending on their mother's egg-laying preferences and larval foodplant availability. *Papilio* (*machaon*) *saharae* larvae from Tizi n'Tiniffift did not eat carrot, but grew rapidly on rue. *Papilio hospiton* was bred successfully on parsnip. *Papilio ladakensis* caterpillars were fed either fennel (the whole plant) or parsnip (only flower buds and fruits; even so, growth was much slower than on fennel, although mature larvae reached about the same final weight). *Papilio everesti* larvae were bred on parsnip (both flowering parts and leaves were consumed). For caterpillars issued from F1 crosses either parsnip, if available, or one of the parental foodplants was used, whereas for F2 larvae care was taken to use only a foodplant accepted by both parental taxa.

Under 'long-nights' conditions, illumination was for 8 to 10 hours, whereas 'short-nights' corresponded to 16 hours daylight. Somewhat unexpectedly given its high altitude environment, we found that diapause is facultative in *ladakensis*: contrary to caterpillars raised under long-nights conditions, all those that had experienced short nights developed into non-diapausing pupae. By contrast, all *everesti* pupae (except for a single female) were found to diapause when larvae were raised under short-nights conditions (in both cases, synchronous machaon broods were used as controls). Some female larvae issued from (male) *hospiton* x (female) *machaon* crosses were prevented from entering diapause at the pupal stage and becoming perpetual nymphs by raising them under short-nights conditions and maintaining temperature above 20 °C (Aubert *et al.*, 1997). Pupae were sexed and weighted three days after metamorphosis.

Pupae that did not resume development after 10 months at 2°C followed by two months at 20°C under short-nights conditions were put back at 2°C for some six additional months before being brought back to 20°C. As this second exposure to cold conditions did not, as a rule, trigger development, those pupae were then scored as perpetual nymphs. Some female perpetual nymphs obtained from male *P. hospiton* x female *P. machaon* and male *P. ladakensis* x female *P. machaon* crosses were injected with ecdysone (Clarke & Willig, 1997) so as to check that adults were viable.

The backcrosses used to generate Figure 8 involved *ladakensis* x *machaon* male hybrids on the one hand, and nine *machaon* females on the other (seven of these were sisters and the remaining two came from the same local stock, collected close to Paris). The *ladakensis* x *machaon* male hybrids had been obtained by crossing *ladakensis* individuals from our stock with *machaon* females, again from the same local stock.

#### DNA extraction, PCR amplification and sequencing

DNA extraction was carried out either as in Aubert *et al.* (1997) or from a single adult leg with a Macherey-Nagel Nucleo Spin Tissue kit. Whenever DNA of the expected length could not be generated from ITS2 or one of the three mitochondrial segments of interest with our routine PCR protocol and oligonucleotides (Table S1), subsequent attempts generally proved successful when alternative, substitute primers were used (see Table S1). In the case of extracts W160, W382, W469 and W475, however, not only did initial PCR amplification fail for more than one gene, but PCR products could be obtained only when the length of DNA to be amplified was reduced. As those four extracts were from dried individuals which had been kept in that state at room temperature for 12 to 28 years, a likely explanation is that these samples contained partially degraded DNA.

#### Alignment of ITS2 sequences

As widely advocated for insightful comparative sequence analysis (e.g. (Michel *et al.*, 2000) and references therein), alignment of ITS2 sequences was optimized manually, by taking simultaneously into account both the sequence and the potential secondary structure of RNA transcripts of the ITS2 segment. For RNA structure prediction by minimization of calculated free energy, we used RNAstructure 6.3 (Reuter & Mathews, 2010). For all taxa analyzed, the predicted ITS2 minimal energy structure consisted of four elongated stems (I to IV in Figure) radiating from a central 'wheel'. This general architecture happens to coincide with the secondary structure derived by comparative analysis of eukaryotic ITS2 sequences, which is remarkably well conserved by evolution (Schultz *et al.*, 2005). Sequences in the basal sections of stems I, II and IV are quite conserved and the same is true of the distal section of stem III (Fig. S1). These are precisely the segments that have been shown in yeast to bind proteins and harbor sites that are essential to pre-ribosomal RNA processing: removal of ITS2 is initiated by endonucleolytic cleavage at the 'C2' site, which is located close to the tip of stem III (Gasse *et al.*, 2015), and requires binding of several additional proteins by the basal parts of that RNA (Biedka *et al.*, 2018). By contrast, comparison of the distribution of ITS2 nucleotide changes within Palearctic populations of *P. machaon* and its close allies (Fig.S1) on the one hand, and the entire *machaon* complex (Fig. S2) on the other, shows that accumulation of mutations can be remarkably rapid in a number of segments. Among these, the middle part of stem III stands out for its ability to tolerate major insertions and deletions (Fig. S2), which only have in common that they do not seem to compromise its potential to fold into a reasonably compact structure that does not leave too many nucleotides exposed in single-stranded segments, as should be expected if that section of the ITS2 spacer happened to lie naked on the outer surface of pre-ribosomal RNA complexes. If that is indeed the case, the function of the middle part of stem dIII could merely be to ensure covalent continuity, together with sufficient conformational flexibility, so as to allow the conserved distal section of that stem to be brought back into proximity of other conserved sections of the ITS2 RNA, just as suggested by our secondary structure drawings (Figs. S1 and S2). However speculative this discussion might sound, it is worth noting that there does exist in yeast pre-ribosomal RNA complexes a protein, Cic1, that binds simultaneously stem I and the distal section of stem III (Biedka *et al.*, 2018).

By comparing potential secondary structures, 98 sequences from 44 Palearctic individuals (other than *hospiton*) and the closely related *P. machaon aliaska* W462 and *P. brevicauda* W382 could be aligned over the entire length of the ITS2 segment, resulting in 71 distinct haplotypes (the ratio of haplotypes over individuals would have been even higher had not the distal section of stem I – with the sequence UAUAUAU in the reference, NW014509617 *machaon* genome – been excluded from this and subsequent analyses, due to extreme sequence instability, even within single individuals; that instability results from the replacement, in about half of the dataset – 50 sequences out of 99 – of a CUCAUAG capping sequence by CUAUAUAUAUAG or longer repetitive sequences). These 71 haplotype sequences of interest were found to differ from one another by a limited number of point substitutions and small indels – of no more than 4 nt, except for a 33 nt deletion in haplotype H127B (Fig. S2). Out of 56 detectable mutational events, 33 consisted in substitutions and 28 in insertions/deletions (when affected positions were contiguous, changes were scored as single events). All these indels are associated with short sequence repeats, thus pointing to replication slippage (Levinson & Gutman, 1987) as an obvious, and particularly efficient, causal mechanism (recall that RNAs that are not destined to be translated should tolerate many insertions and deletions as long as these neither interfere with the folding processes that lead to the final, functional structures, nor compromise the ability of key structures and sequences to bind their protein and RNA partners).

### EXTENDED DICUSSION

#### Dysregulation of pupal diapause in the progeny of crosses involving Nearctic relatives of *P. machaon*

Information about dysregulation of pupal diapause in crosses involving Nearctic relatives of *P. machaon* is scattered in a number of reports, a majority of which are more than 40 years old. Several experimenters complain about larvae being affected by diseases, so that numbers of adults are often very small. One additional difficulty is that reciprocal crosses were not always analyzed separately. For instance, (Tyler *et al.*, 1994) when quoting Heitzman (1973; pers. comm.) state that 'twenty *joanae* x *polyxenes* crosses (both ways)' gave '319 male and 285 female pupae from which 299 males but only 75 females emerged, the latter mostly much later'. Similarly, in Table 1 of (Ae, 1979), which summarizes results of a large number of interspecific crosses within the vast *Papilio* genus, the outcomes of reciprocal crosses are pooled together, so that for *polyxenes* and *gothica*, for instance, it is only stated that four crosses yielded 82 males and 20 females. In this particular case, however, the author added in the main text that 'the findings may be different when a reciprocal cross is carried out. Thus in a cross between *Papilio polyxenes* female and *P. bairdii brucei* male (= *P. gothica*), the writer obtained seventy-eight males only, but in the reciprocal cross he obtained nine males and seventeen females (Ae, 1964).' Next comes the question of the actual identity of the male parent. '*P. gothica*', which was described as a novel species by (Remington, 1968), was concluded to be 'only a minor high mountain ecotype of *P. zelicaon*' by (Clarke & Sheppard, 1970) and most subsequent authors. Still, the fact is that the wing patterns of *gothica* are somewhat similar to those of *P. bairdii brucei* (definitely another species) and the rather poor quality of photographs in Ae's works (Ae, 1964) (Ae, 1979) does not make it possible to decide which taxon was actually used. Fortunately, an essential piece of information is to be found in Table 1 of (Remington, 1968), in which 'Cross P1-28', between a *polyxenes* female and a *gothica* male, with a progeny consisting of 23 males, no female and a single intersex individual, is stated to come from Ae's work; Ae and Remington collaborated and Cross P1-28 is mentioned indeed in Table 3 of (Ae, 1964): it did yield a single gynandromorph and no adult female. However, the number of male offsprings was 78, not 23, and the author added in the Text that 'Forty-five pupae, which were comparatively large and thus presumably females, did not emerge by the end of the fall. These pupae were refrigerated till next spring and again kept in the rearing room from spring to fall. But no indication of emergence appeared'. Moreover, a reciprocal cross (also in Table 3) 'which larvae were reared in July at the Rocky Mountain Biological Laboratory, produced only females in the same year and produced only males in the next spring': this is cross 'B2', which yielded a total of 9 males and 17 females. In short, crosses between single-brooded *P. zelicaon* from Colorado and multiple-brooded P. *polyxenes* yield (possibly oversize) female perpetual nymphs when the male parent is *zelicaon*, whereas the female (not male) progeny is unable to enter diapause when the father is *polyxenes*: this is no different from what we observed when crossing *P. hospiton*, *P. ladakensis* or *P. oregonia* on the one hand, with *P. machaon* on the other.

Confirmation that *P. zelicaon* is incompatible with *P. polyxenes* was provided by (Oliver, 1969), whose *zelicaon* stock was from Los Angeles (California), where this species is multiple-brooded. Crosses between *zelicaon* males and *polyxenes* females resulted in 36 male pupae, from which 31 adults emerged, and 36 female pupae, from which not a single adult emerged. As for *polyxenes*, many crosses have been carried out with either males or females of European *P. machao*n e.g. (Clarke & Sheppard, 1953) (Clarke & Sheppard, 1955a) (Oliver, 1969) (Clarke *et al.*, 1977) (Blanchard & Descimon, 1988) and no significant distortion of sex ratio in hybrid progenies was reported. Therefore, *polyxenes* must lie close to *machaon* on the scale in Figure 7a and consequently, *zelicaon* should fall somewhere near *hospiton*, *ladakensis* and *oregonia*. Proximity of *zelicaon* and *oregonia* on that scale is further supported by (Thompson, 1988a) obtaining large numbers of *zelicaon* x *oregonia* F1 adult females from overwintering pupae independently of whether the parents were a male *oregonia* and female *zelicaon* or vice versa. Finally, even though only limited numbers of pupae were involved, it is worth mentioning that observations reported by (Clarke & Sheppard, 1953) – namely, that female pupae from a cross between male *zelicaon* (from California) and female *machaon* did not hatch, whereas in the reciprocal cross two out of three females emerged before males – are fully consistent with what precedes.

Information about other taxa is limited. Several crosses that involved *P. brevicauda* have been reported, but numbers of offsprings were very low, except for a cross between male *polyxenes* and female *brevicauda* that yielded seven males and eight females, with, interestingly, females emerging first (Clarke & Sheppard, 1953)). In another paper (Clarke & Sheppard, 1955a), these authors also wrote that 'It will be noticed [ ] that when some pupae remain from a brood long after the others have emerged, *P. brevicauda* is nearly always one of the ancestors'. In yet another series of experiments, female hybrids from crosses between *brevicauda* males and *machaon* females (the latter from Austria and Luxembourg) entered diapause (and emerged after overwintering), whereas their brothers underwent direct development (Adam Cotton, personal communication). Altogether, these data suggest that there exists some moderate incompatibility between brevicauda and *polyxenes*/*machaon*. Thus, single-brooded *brevicauda* should fall in between *machaon*/*polyxenes* on the one hand, and *hospiton* on the other.

It should finally be mentioned that a cross between a male *machaon britannicus* and a female *machaon hippocrates* was found to result in a dearth of hybrid females and their late emergence (Clarke & Sheppard, 1956b). Since crosses between multiple-brooded *hippocrates* and European *machaon* by us (data not shown) and others (e.g. (Ae, 1964)) did not reveal significant sex-linked biases in pupal development and adult eclosion, the 'culprit' is most likely to be single-brooded *britannicus*, which should therefore be placed somewhere next to *brevicauda*.

**TABLE S1** Oligonucleotide primers used in this work

| gene | name | orientation | sequence 5′ to 3′ | primer function |
|---|---|---|---|---|
| ND1 | ND1-94 | Forward | CGTCAAGCTTTAGGTTATATTCAAATTCG | |
| | ND1rev619 | Reverse | ATCAAAAGGTGTTCGATTAGTTTC | |
| | ND1-6632 | F | TGATTATTAATTCCTTATTATTTTAA | substitute(1) |
| | ND1-6631 | R | TAATCTAACTTCATATGAAATCGTTTG | substitute |
| COX1 | CO-21686 | F | ATTCAACAAATCATAAAGATATTGG | |
| | CO-2192 | R | CCCGGTAAAATTAAAATATAAACTTC | |
| | CO-362 | F | AGAAGAATYGTAGAAAATGGAGCAGGAAC | substitute |
| | CO-346 | R | AGTTCCTGCTCCATTTTCTA | substitute |
| COX2 | CO-6281 | F | TTCTAATATGGCAGATTATATGTAATGGATTTAA | |
| | CO-3782 | R | GAGACCATTACTTGCTTTCAGTCATCT | |
| | CO2midS | F | AAATCAATTGGTCATCAATGATA | substitute |
| | CO2midR | R | ATAGTAACTACTGGTTAACTAAA | substitute |
| ITS2 | ITS3B | F | GGTCGATGAAGAACGCAGTTA | |
| | ITS4B | R | TTTCCTCCGCTTACTAATATGCTTA | |
| | ITS3C | F | TGGACGGTGGATCACTTGG | substitute |
| | ITS6 | R | AACCTCCCGAACACCACA | substitute |
| | ITS7 | F | TAGCGAACAATTGACGGTT | substitute |
| | ITSmidS | F | CGACTCTTCGACGTAAAAAATC | substitute |
| | ITSmidR | R | GATTTTTTACGTCGAAGAGTCG | substitute |
| | ITSmac | F | TAAAAATCACACTGTTCACTCA | haplotype-selective |
| | ITSsah | F | TAAAAATCACACTGTTCACTA | haplotype-selective |
| | ITSZanS | F | AAATATAACCTCTCAGACACAC | haplotype-selective |
| | ITSLadS | F | AAATATAACCTCTCAGACACAT | haplotype-selective |
| | ITSZanR | R | CGCTCTTCGGACGTCGT | haplotype-selective |
| | ITSLadR | R | CGCTCTTCGGACGTCGA | haplotype-selective |
| | ITSmacR | R | TGCCGTGTTTTCACACAC | haplotype-selective |
| | ITShippoR | R | TGCCGTGTTTTCACACAG | haplotype-selective |

(1) Substitute primers for refractory samples that failed to yield the expected PCR product on an initial DNA amplification attempt.

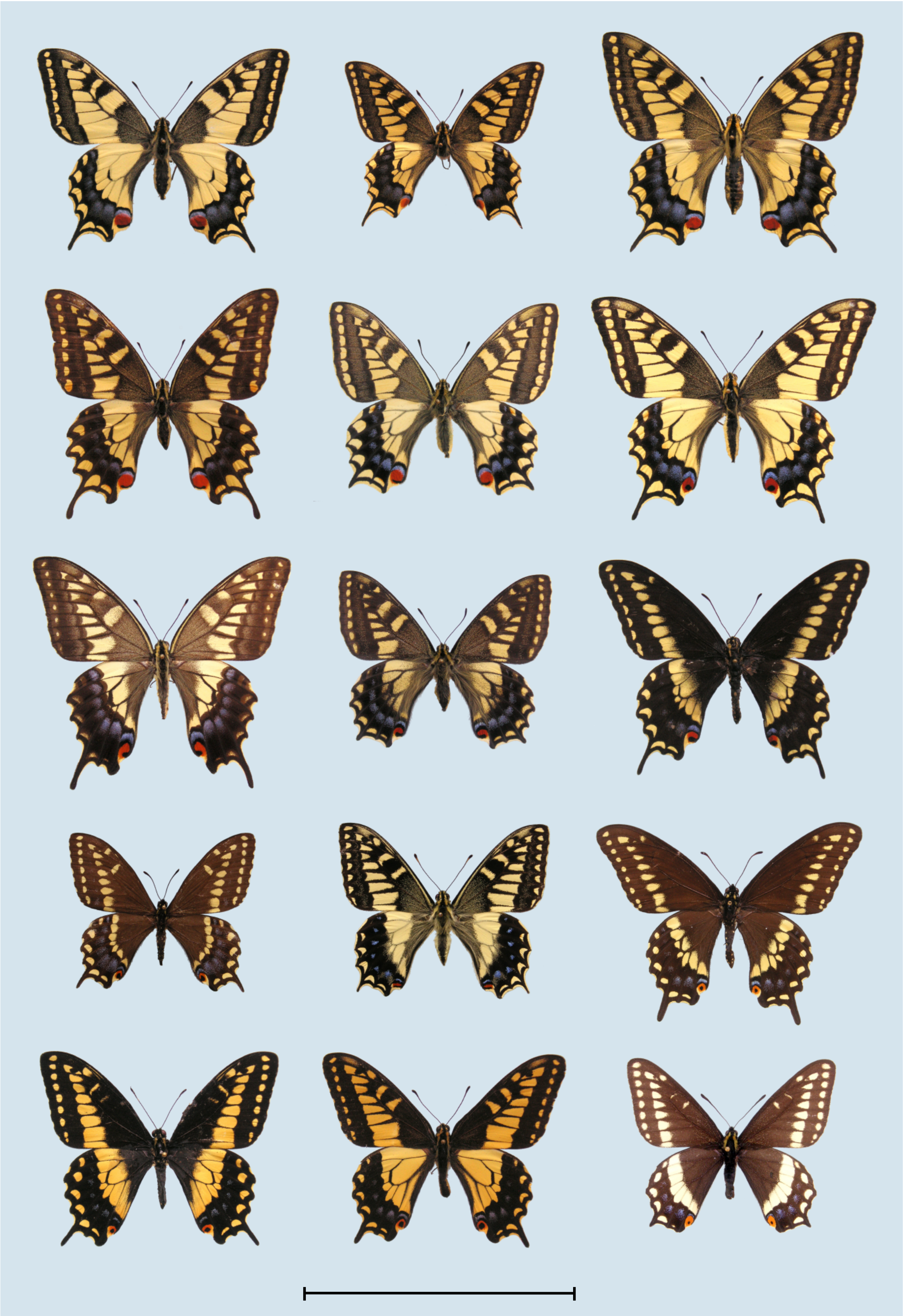

**Figure S1.** Adults of the *Papilio machaon* complex. Top row, from left to right: *P. machaon gorganus* ♀, France, Yvelines; *P. machaon mauretanica* ♂ (W181), Algeria, Algiers; *P. machaon saharae* ♀ (W192), Morocco, Tizi n'Tiniffift. 2nd row: *P. machaon asiatica* ♀, Nepal, Kodari; *P. ladakensis* ♀, India, Ladakh, Leh; *P. oregonia* ♀ (W472), USA, Oregon, Fulton Canyon. 3rd row: *P. hippocrates* ♀, Japan, Kyoto; *P. everesti* ♀, China, Tibet, Nyalam; *P. bairdii* ♂ (black form, W502), USA, Arizona, White Mountains. 4th row: *P. brevicauda* ♂ (W382), Canada, Newfoundland; *P. hospiton* ♂, France, Corsica, Restonica; *P. polyxenes asterius* ♂, USA, Illinois. 5th row: *P. polyxenes stabilis* ♂ (W491), Costa Rica; *P. zelicaon* ♂, USA, California, Long Beach; *P.indra* ♂, USA, California. Length of bar at bottom is 80 mm.

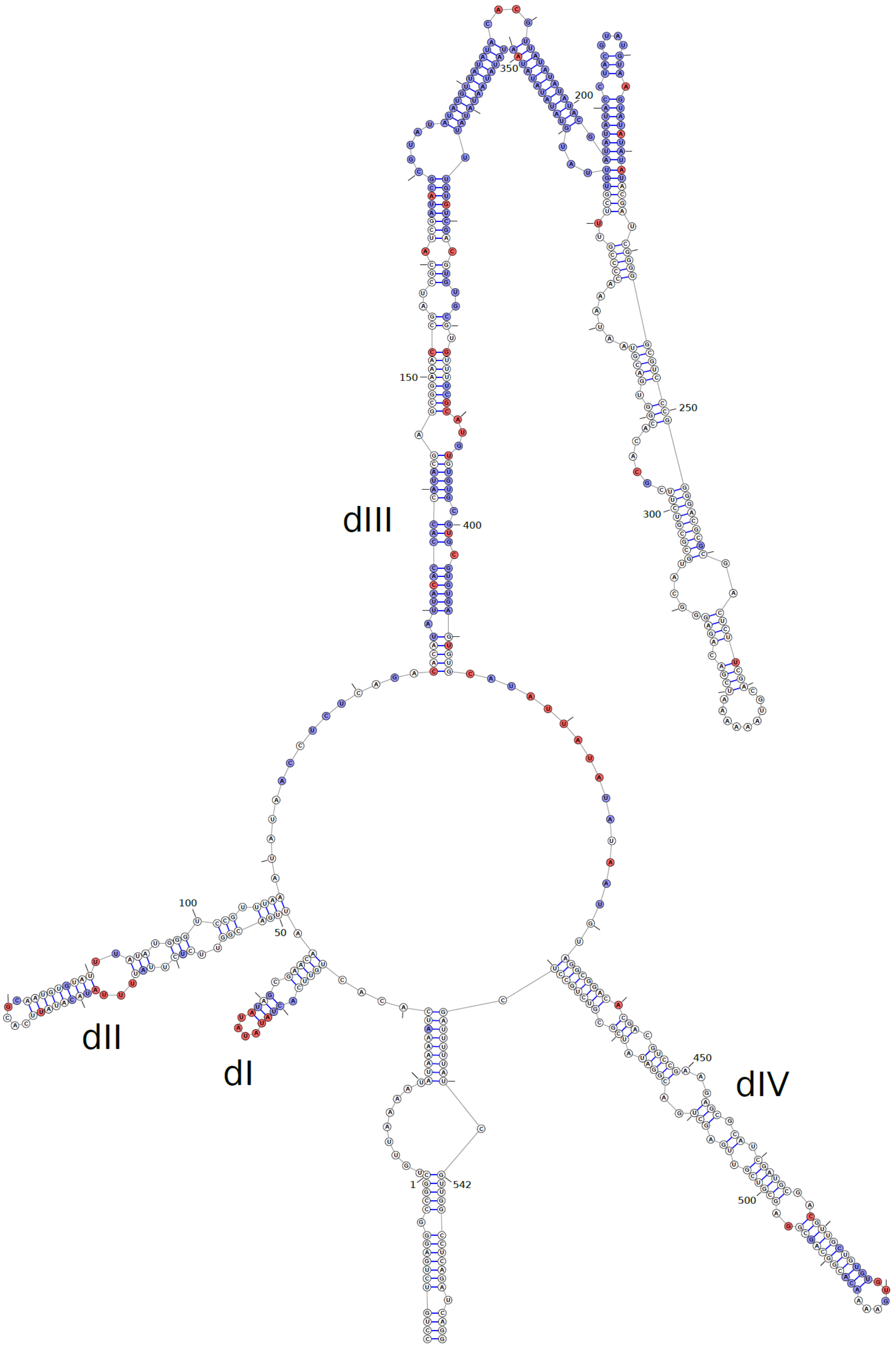


**Figure S2.** Predicted secondary-structure model of ITS2 RNA for the most common *P. machaon* ITS2 haplotype (see Fig. 4). Sites colored in red are variable among sequenced Old World individuals (*P. machaon aliaska* and *P. brevicauda* were included, while *P. hospiton* and also the *P. machaon rathjensi* H127B haplotype, with a 33 nt deletion in stem III, were not taken into account). Sites in blue were found to have been affected by at least one mutational event in our entire dataset.

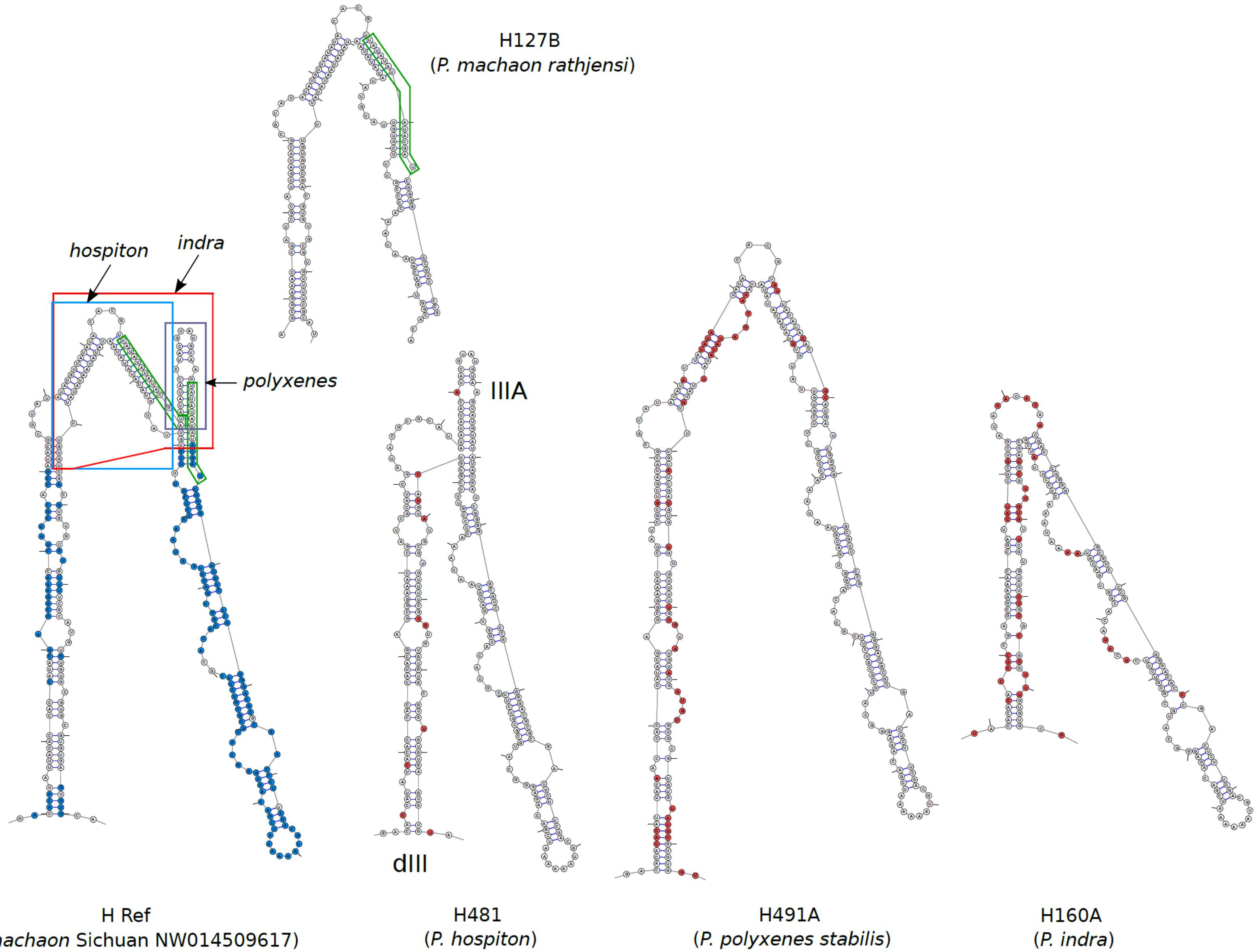


**Figure S3.** Predicted ITS2 stem III structures for five divergent haplotypes. In the machaon HRef haplotype structure, nucleotides in blue are conserved throughout our dataset. Red, blue and black boxes show nucleotides missing in the *indra* H160A, *hospiton* H481 and *polyxenes* H491A haplotypes. Green boxes indicate two 14-nt identical sequences whose recombination generated a 33-nt deletion in the H127B haplotype. Nucleotides in the dIII stems of *P. hospiton*, *P. polyxenes* and *P. indra* that differ from their counterparts in *P. machaon* are circled in red.

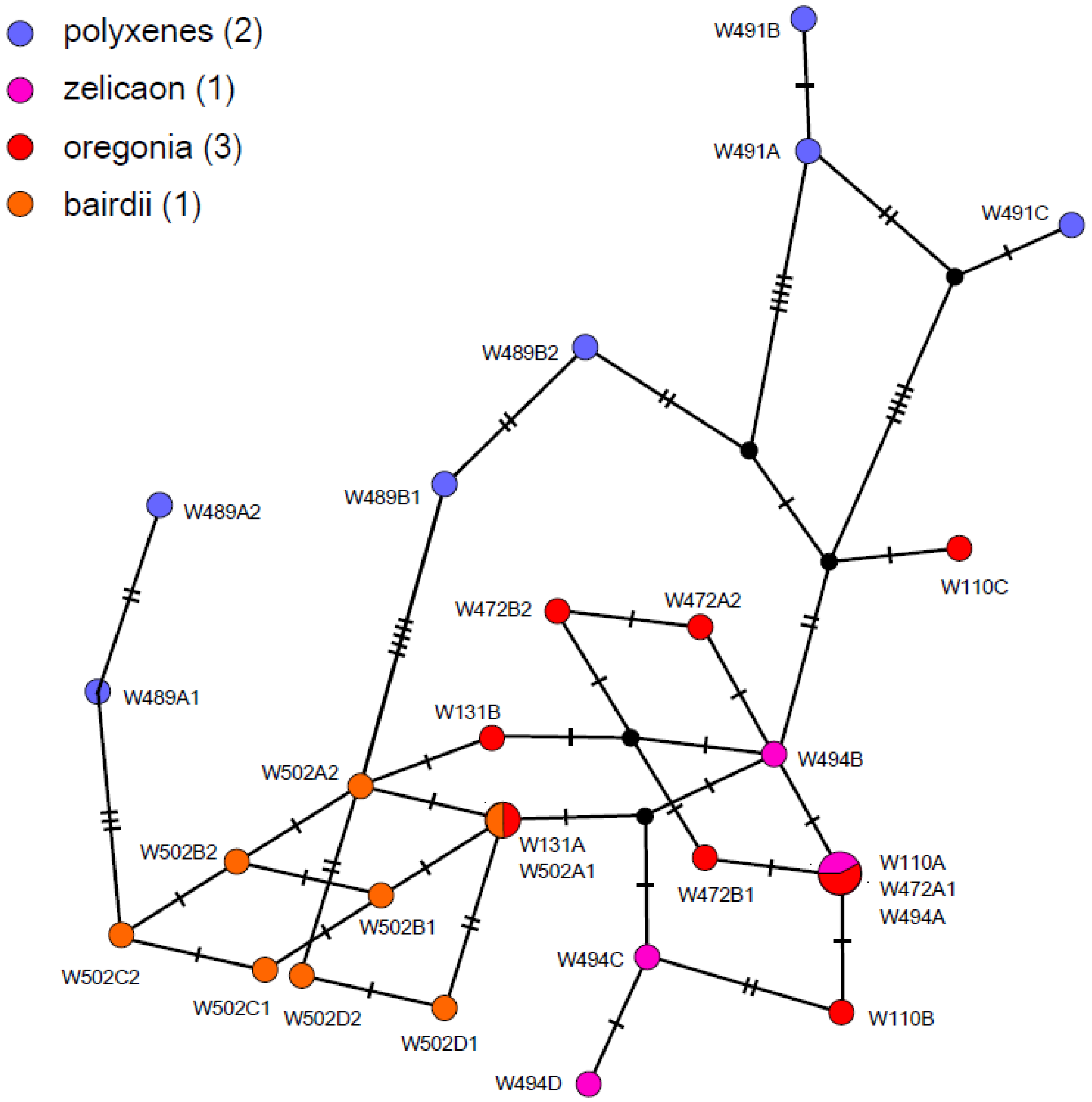


**Figure S4.** TCS network of ITS2 haplotypes from seven North American individuals. For coding of node size and color and node-connecting lines see legend to Fig. 4.

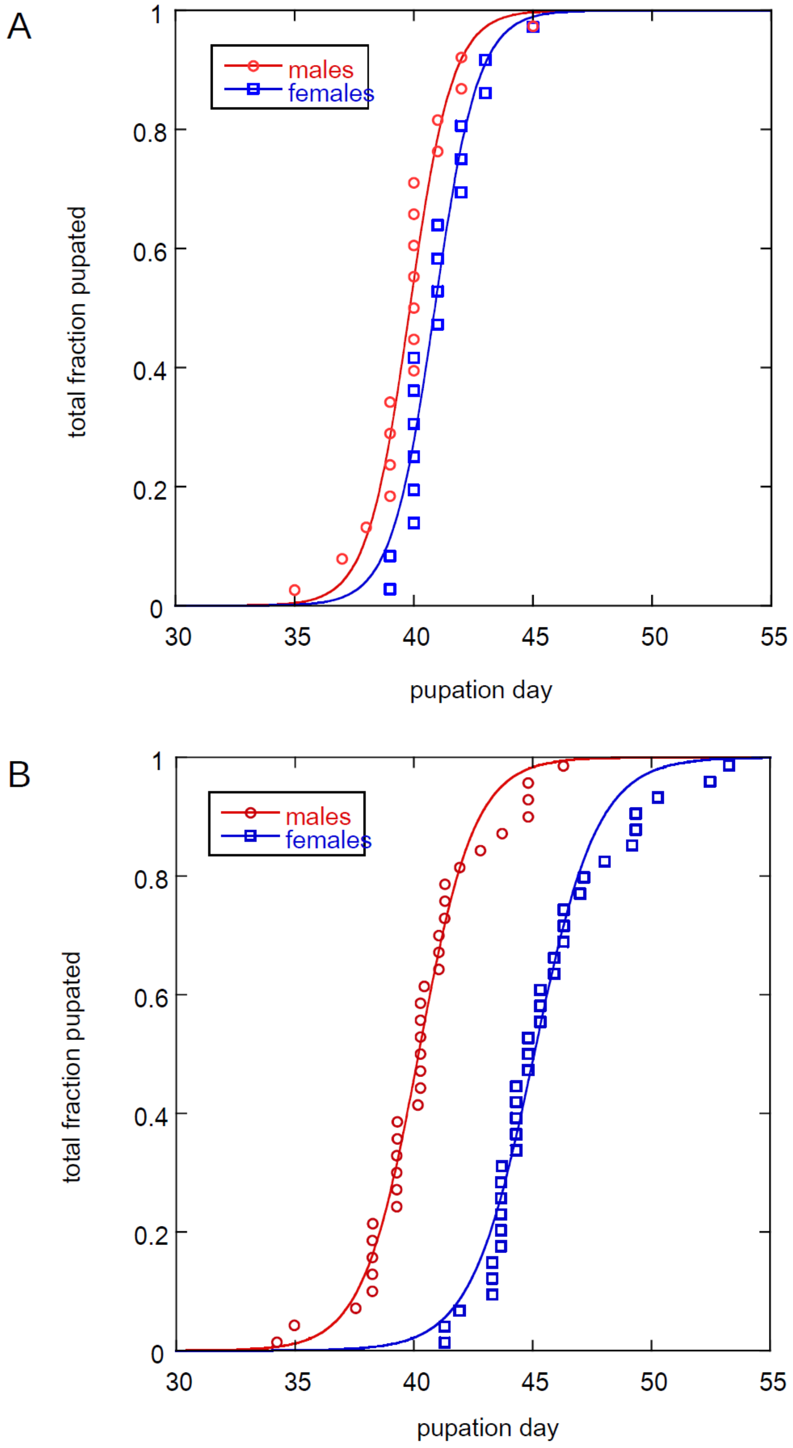


**Figure S5.** Comparison of pupation time for female perpetual nymphs and their male siblings. (A) *P. machaon* control. Larvae from the same egg-laying session (about 4-hour duration) were reared together under long nights at 20 °C. Abscissa, days after egg hatching; data were fitted to the equation of a sigmoidal curve, $1/(1+\exp(-a*(x-m)))$, where m is the mean pupation date (39.80 ± 0.08 days for males, 40.90 ± 0.09 for females). Mean pupal weights were 0.942 ± 0.017 g for males, 1.071 ± 0.017 g for females, resulting in a weight ratio of 1.14. (B) Eggs were obtained from three *P. machaon* females that had been paired with *P. ladakensis* males. Larvae that shared the same mother and originated from the same egg-laying session (about 4-hour duration) were reared together under long nights at 20 °C. Abscissa and fitting as in panel (A). The sex and weight of pupae was determined three days after metamorphosis and data were pooled for all three mothers, revealing a 4.9-day lag in pupation of females, compared with males (mean pupation date was 40.20 ± 0.05 days for males, 45.07 ± 0.07 for females). Mean pupal weights were 0.795 ± 0.003 g for males, 1.215 ± 0.006 g for females, resulting in a weight ratio of 1.53.

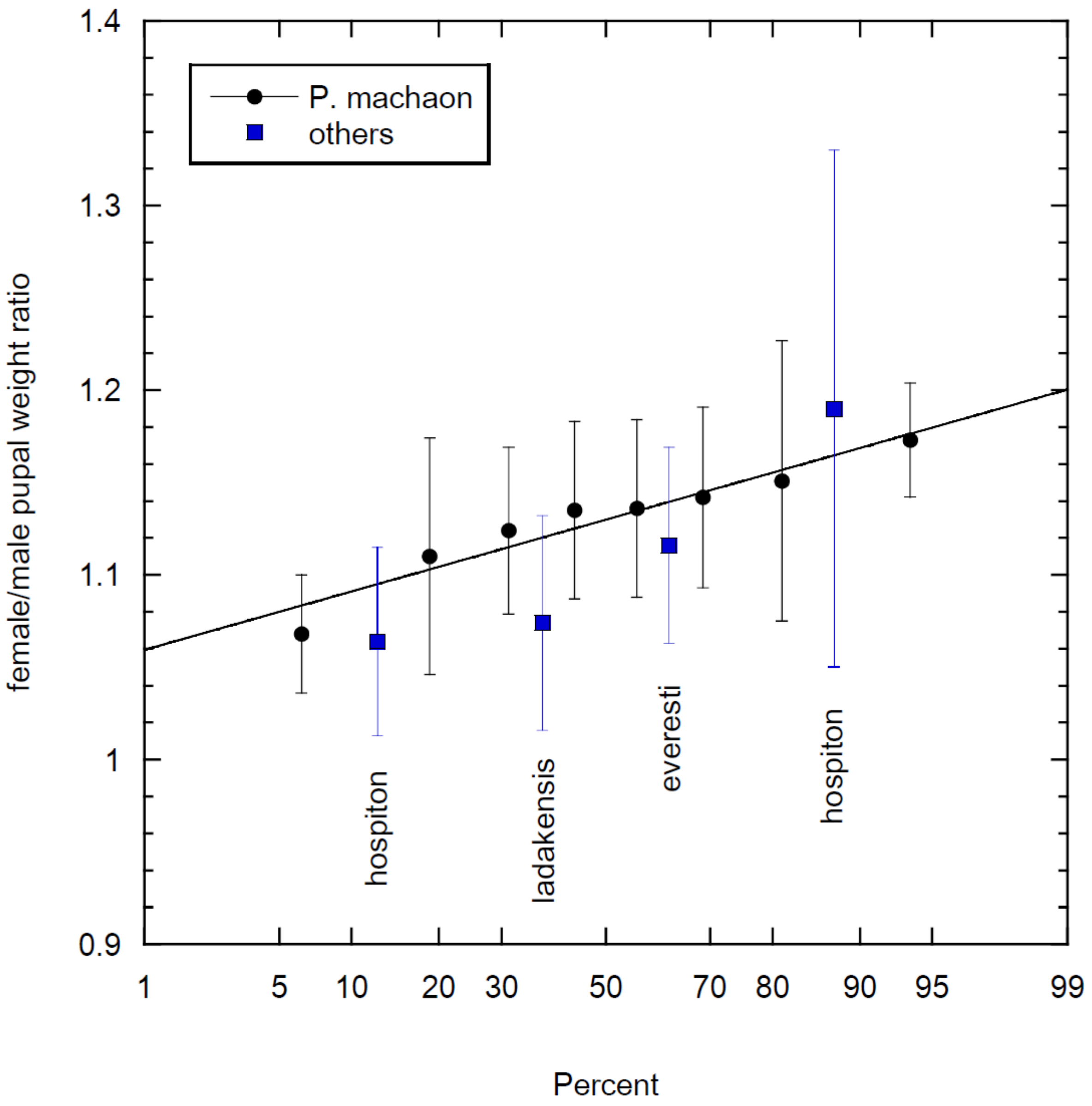


**Figure S6.** Distribution of female over male pupal weight ratios in intraspecific crosses. Bars indicate standard errors. Average pupal weight ratio and its standard error for 8 unrelated *P. machaon* broods (228 males, 225 females) was 1.13 ± 0.01.

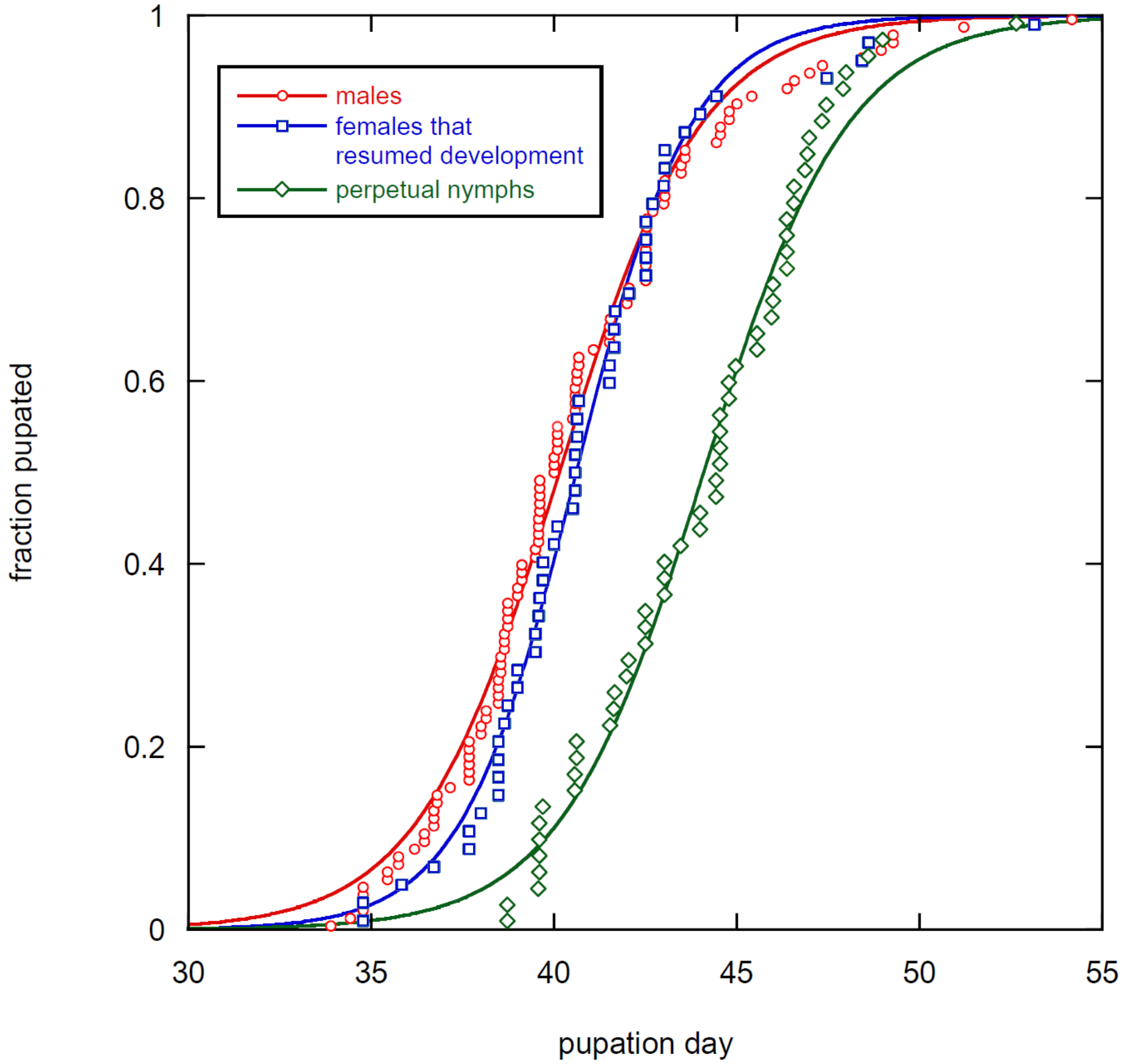


**Figure S7.** Distribution of pupation times in the progeny of backcrosses of (*machaon* x *ladakensis*) male hybrids to *machaon* females. Same experiment as in Figure 8, except that individuals whose pupation time was unknown or which had not been reared at 20°C were discarded. Mean pupation dates were 40.15 ± 0.03 days for males, 40.61 ± 0.03 days for females that resumed development, 44.10 ± 0.05 days for perpetual nymphs.

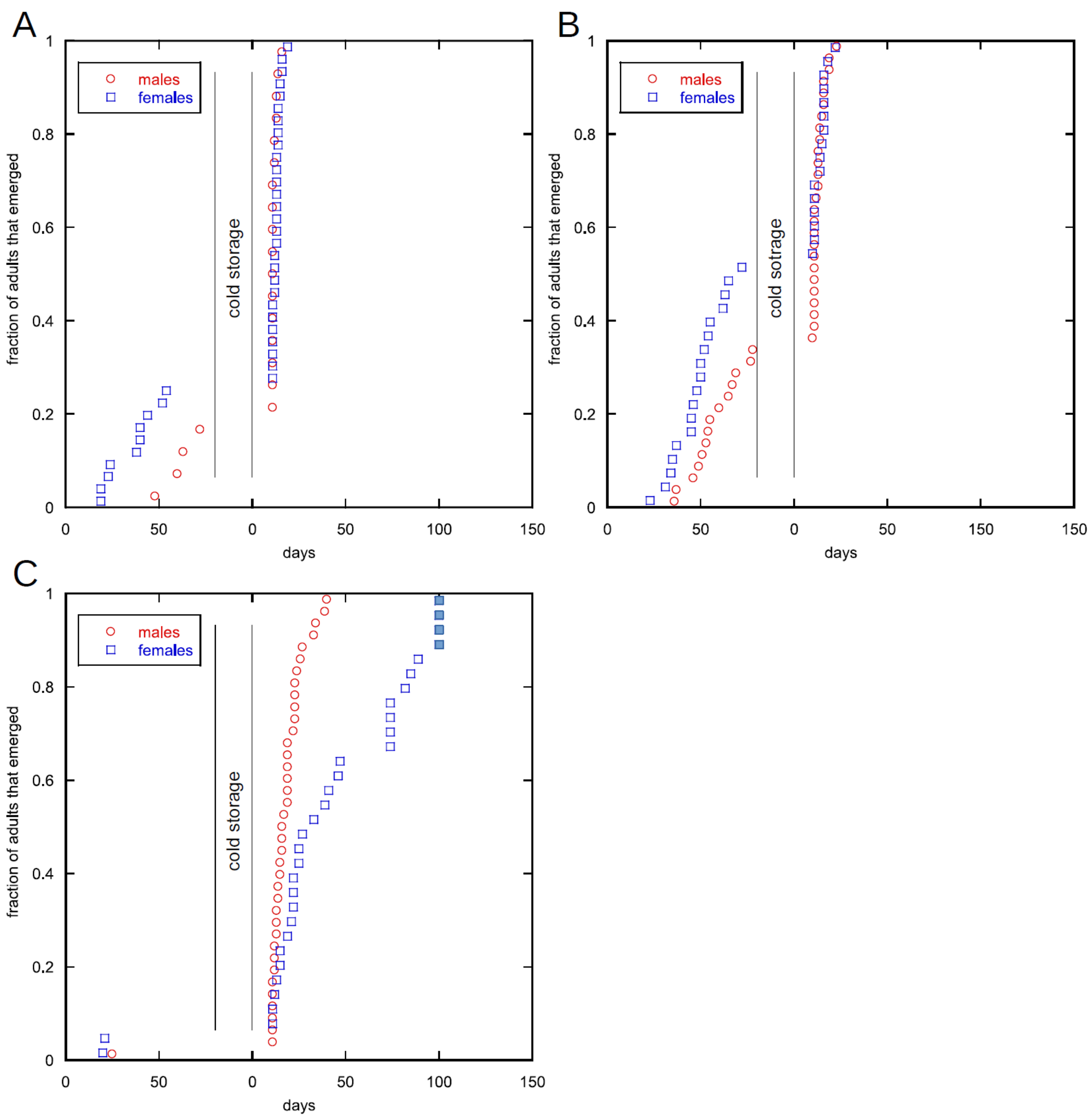


**Figure S8.** Patterns of adult emergence in backcrosses of second-generation *hospiton* x *machaon* hybrids to *P. machaon* determined by presence versus absence of the Z chromosome from *P. hospiton*. The hybrid parents in all three panels were siblings obtained from the same cross between a (male *hospiton* x female *machaon*) female hybrid and a *machaon* male. Larvae from all crosses were raised synchronously under long-nights conditions. (A), (B) Progenies of crosses between a female hybrid and a male *P. machaon* (*P. machaon* parents in the two crosses were siblings). (C) Progeny of crosses between male hybrids and a female *P. machaon* (data from three separate crosses that used *P. machaon* siblings were pooled). Four female pupae that had failed to develop at the end of the experiment (filled symbols) may have been perpetual nymphs.

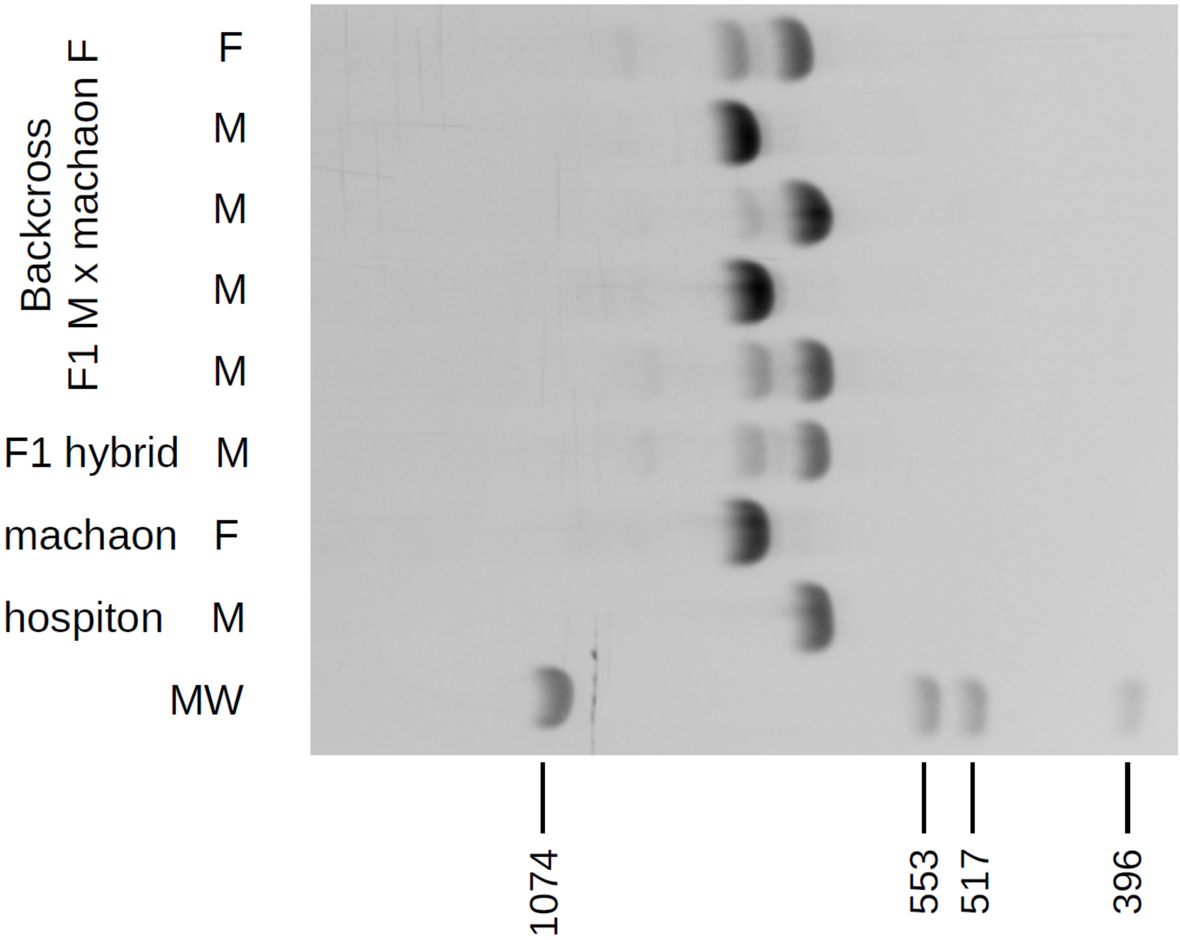


**Figure S9.** PCR amplification of ITS2 DNA from a F1 *hospiton* x *machaon* hybrid and from five individuals obtained by backcrossing a male F1 hybrid with a *machaon* female. MW, molecular weight markers (base pairs); M, male; F, female. Predicted lengths of PCR products of hospiton and machaon are 660 and *ca* 728 bp, respectively.

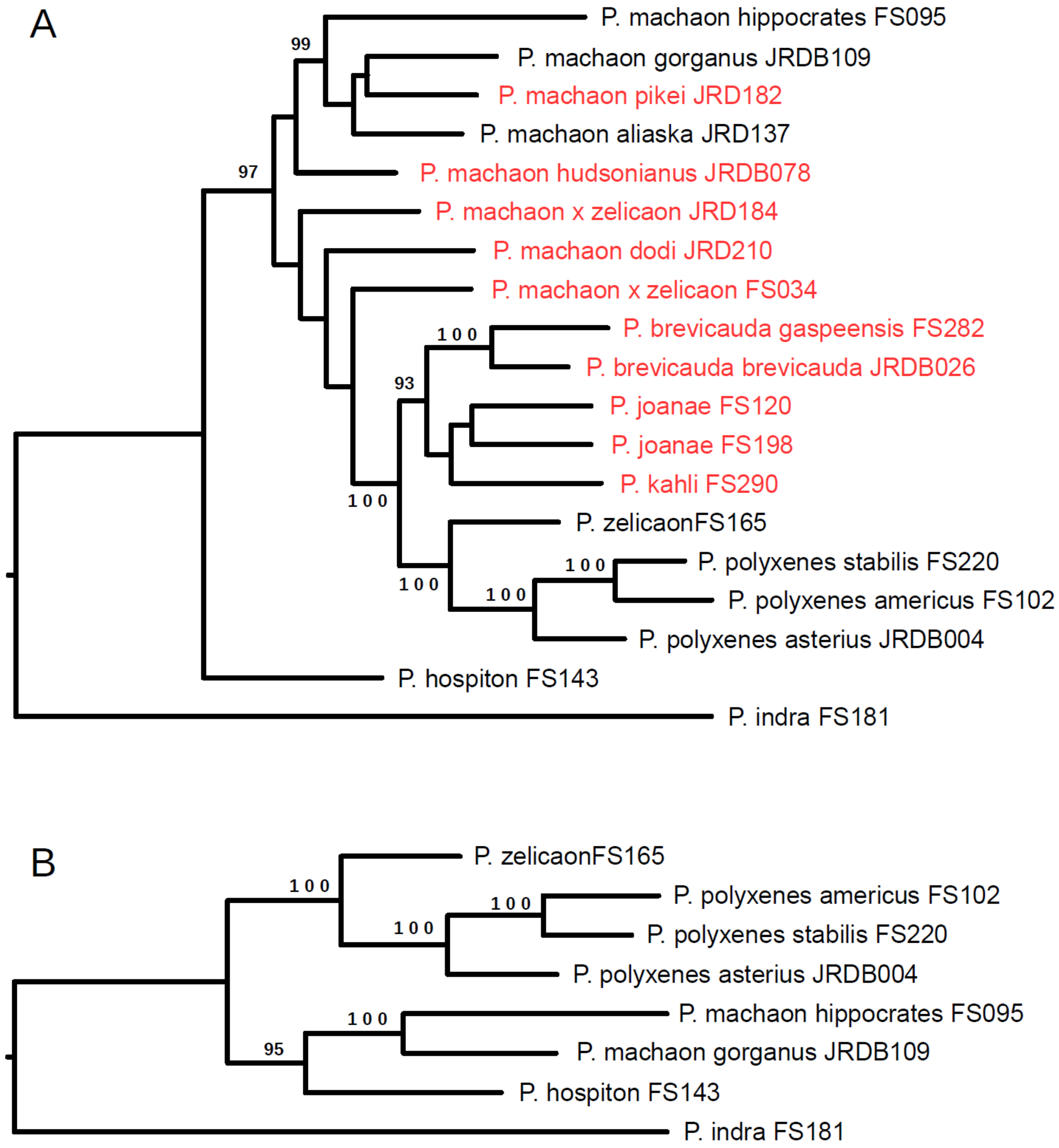


**Figure S10.** Consequences of including Nearctic individuals of hybrid origin on the phylogenetic position of *P.* hospiton. Maximum likelihood phylogenetic trees were generated by IQ-TREE version 2.2 (Nguyen *et al.*, 2015) from concatenated SNP data (Dupuis and Sperling 2020) resulting in an alignment length of 1286541 nucleotides, including flanking sequences. We used the same 19 specimens as in the original publication, with one exception: *zelicaon* individual JRD327, from Alberta, was replaced by specimen FS165, from Hemet, California , which we deemed less likely to have incorporated DNA from *machaon*-related lineages (names in red in panel A correspond to individuals of presumed or proven hybrid origin). The resulting alignments had 2334 (19-specimens set) and 894 (8-specimens set) parsimony informative sites (use of the authors' 'exemplar' dataset – without missing data, 771 SNPs – resulted in identical – panel B – or similar – panel A – topologies, but somewhat lower bootstrap values). Best-fit substitution model was chosen according to BIC criterion (Kalyaanamoorthy *et al.*, 2017). Branch support values (indicated in the figure when above 90 per cent) were obtained with the Ultrafast bootstrap approximation (Hoang *et al.*, 2018).
